# Age-associated inflammatory monocytes are increased in menopausal females and reversed by Hormone Replacement Therapy

**DOI:** 10.1101/2025.03.28.645904

**Authors:** RPH De Maeyer, J Sikora, OV Bracken, B Shih, AF Lloyd, H Peckham, K Hollett, K Abdelhamid, W Cai, M James, PE Pfeffer, M Vukmanovic-Stejic, AN Akbar, ES Chambers

## Abstract

Biological sex is a crucial, but poorly understood variable in age-related susceptibility to infections and chronic inflammation – inflammageing. Monocytes are important immune cells responsible for initiating and resolving inflammatory responses to infection. While changes in monocyte populations result in increased susceptibility to infection, there is limited research on the impact of age and sex on human monocyte phenotype and function. The aim of this work was to dissect the impact of increasing age and biological sex on human monocyte phenotype and function.

Here we show that older females have increased inflammatory intermediate and non-classical monocytes compared to young. These monocyte subsets were the most inflammatory *ex vivo* and their frequency correlated with markers of systemic inflammation. Proteomic analysis of sorted monocyte populations demonstrated that the three human monocyte subsets have largely distinct phenotypes. Additionally, proteomic analysis identified key age-associated protein pathways, including complement cascade and phagocytosis, downregulated in monocytes from older compared to younger individuals. We confirmed the proteomics findings showing that circulating C3 concentrations were reduced with age in females but not males. This decrease in complement in older females resulted in reduced monocyte phagocytosis. Crucially, we demonstrate that in peri/menopausal females, Hormone Replacement Therapy (HRT) reversed this expansion in intermediate monocytes and decreased circulating CRP as compared to age matched controls. Importantly peri/menopausal females on HRT had increased C3 serum concentrations and significantly improved monocyte phagocytosis.

The data presented here indicate the importance of menopause in ageing monocyte phenotype and function. This data highlights the potential use of HRT in restoring monocyte function in females during ageing.

## Introduction

Ageing is a global burden. Older people are living longer with increased morbidity and mortality from infections such as SARS-CoV-2 and influenza^1, 2^, malignancy^3^ and reduced vaccine efficacy^4^ impacting upon health-span. In addition to age, the biological sex of the individual greatly influences which infectious disease poses higher risk of morbidity and mortality. Older males are known to be at increased risk of morbidity and mortality from SARS-CoV-2^1, 5^ and community acquired pneumonia (CAP)^6, 7^, whilst older females have increased morbidity and mortality from influenza^8^ and non-tuberculous mycobacterial pulmonary disease (NTM-PD)^9, 10^. Menopause is a key life-course event in females that leads to increased incidence of inflammatory diseases such as asthma^11, 12^ and increased risk of bacterial urinary tract infections^13^

Inflammageing is a phenomenon whereby circulating inflammatory mediators such as C Reactive protein (CRP), IL-6 and IL-8^14^ increase during ageing. This process underlies age-associated pathologies including frailty^15^, neurodegeneration^16^ and mortality^17, 18, 19^, and is linked to poor vaccine efficacy^20, 21^. Furthermore, monocyte-derived inflammation negatively impacts antigen-specific immunity in human skin^22, 23^. Multiple factors contribute to the inflammageing phenomena however, mononuclear phagocytes and in particular monocytes can contribute to inflammatory milieu through Toll-like receptor (TLR) binding to Lipopolysaccharide (LPS) leaked into the circulation through increased gut permeability^24^ and failure to resolve inflammation due to defects in efferocytosis^25^.

One of the key immune cells involved in coordinating immune responses to pathogens is the monocyte^24, 26, 27^. Monocytes are circulating mononuclear phagocytes historically presumed to be solely precursor cells of tissue macrophages and dendritic cells. However, increasingly it has become apparent that monocytes are important innate immune cells in their own right, and have important effector functions such as early pathogen recognition and initiation of inflammatory responses and phagocytosis^28^. Human monocytes can be separated into three populations based on their cell-surface expression of CD14 and CD16, with CD14+CD16-classical monocytes, CD14+CD16+ intermediate monocytes and CD14-CD16+ non-classical monocytes^29^. CD16+ monocytes have been shown to be expanded in the peripheral blood of older adults^30, 31^. Highlighting their role in disease, it has previously been shown that older adults are at increased risk of Influenza infection due to an age-related defect in RIG-I signalling in monocytes^27^. Furthermore, defective TNF production from monocytes of older people has also been associated with impaired pneumococcal immunity^24^. Earlier studies using mixed cell populations have yielded conflicting data as to the effect of age on inflammatory cytokine production after stimulation *in vitro*^32, 33, 34^. In addition, there is limited data as to the impact of the combination of age and sex on human monocyte function.

Indeed, the effect of biological sex in combination with age on immunosenescence is less well understood. Previous studies have shown that monocytes from males were more inflammatory *in vitro* as compared to females when stimulated with LPS^35^, these data were not extrapolated by age. To date there has been no study that assesses the impact of increasing age and biological sex on monocyte phenotype and function. Therefore, the aim of this paper is to dissect the impact of increasing age and biological sex on human monocyte phenotype and function.

## Materials and Methods

### Study Design

Ethics were approved by the Queen Mary Ethics of Research Committee reference number QMERC23.059 and by the University of Oxford Medical Sciences Interdivisional Research Ethics Committee (ref: R80191/RE001). Individuals 18 years or older were recruited into the study, and full written informed consent was obtained from each donor. 80mls of peripheral blood was taken from each donor and processed as described below. Individuals were excluded from the study if they had a recent history (≤5 years) of neoplasia, inflammatory disorders which required immunosuppressive medication, were anaemic or had a current pregnancy.

### Whole blood flow cytometry

Whole blood was labelled with the following cell surface antibodies: CD14 (HCD14), CD16 (3G8), CD19 (HIB19), CD20 (2H7), CD56 (HCD56), CCR2, CX3CR1, HLA-DR (L243) and Zombie Green Fixable Viability Kit (Biolegend). Cells were incubated with antibodies for 45 mins at 4°C. After incubation red blood cells were removed using FACs Lysis buffer (BD Biosciences). Samples were subsequently washed twice in PBS and then assessed by flow cytometric analysis on a BD Fortessa or on a Cytek Aurora.

FCS files of monocyte populations were exported using FlowJo Version X (BD Biosciences). Clustering and UMAP analysis of monocytes was performed using the R package ‘CATALYST’^36^. Briefly, fcs files are loaded into R studio, combined with the metadata and prepared as a single cell experiment, which is then subject to clustering based on expression of CD14, CD16, CCR2, HLA-DR, CD86, SLAN, CLA and CX3CR1. UMAPs were generated based on 4000 events from each sample.

### Monocyte isolation

Peripheral blood mononuclear cells were isolated by density centrifugation using Ficoll-Paque (Amersham Biosciences). Monocytes were isolated by negative selection according to the manufacturer’s instructions (Miltenyi Biotec).

### Monocyte Cell Sorting

Negatively isolated monocytes were labelled with CD14 A700 (HCD14), CD16 BV786 (3G8), CD19 FITC (HIB19), CD20 FITC (2H7), CD56 FITC (HCD56) and HLA-DR BV510 (L243) for 45 mins at 4°C. Cells were washed twice with PBS. Cells were sorted using the BD FACSAria™ III Cell Sorter (BD Biosciences). Monocytes were identified as being FITC negative and HLA-DR positive, within this population three monocyte populations were sorted classical (CD14^+^CD16^-^), intermediate (CD14^+^CD16^+^) and non-classical (CD14^-^CD16^+^) (Supplementary Figure 1). Sorted monocytes were either cultured *in vitro* or collected for proteomics analysis as described below.

### Monocyte culture

Negatively isolated or sorted monocytes were cultured in RPMI supplemented with 10% FCS, 2mM glutamine and 100U/ml penicillin/streptomycin. Cells were cultured at 5% CO_2_ for 24 hours. Supernatants were collected for cytokine analysis.

### Phagocytosis assay

100 µl whole blood obtained from K2EDTA vacutainers was added to pre-plated antibody cocktail containing CD14 APC-Fire810 (63D3), CD16 eFluor450 (CB16), HLA-DR BUV563 (L243), CD115 BV711 (9-4D2-1E4), CD10 BV750 (HI10a) and Zombie-NIR live/dead dye. Samples were challenged with 12.5 µl pHrodo red-labelled *Escherichia coli* (*E.coli*) BioParticles (ThermoFisher, P35361) or PBS control and left for 30 minutes at 37°C. 500 µl Fix/Lyse (eBioscience) solution was added to stop phagocytosis and lyse red blood cells. Samples were washed and acquired on a Sony ID7000 Spectral Analyzer.

### Supernatant cytokine assessment

Cytokine concentrations in culture supernatants were assessed by cytometric bead array according to the manufacturer’s protocol. Samples were analyzed using a Novocyte flow cytometer (Agilent).

### Proteomics

Sorted monocytes were collected into eppendorfs containing RPMI with 1% FCS. Subsequently the pellets were washed in cold PBS and then pellets were snap frozen on dry and ice and stored at - 80°C. Samples were lysed in 400 µl lysis buffer (5 % SDS, 10 mM TCEP, 50 mM TEAB in highly pure H_2_O) and shaken at RT for 5 min at 1000 rpm, followed by boiling at 95 ◦C for 5 min at 500 rpm. Samples were then shaken again at RT for 5 min at 1000 rpm and sonicated for 15 cycles of 30 s on/ 30 s off with a BioRuptor (Diagenode) before Benzonase was added to each sample and incubated at 37 ◦C for 15 min to digest DNA. Next, samples were alkylated with 20 mM iodoacetamide for 1 h at 22 ◦C. Protein concentration was determined using EZQ protein quantitation kit (Invitrogen) following the manufacturer instructions. Protein isolation and clean up was performed using S-TRAP™ (Protifi) columns before digestion with trypsin at 1:20 ratio (enzyme:protein) for 2 hours at 47 ◦C and digested peptides were eluted from S-TRAP™ columns using 50 mM ammonium bicarbonate, followed by 0.2 % aqueous formic acid and 50 % aqueous acetonitrile containing 0.2 % formic acid and dried overnight. Peptides were resuspended in 1% formic acid and injected onto a nanoscale C18 reverse-phase chromatography system (UltiMate 3000 RSLC nano, Thermo Scientific) and electrosprayed into an Orbitrap™ Exploris 480 Mass Spectrometer (Thermo Fisher).

Raw files are processed through Spectronaut to identify and quantify proteins, which is then run through Perseus which converts the intensity values of each protein into copy numbers. Sample showing zeros in more than 80% of the proteins were removed, and proteins with >0 count in 5 or more samples were retained. Generalised Mass Spectrum (GMS) missing peaks imputation Lasso algorithm from GMSimpute (v0.0.1.0)^37^ was used to impute missing data. Statistical analysis was performed on the imputed dataset using limma (v3.60.5)^38^. Contrast fit were made, treating cell state (CD14+, CD14+CD16+, CD16+) as ordinal variable and age groups as categorical variable. Following lmFit and eBayes, TopTable (age group; adj.p < 0.05) and TopTreat (cell state; adj.p <0.05 and log fold change, logFC > 1.5) with Benjamini-Hochberg correction.

Subsequently, protein expression was determined the package MS-DAP^39^. Spectronaut files were combined with metadata and mapped to human isoform fasta files. Filters were applied so that peptides were quantified in at least 10 replicates prior to normalisation. MS-DAP recommends a normalisation algorithm of ‘vsn’ and ‘modebetween_protein’, which was applied to this analysis. The rollup algorithm from peptide to protein was maxlfq. The differential expression algorithm for the different contrasts was msqrob, with a q value threshold of 0.05. Significance of log2 fold changes was calculated using bootstrap analysis in MS-DAP. Pathway analysis was performed using the clusterProfiler package^40^. All data was visualised in R studio.

### Serum Protein analysis

The serum levels of CRP, C3, and C4 were measured using sandwich ELISA kits following the manufacturer’s instructions. Dilutions were made as needed to ensure that the readings fell within the ranges of the assays. The concentration of CRP was measured using the DuoSet Human C-Reactive Protein/CRP DuoSet ELISA kit (Cat. No DY1707) from R&D Systems (Minneapolis, MN, USA). The concentrations of C3 (Cat. No ab108822) and C4 (Cat. No ab108824) were determined using Abcam kits (Abcam™, Cambridge, UK). The sensitivities of these kits were approximately 994 µg/mL for C3 and 0.53 µg/mL for C4. The precision (intra-assay and inter-assay) for all the assays was approximately 10%.

### Statistical analysis

Statistical analysis was performed using Prism version 10.0.0 (GraphPad). Data were assessed for normality, and the subsequent appropriate two-sided statistical test was performed as indicated in the figure legend. For the correlations Pearson’s correlation was performed in Prism.

## Results

### Increased frequency of CD16 expressing monocytes in older adults

To assess the phenotype of monocytes in young (<40 years) and older adults (≥65 years), whole peripheral blood was labelled for flow cytometric analysis and monocytes were identified from live single cells as being lineage negative (CD3, CD19, CD20 and CD56) and HLA-DR+ (Supplementary Figure 1). Monocyte subsets were identified based on the expression of CD14 and CD16; classical monocytes as CD14+CD16-(CD14+), intermediates as CD14+CD16+ and non-classicals as CD14-CD16+ (CD16+). Older adults (mean age 73.1; 17 females and 14 males) had a significant increase in intermediate and non-classical monocytes compared to younger adults (mean age 29.2; 26 females and 17 males) (Figure 1A and B). This was further examined using alternative markers for monocytes CCR2 and CX3CR1 and in line with earlier work from Patel et al^29^, it was confirmed that there was a significant increase in the frequency of non-classical monocytes (CCR2-CX3CR1+) in older adults as compared to young (Figure 1C and D). As expected, the expression of CCR2 and CX3CR1 correlated with CD14 and CD16 (Figure 1E), demonstrating that non-classical and intermediate monocytes were increased in older adults as compared to young. Unsupervised analysis of the data was performed where cells were clustered using the FlowSOM algorithm, which identified four unique clusters: CLA+CD14+, CLA-CD14+, CD14+CD16+ and CD16+. These clusters were visualised via a UMAP (Figure 1F and G). It was observed that there was no difference in the surface expression of proteins studied (CLA, CD14, CCR2, CD86, CD16, SLAN, CX3CR1 and HLA-DR) according to age (Supplementary Figure 2). Suggesting the age-related change in monocyte subset quantification is likely due to increased monocyte differentiation. Collectively this data shows that the proportion of monocyte subsets significantly changes in ageing.

**Figure 1:**
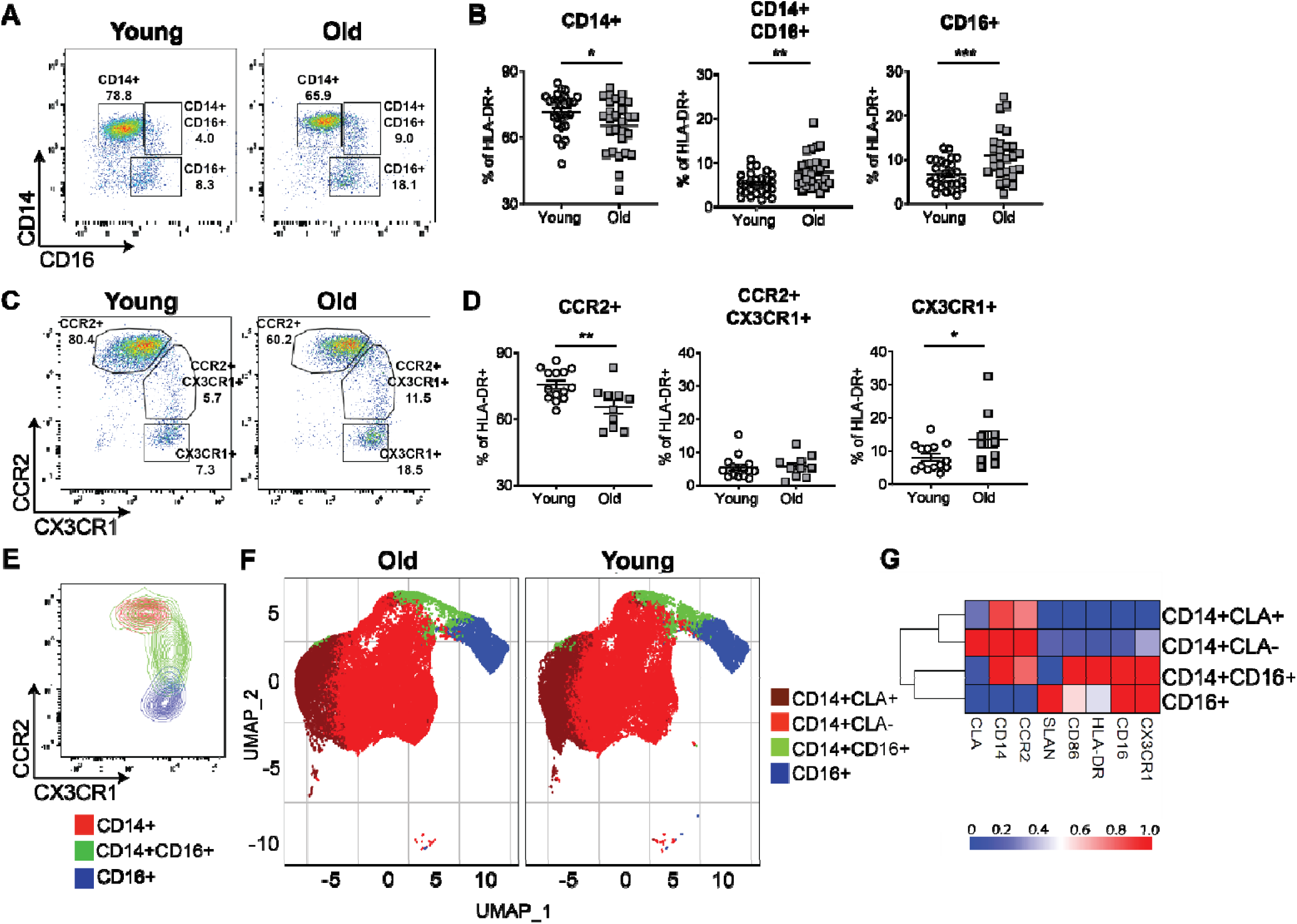
Increased frequency of non-classical and intermediate monocytes in older adults. Whole blood was assessed by flow cytometry and monocytes were identified as being Lineage negative HLA-DR+. **A,** representative flow plots and **B,** cumulative data showing the frequency of CD14+ (classical), CD14+CD16+ (DP; intermediate) and CD16+ (non-classical) and **C,** representative flow plots and **D,** cumulative data showing the frequency of CCR2+ (classical), CX3CR1+CCR2+ (DP; intermediate) and CX3CR1+ (non-classical) monocytes monocytes in the peripheral blood of young (white; <40yrs) and old (grey; ≥65yrs) donors. **E,** a representative image showing an overlay plot of CD14+ (red), CD14+Cd16+ (green) and CD16+ (blue) monocytes according to expression of CCR2 and CX3CR1. Bioinformatic analysis was performed on monocytes and analysed based upon the monocyte markers SLAN, CLA, CCR2, CD14, CD16, CD86, HLA-DR and CX3CR1. **F**, UMAP analysis of monocytes from young and old monocytes and **G,** a heatmap of the marker expression in the four clusters. B, D, and H, assessed by t-test. * = p<0.05; ** = p<0.01; *** = p<0.001.

### CD16 expressing monocytes are more inflammatory and correlate with inflammageing

Inflammageing is defined as low grade chronic inflammation characterised by elevated circulating inflammatory proteins such as C Reactive Protein (CRP)^14, 41^, which is observed during ageing. As expected, we observed a significant increase in serum CRP in older adults as compared to young (Figure 2A). To determine if the increased proportion of CD16 expressing monocytes in older adults corresponds to increased marker of inflammageing, CRP was correlated with frequency of monocyte populations. It was observed that there was a significant positive correlation between serum CRP concentrations and the frequency of CD16+ non-classical monocytes (Figure 2B), with the individuals who had the most CD16+ non-classical monocytes having the highest serum CRP concentration. Furthermore, a significant negative correlation was observed between CRP serum concentrations and the frequency of CD14+ monocytes.

**Figure 2:**
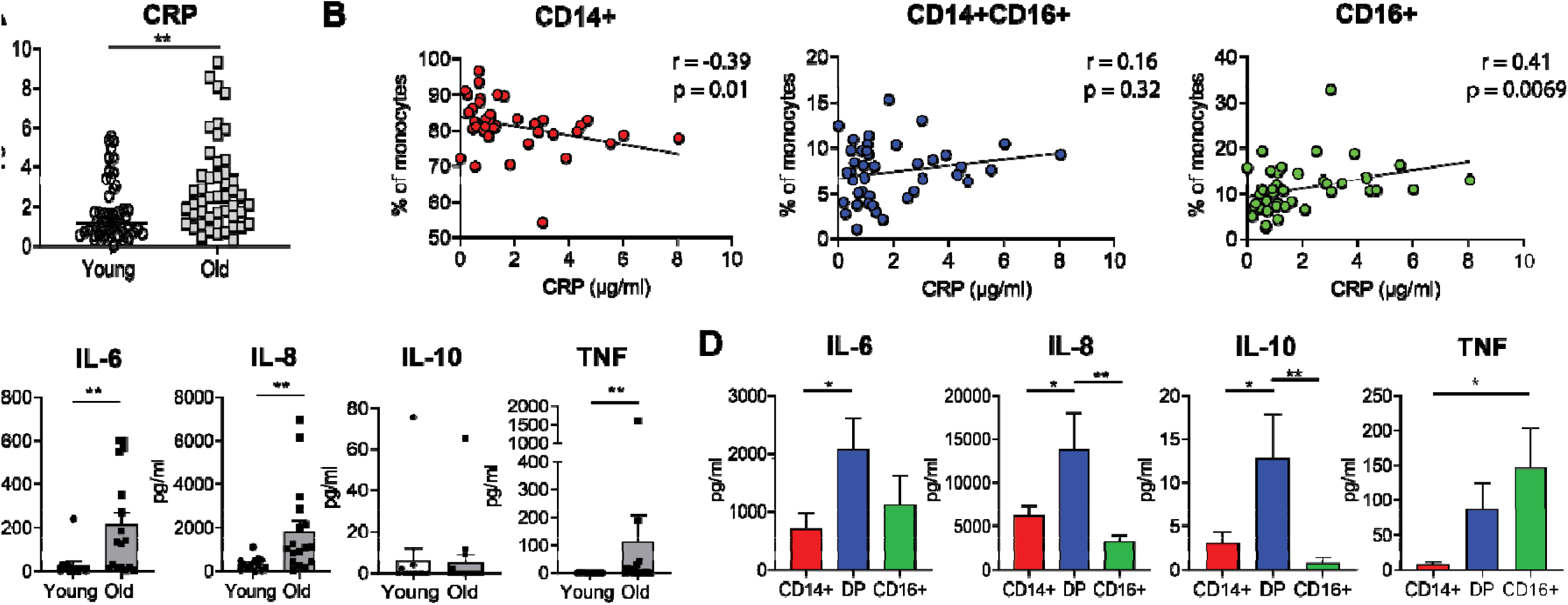
CD16+ monocytes are more inflammatory and correlate with C Reactive Protein a marker of inflammageing. Serum C Reactive Protein (CRP) concentrations were assessed in biobanked serum samples by ELISA **A,** Serum CRP concentrations in young (white) and old (grey) donors and **B,** Serum CRP correlated with frequency of monocyte populations in the peripheral blood of the same individual. **C,** Monocytes were isolated from the peripheral blood and cultured (unstimulated) for 24 hours, supernatants were collected and assessed by cytometric bead array cumulative cytokine production from young (white) and old (grey) donors. **D,** Cumulative data of monocytes sorted into three populations classical (CD14+), intermediate (CD14+CD16+) and non-classical (CD16+) and cultured (unstimulated) for 24 hours. Cytokine production after 24 hours from CD14+ (red), CD14+CD16+ (green) and CD16+ (blue). A, assessed by Kruskal-Wallis test with Dunn’s multiple comparisons test and B, assessed by Pearson’s correlation test. C and D, assessed by Mann Whitney. E, assessed by One-way ANOVA with Tukey’s multiple comparison test. * = p<0.05; ** = p<0.01.

Alongside higher levels of CRP, inflammageing is also associated with increased levels of IL-6, IL-1 and TNF^14, 41^. To investigate the effect of inflammatory cytokine expression, monocytes were isolated from young and old donors and cultured for 24 hours (unstimulated). Cytokine expression in the supernatant was assessed by cytometric bead array and it was observed that monocytes from older individuals produced significantly more IL-6, IL-8 and TNF, but not IL-10 than monocytes from younger individuals (Figure 2C). To determine which monocyte subsets from older adults, make more inflammatory cytokines, monocytes subsets were sorted by FACs and cultured at the same cell concentration, separately for 24 hours. CD16+CD14+ monocytes expressed significantly more IL-8, IL-8 and IL-10 as compared to the other monocyte subsets (Figure 2D). However, CD16+ non-classical monocytes secreted significantly more TNF as compared to CD14+ monocytes (Figure 2D). No significant difference in cytokine production by the individual monocyte populations was observed with age (Supplementary Figure 3). This indicates that the inflammageing process is linked to increased frequency of intermediate and non-classical monocytes found in older individuals are implicated in the inflammageing, as opposed to increased cytokine production by individual monocyte cells with age.

### The proteome of the three different monocyte populations is significantly different indicating distinct functionality

Due to the effects of age on monocyte phenotype and function and the significant differences observed proteomic analysis was performed on monocytes from young and old donors and to enable deeper phenotyping of the effect of age on monocyte subsets, we performed proteomic analysis of the individual monocyte populations (CD14+ classical; CD14+CD16+ intermediate; CD16+ non-classical) from 8 young and 8 old donors. Protein expression is the main mechanistic indictor of function and can therefore provide detailed insight into functional differences between monocyte populations. The CD14 and CD16 expression levels from the proteomic data was utilized to confirm that their levels correspond to the CD14 and CD16 labelling from FACs. PCA analysis showed distinct proteomes of the three monocyte populations (Supplementary Figure 4A) and protein expression of CD14 and CD16 was observed in the expected populations (Supplementary Figure 4B). The number of proteins quantified between the three different monocyte populations was also not statistically different (Supplementary Figure 4C).

The top significantly differentially expressed proteins according to monocyte population (transitioning from CD14+, CD14+CD16+, through to CD16+) were assessed and it was observed that there were significant proteomic differences between the three different monocyte populations (Figure 3A; Supplementary Table 1). The package MS-DAP was used to quantify protein intensity in monocyte populations. Differential expression analysis using the algorithm MSqRob was employed to quantify proteomic differences between the subsets. The differentially expressed proteins in each comparison were visualised using a Venn diagram (Figure 3B) revealing 416 uniquely expressed proteins in CD14+, as compared to CD14+CD16+ and 231 uniquely expressed proteins in CD14+CD16+ as compared to CD16+ and 213 uniquely expressed proteins in CD16+ as compared to CD14+ (Figure 3B). A full list of differentially regulated proteins identified in the heatmap can be found in Supplementary Table 2. To visualise the differences between these contrasts we used volcano plots to view the top 25 upregulated and downregulated proteins, further showing the unique signatures between the different subsets (Figure 3C, D and E). Given, the differences in monocyte subset proportions with age and the distinct proteomic differences of the three different monocyte populations this data support the hypothesis that there will be overall alteration in monocyte functionality with increasing age.

**Figure 3:**
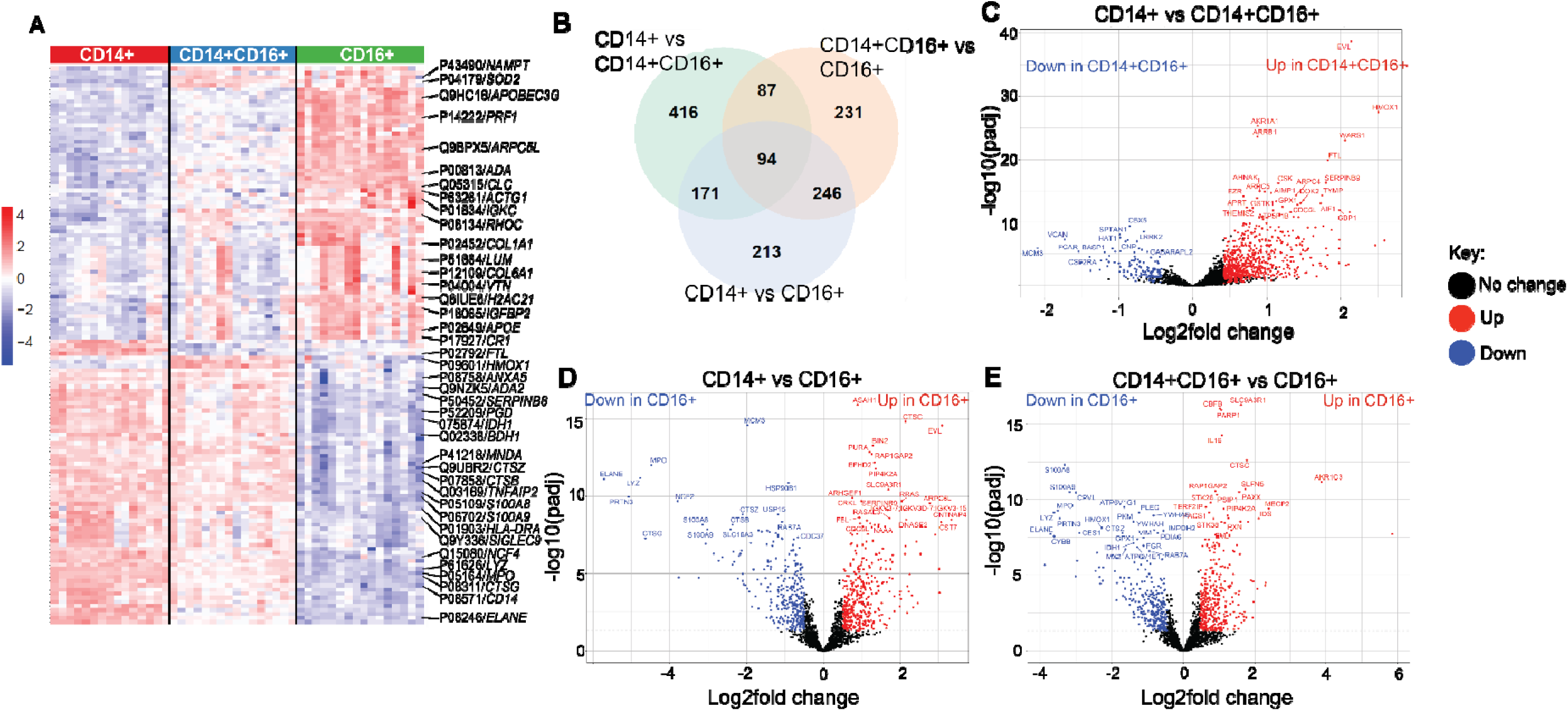
Proteomic composition of the three different monocyte populations is significantly different. Monocytes were isolated from the peripheral blood and sorted *ex vivo* into three populations classical (CD14+; red), intermediate (CD14+CD16+; blue) and non-classical (CD16+; green), proteomics analysis was performed on the ex vivo cell pellets. **A,** heatmap of significantly differentiated protein copy numbers including highlighted proteins (with gene names). Proteomics as analysed by the package MSDAP to assess over protein expression between the three different monocyte subsets **B,** Venn diagram showing differentially expressed proteins in the three monocyte subsets and significantly differentially expressed proteins in **C,** CD14+ vs CD14+CD16+ and **D,** CD14+ vs CD16+ and **E,** CD14+CD16+ vs CD16+.

### The proteome of monocytes is significantly altered with age

There was a subtle but striking separation of the monocyte subsets on PCA analysis by age (Supplementary 4D). Subsequently, the proteomic dataset was analysed to determine which proteins were significantly altered with age in all monocyte populations. It was determined that there were 369 proteins whose expression was significantly altered with age were identified, the vast majority of these were downregulated in old compared to young (Figure 4A; Supplementary Table 2). Proteins significantly increased in older monocytes included those involved in endocytosis (HGS, EEA1), intracellular signalling (MAP2K4, PPIP5K2), ubiquitination (WDR48, CUL4A), intracellular organelle motility (KTN1), chromatin-remodelling (SMARCB1) and metabolism (CLPB) (Figure 4B; full list of significantly changes proteins can be seen in Supplementary Table 3). Proteins which were significantly decreased in older monocytes as compared to young, included complement proteins (C3, C4 and C1QBP), proteins involved in migration (CD44; TUBB4B, TUBB, ITGAL), metabolism (GAPDH) and scavenger receptors (CD68 and CD48) (Figure 4C).

**Figure 4:**
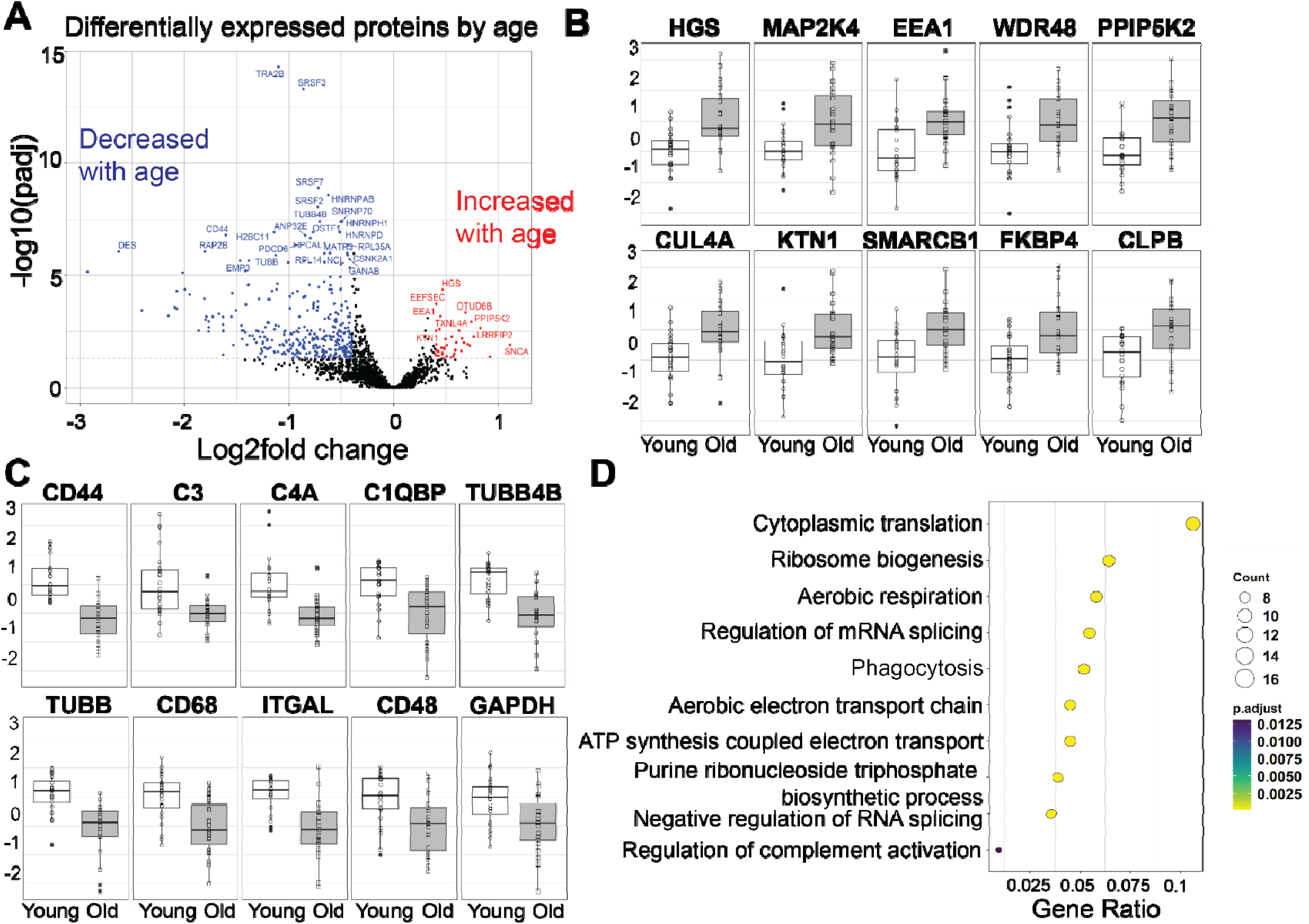
Older monocytes have significantly altered proteome including reduced phagocytic proteins. Monocytes were isolated from the peripheral blood and sorted into three populations classical (CD14+), intermediate (CD14+CD16+) and non-classical (CD16+), proteomics analysis was performed on the ex vivo cell pellets. **A,** Volcano plot showing the differentially regulated proteins in young and old monocytes **B,** Significantly increased proteins in old monocytes as compared to young and **C,** significantly decreased proteins in old monocytes as compared to young. **D,** pathway analysis of pathways significantly decreased in old monocytes as compared to young.

To dissect which pathways were specifically decreased with age, pathways analysis using GO terms was performed. There was a significant downregulation in metabolic pathways such as aerobic respiration, purine ribonucleoside triphosphate biosynthetic process and phagocytosis associated pathways such as phagocytosis and regulation of the complement cascade (Figure 4D). Indeed, when phagocytosis proteins were assessed individually, they were all significantly downregulated in older monocyte populations as compared to young (Supplementary Figure 5). Collectively this proteomic data proposes suggest that older monocytes have significantly altered proteome with reduced metabolic and phagocytic pathways.

### Monocyte phenotype and phagocytic function is significantly altered in older females

As phagocytosis and complement pathways were significantly reduced in older monocytes (Figure 4D), the expression of complement proteins in serum was investigated to determine if this decrease was also seen systemically. Biobanked serum from young and old donors were assessed for C3 and C4 concentrations by ELISA. There was significantly less C3 but not C4 in older donors as compared to young (Figure 5A). Donors recruited in this study had a female bias and as previous data suggests there may be differences in function of monocytes according to biological sex^35^. Thus, we separated the complement data according to biological sex and observed that older females had a significant reduction in C3 and C4 as compared to young females (Figure 5B). This was not observed in males. In addition, there was a significant inverse correlation between serum C3 concentration and age in females but not in males (Supplementary Figure 6).

**Figure 5:**
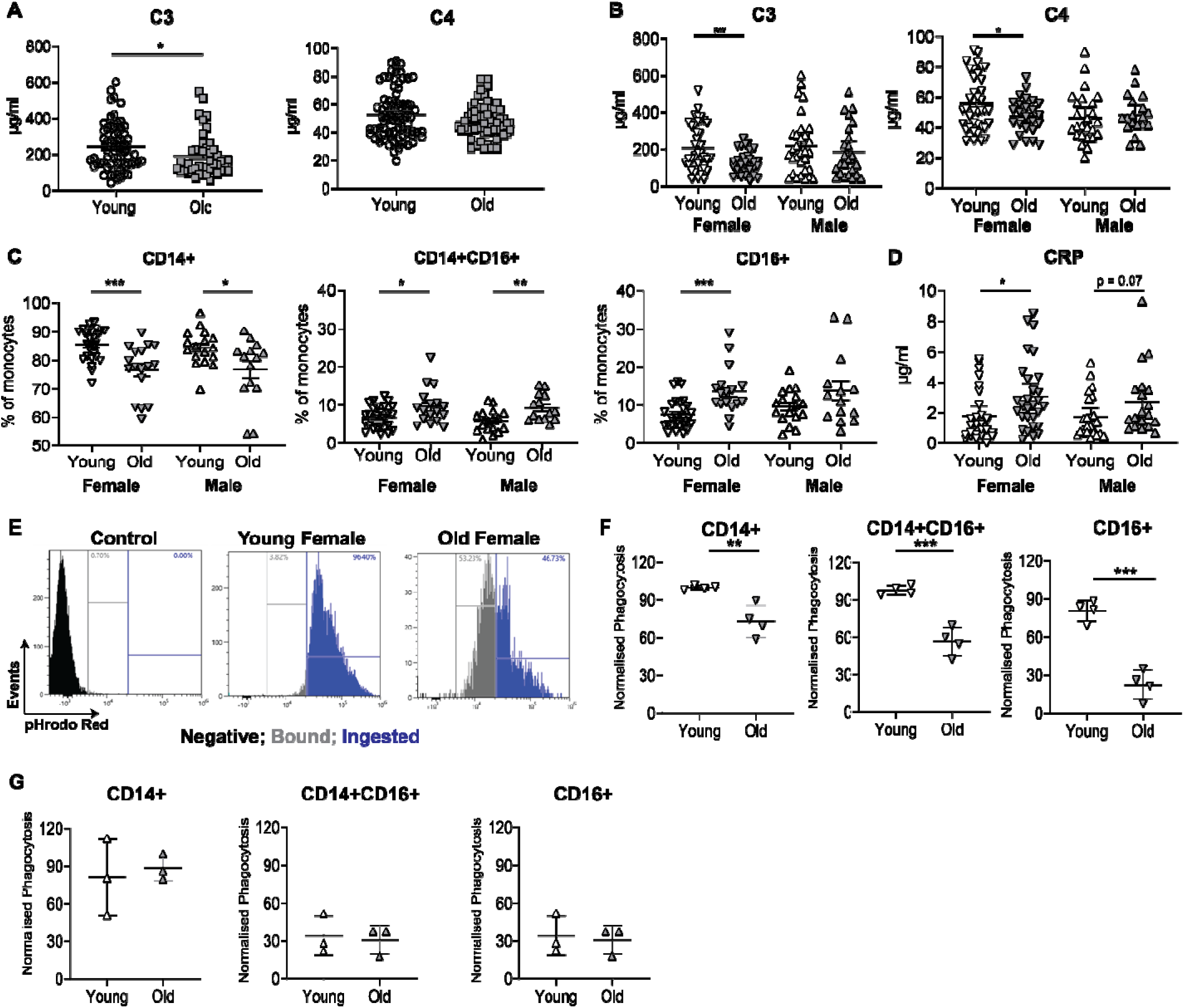
Reduced complement and phagocytosis in older females. Serum samples were assessed for C3 and C4 concentrations by ELISA **A,** serum C3 and C4 concentrations in young (white) and old (grey) donors. **B,** Serum C3 and C4 and **C,** proportion of monocytes and **D,** serum CRP concentrations split according to age (young,<40yrs,white; old, ≥65 yrs, grey) and sex (males upwards triangles; females downwards triangles). Whole blood phagocytosis assay was performed **E,** representative dot plot with control gate (black), bound gate (grey) and ingested gate (blue) and cumulative data in CD14+, CD14+CD16+ and CD16+ in **F,** female individuals and **G,** male individuals. B-D and F assessed by unpaired t test. * = p<0.05; ** = p<0.01; *** p<0.001.

Given the striking difference in serum C3 concentrations in females with age, we analysed monocyte proportions according to age and sex to assess covariance. We observed a decrease in CD14+monocytes in both males and females with age, but the difference was more significant in females (Figure 5C). There was a significant increase in intermediate monocytes with age in both sexes, however, an increase in CD16+ monocytes were only observed in older females (Figure 5C). Serum CRP was found to be significantly increased in females with age, whilst a trend was observed in males (Figure 5D). When the correlation between CRP and monocyte proportions was assessed CD16+ monocytes positively and CD14+ monocytes negatively correlated in females but not males (Supplementary Figure 7), showing similar significance in females as observed in Figure 2B.

To determine if the change in monocyte proportions and circulating complement in older females corresponded to functional deficiency, a whole blood phagocytosis assay was performed. Whole blood was incubated with fluorescently labelled *E. coli* and phagocytosis assessed thirty minutes after the incubation. It was observed that there was significantly less phagocytosis in older females as compared to young (Figure 5E and F). There was no significant difference in male donors according to age (Figure 5G), demonstrating that this was a female age-associated effect.

Collectively this data shows that there is an age-associated defect in monocytes from females with age. This results in a significant reduction in monocyte phagocytosis in females but not males, due to reduced circulating C3.

### HRT alters monocyte phenotype and improves function in peri-/menopausal females

One key life-course event which occurs in females but not males is the menopause. Menopause is associated with the reduction and eventual loss in female sex hormones from the ovaries. Many women counteract this loss of female sex hormones via the use of Hormone Replacement Therapy (HRT)^42^. To assess whether the loss of sex hormones as a result of menopause, leads to the change in monocyte phenotype in older females, peri/menopausal women on HRT (HRT 51.7yrs; age- and sex-matched controls 51.0yrs) were recruited and assessed for monocyte phenotype by flow cytometry. Females who were on HRT had significantly less CD14+CD16+ intermediate monocytes and CD14+ classical monocytes as compared to age matched control (Figure 6A) although no significant different was observed in CD16+ non-classical monocyte population. Menopause and the loss of sex hormones seems to be a key driver in the change in monocyte proportions, as the older women taking HRT have a similar frequency of CD14+CD16+ monocytes as compared to young females. As we previously identified a correlation between inflammatory monocytes and serum CRP, serum CRP levels were assessed in the two female middle-aged cohorts. A significant decrease in CRP concentrations was observed in those individuals taking HRT as compared to age matched controls (Figure 6B). This raises the question of functionality of monocytes during ageing and how is HRT affecting these functions. The concentration of C3 and C4 was assessed in the serum of peri-/menopausal females with and without HRT. It was observed that there was significantly more circulating C3 in the serum of females on HRT as compared to age matched controls (Figure 6C). Finally, we observed that peri-/menopausal females on HRT had a significantly increased phagocytosis as compared to age matched controls (Figure 6D).

**Figure 6:**
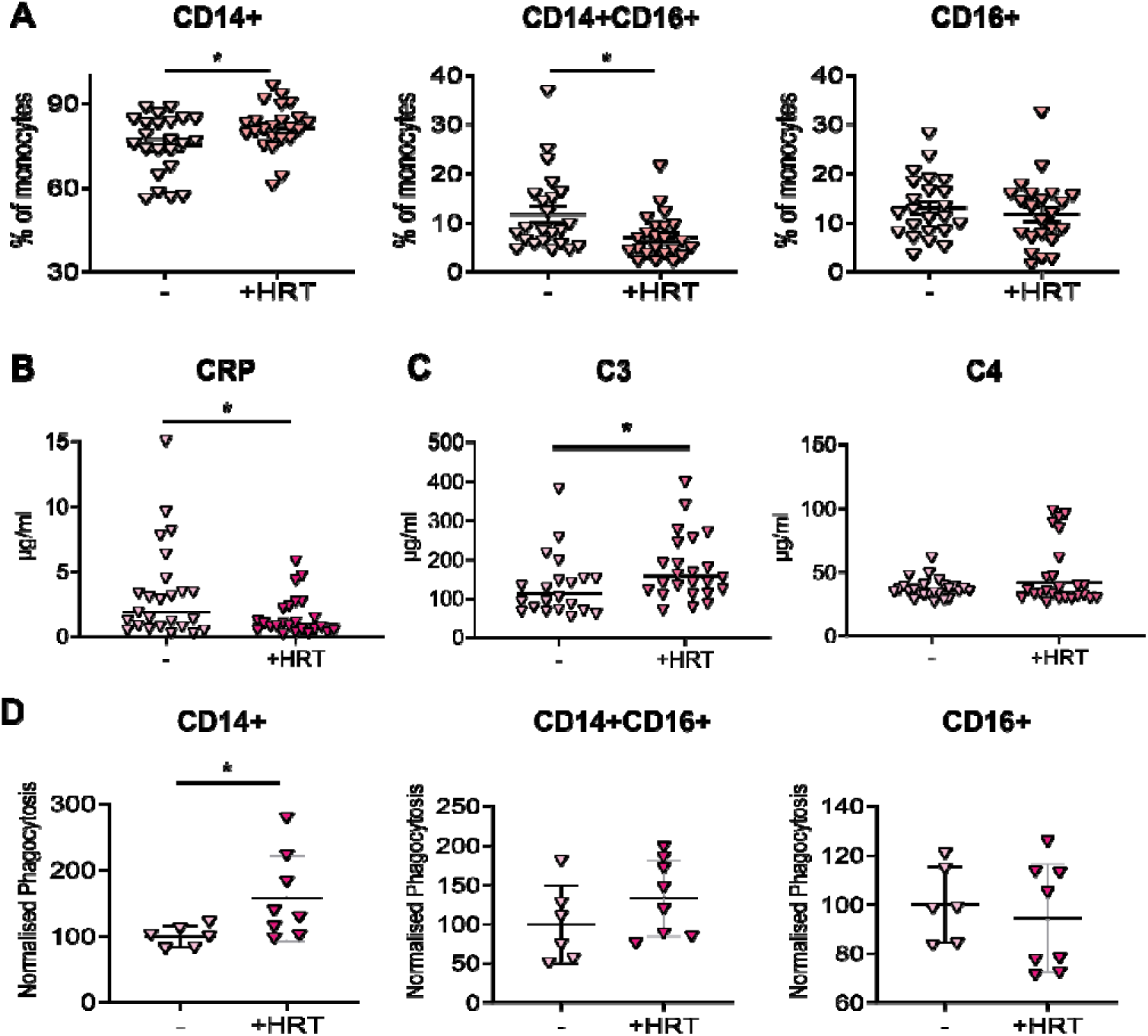
HRT rescues female age-associated intermediate monocyte expansion and phagocytosis activity. Whole blood and serum were collected from peri-/menopausal females who were either taking or not taking Hormone Replacement Therapy (+HRT). **A,** Whole blood was assessed by flow cytometry and monocytes were identified as being Lineage negative HLA-DR+, frequency of CD14+ (classical), CD14+CD16+ (DP; intermediate) and CD16+ (non-classical) monocytes. **B,** CRP serum concentrations measured by ELISA. **C,** Serum C3 and C4 concentrations measured by ELISA. **D,** cumulative whole blood phagocytosis data in CD14+, CD14+CD16+ and CD16+A,-D, assessed by t-test. * = p<0.05

These experiments show that menopause is associated with a reduction in C3 and phagocytosis with an associated increase CD16+ monocytes, which can be reversed by HRT.

## Discussion

This is the first proteomic and functional analysis to assess the impact of age and sex on monocyte phenotype and function. We, like others, observed a relative expansion of intermediate and non-classical monocyte subsets at the expense of classical monocytes with age^30, 43, 44^. We observed a particularly pronounced phenotype in older females, compared to males. We also show that these expanded subsets are more inflammatory when left unchallenged *ex vivo*, and that a shift away from classical to non-classical monocytes correlates with markers of inflammageing such as systemic CRP. Proteomic analysis of monocyte populations identified a significant downward shift in protein expression with age especially in pathways involved with metabolism, complement activation and phagocytosis. These findings were validated by a significant reduction of serum C3 and the reduced phagocytic capacity of monocytes in older females. Finally, HRT use in peri-/menopausal women rescue the female age-associated alterations in monocytes, increasing circulating C3 and enhancing monocyte phagocytosis. These data suggest that intermediate and non-classical monocytes inherently underlie systemic inflammatory features. Thus, their expansion in older people, in particular older women, might be a driver of inflammageing which can be targeted using HRT.

The change to monocyte subset proportions in age has been assessed previously, and our data replicates other datasets^30, 33^ showing older adults have an increase in CD16+ monocytes. Although some studies have not always found these differences, we posit this could be a result of the effect of other variables like biological sex^45^. To our knowledge, only one study assessing monocyte phenotype with age extrapolated the data based upon biological sex^46^. However, we observed that older females have a more significant increase in CD16+ monocytes, a population that has been linked to inflammation and inflammatory disease states^47^. Understanding the effect of biological sex and sex hormones on immunity has become an important and necessary research area. Recently, it has been observed that oestrogen can increase class-switched B cells in females on an XX chromosome background^48^. Whilst testosterone, administered via gender-reaffirming hormone therapy in transgender males can make monocytes more inflammatory with increased cytokine production. This was the first data set to assess the impact of age and biological sex on monocyte phenotype and function^49^. Although we did not observe any difference with sex regarding inflammatory cytokine production further research may be needed to determine at what age testosterone and oestrogen have the biggest influence on inflammatory cytokine production from monocytes. It is clear from this data that the menopause has a dramatic impact upon immunity in females, and as such this is an important area of research to evaluate further.

We show that older monocytes make significantly more inflammatory cytokines compared to younger monocytes when cultured for 24 hours in the absence of stimulation. This increase in inflammatory cytokine production is explained by the increase in the frequency of CD16+ monocytes in older people, rather than the per cell expression of pro-inflammatory cytokines. This is in line with earlier studies which showed that there was no difference in sorted monocyte population inflammatory cytokine production with age and speaks to the importance of correcting imbalanced monocyte populations with age/sex or in disease contexts ^31^ In contrast, we did observe cellular dysfunction in terms of phagocytosis in older females. Previous studies into aged mouse models have observed that macrophages from older mice have reduced phagocytosis^50, 51^. More recently we observed that macrophages in a skin blister model have reduced efferocytosis in older adults^25^. Ours is the first study to look at the impact of age and sex on monocyte phagocytosis, in which we observed reduced phagocytosis in older compared to younger females, with males showing no age-dependent change in phagocytic function. These data provide a hypothesis for why older females are more susceptible to bacterial infections such as NTM and bacterial UTIs.

Crucially, we have shown the importance of HRT in the restoration of monocyte subset proportion and function in older females. HRT has been transformative in improving the lives of menopausal and perimenopausal women by improving the clinical symptoms that some women suffer from during this time^52^. Our study is the first to show the impact that this hormone replacement has on the function of the ageing female immune system. We show that HRT reverses the age-related increase in inflammatory monocyte subsets and restores monocyte phagocytic functionality. Whilst there is much discussion on the benefits and risks of HRT^42, 53, 54, 55^, there is currently little evidence supporting how, at the molecular level at least, these drugs could be crucial in improving the immune health-span of females during ageing. It should serve as a call for more dedicated studies looking at HRT and lowered infection risk in older women.

To conclude, we have shown that being older and female dramatically impacts upon monocyte phenotype and function, with increased inflammatory CD16+ monocytes and reduced phagocytosis contributing to inflammaging and as such decreased health-span. Collectively these data highlight the importance of studying age and sex as biological variables within experiments and provide evidence that HRT could be instrumental in modulating age-dependent inflammation and the complications thereof.

## Acknowledgements

The authors would like to thank all the participants who gave blood for this study. The authors would like to thank Professor Doreen Cantrell and members of the Cantrell group and the Fingerprint proteomics facility at the University of Dundee for proteomics analysis and advice. We would also like to thank Dr Elizabeth Rosser and Professor Coziana Ciurtin for their collaboration on sex differences in immune responses and supervision of Dr Hannah Peckham.

## Author contributions

ESC designed study, performed experiments, analysed data and wrote the first draft. RDM performed experiments, analysed data and wrote the first draft. JS, KA and WC performed experiments. MJ, HP and PEP were involved in sample recruitment and collection. MVS and ANA were involved with early study design. OVB and BS performed the proteomic bioinformatic analysis. AM performed the proteomic analysis. All co-authors read the final version of the manuscript.

## Data availability statement

Proteomics data will be deposited upon acceptance at peer review journal. All other data is available upon request from the corresponding author.

## Supplementary Figures

**Supplementary Figure 1:**
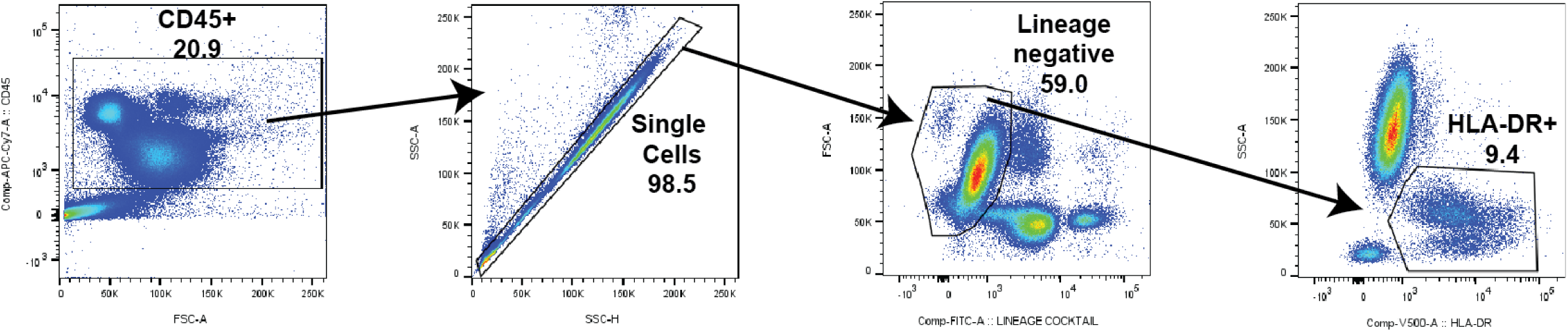
Example gating strategy to identify monocytes in whole blood. Figure shows representative gating strategy to identify monocytes in whole blood was assessed by flow cytometry. Leukocytes were identified as being CD45+, subsequently single cells were identified using SSc-A and SSc-H. Next lineage negative (CD3, CD19, CD20, CD56) cells were identified then HLA-DR+ cells were selected to assess monocyte populations.

**Supplementary Figure 2:**
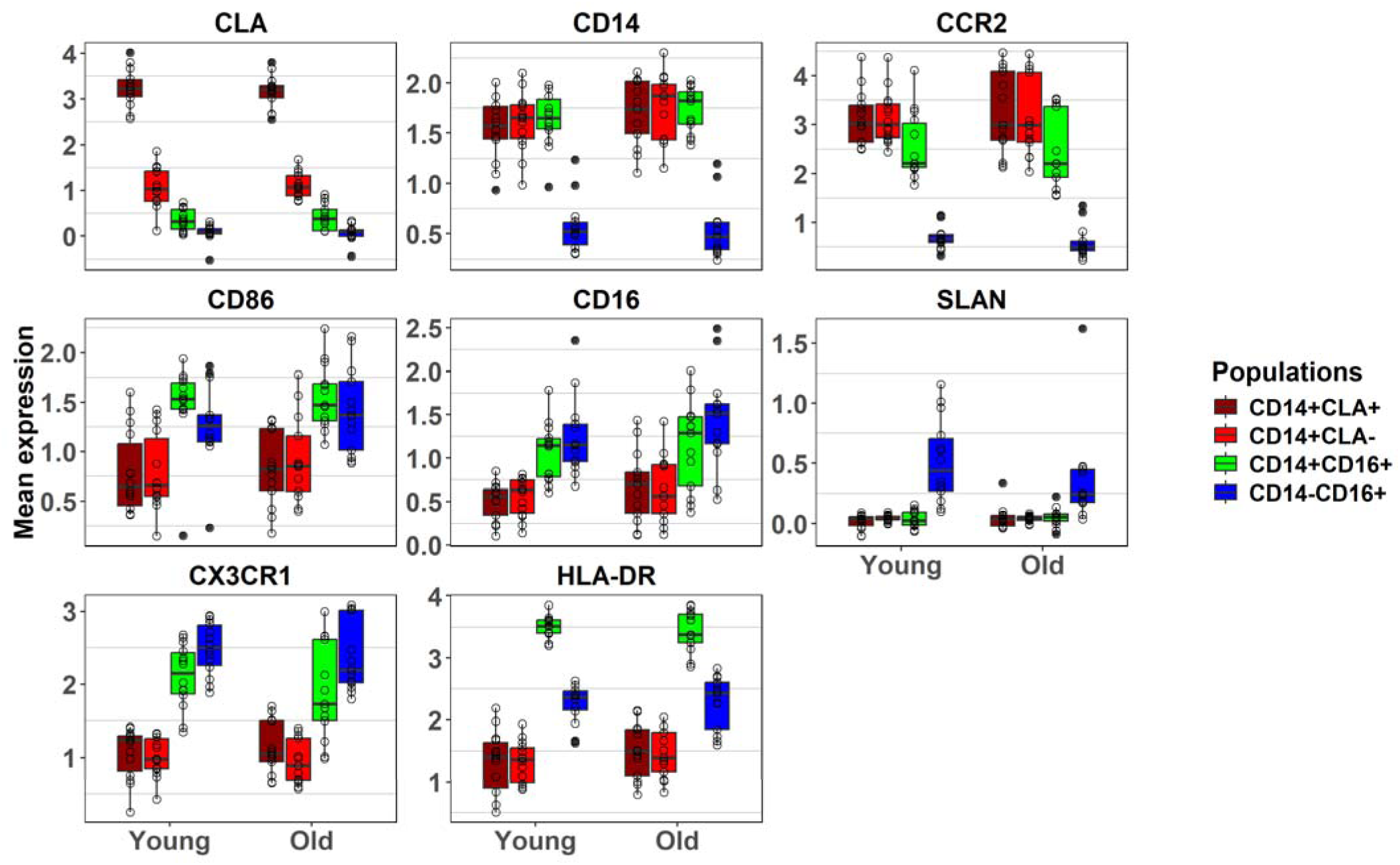
Marker expression in the four different UMAP populations separated according to age. Whole blood was assessed by flow cytometry and monocytes were identified as being Lineage negative HLA-DR+. Bioinformatic analysis was performed on monocytes and analysed based upon the monocyte markers SLAN, CLA, CCR2, CD14, CD16, CD86, HLA-DR and CX3CR1. Marker expression in the four groups identified in Figure 1E are split according to age

**Supplementary Figure 3:**
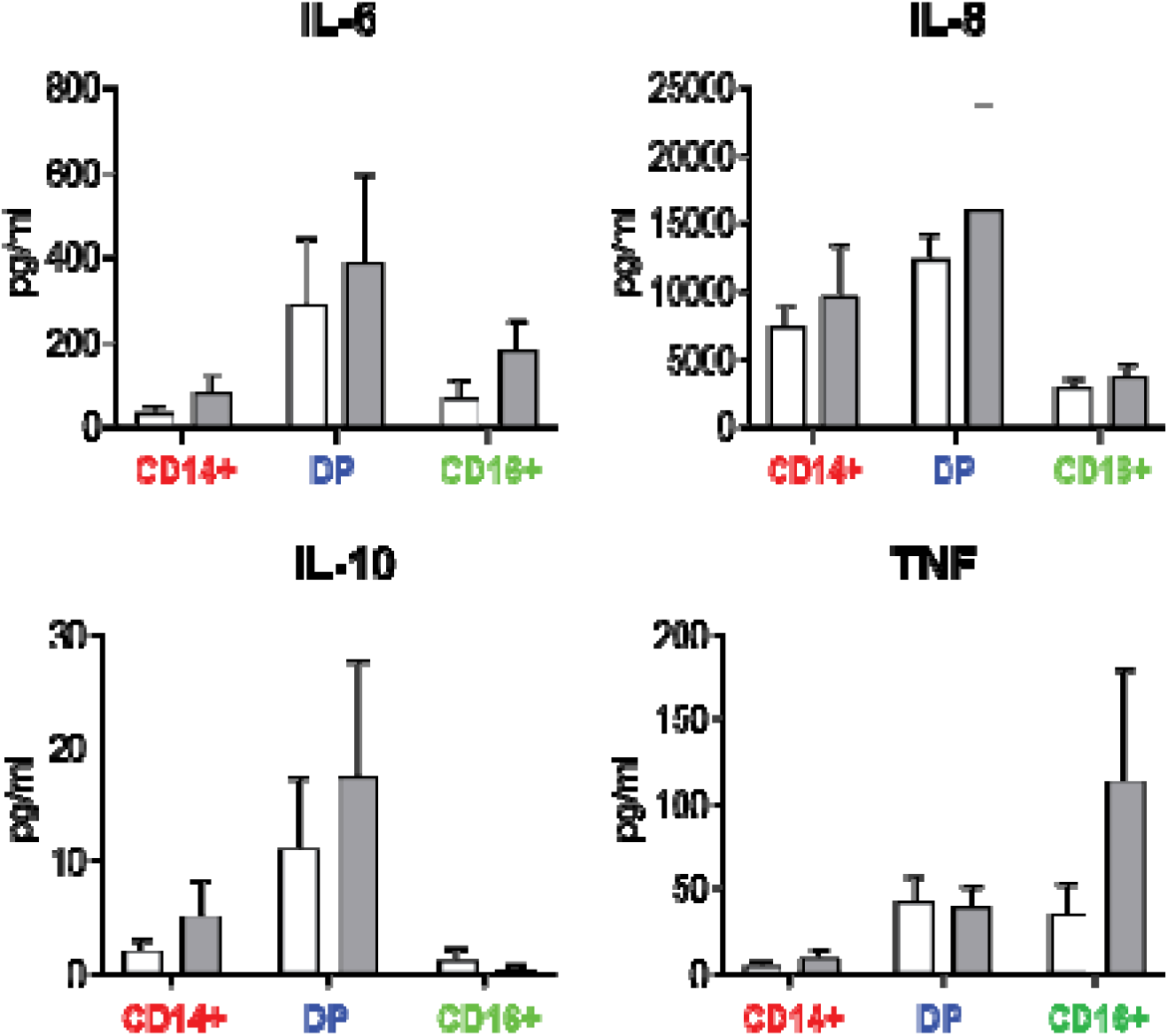
No significant difference in cytokine production from monocytes with age. Monocytes were isolated from the peripheral blood and sorted into three populations classical (CD14+), intermediate (CD14+CD16+) and non-classical (CD16+), cells were cultured (unstimulated) for 24 hours. Cumulative data showing cytokine production from sorted populations separated according to age monocyte

**Supplementary Figure 4:**
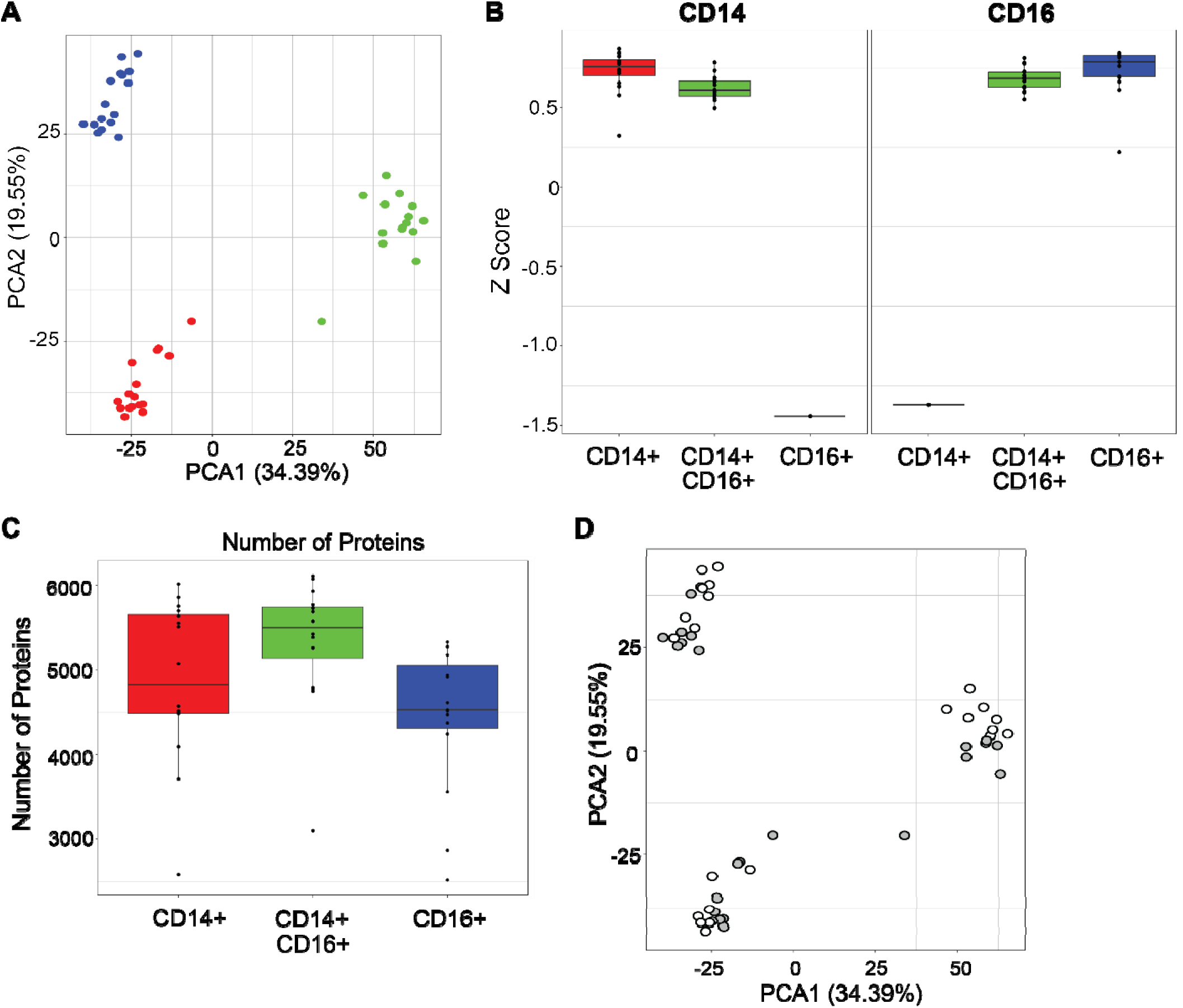
Protein expression within the sorted monocyte populations. Monocytes were isolated from the peripheral blood and sorted into three populations classical (CD14+; red), intermediate (CD14+CD16+; green) and non-classical (CD16+; blue), proteomics analysis was performed on the cell pellets. **A,** PCA analysis of the sorted monocyte populations and **B,** CD14 and CD16 protein expression in the sorted monocyte populations to ensure successful sort and **C,** total protein number in the sorted monocyte populations and **D,** PCA analysis separated according to age of donor with young (white) and old (grey).

**Supplementary Figure 5:**
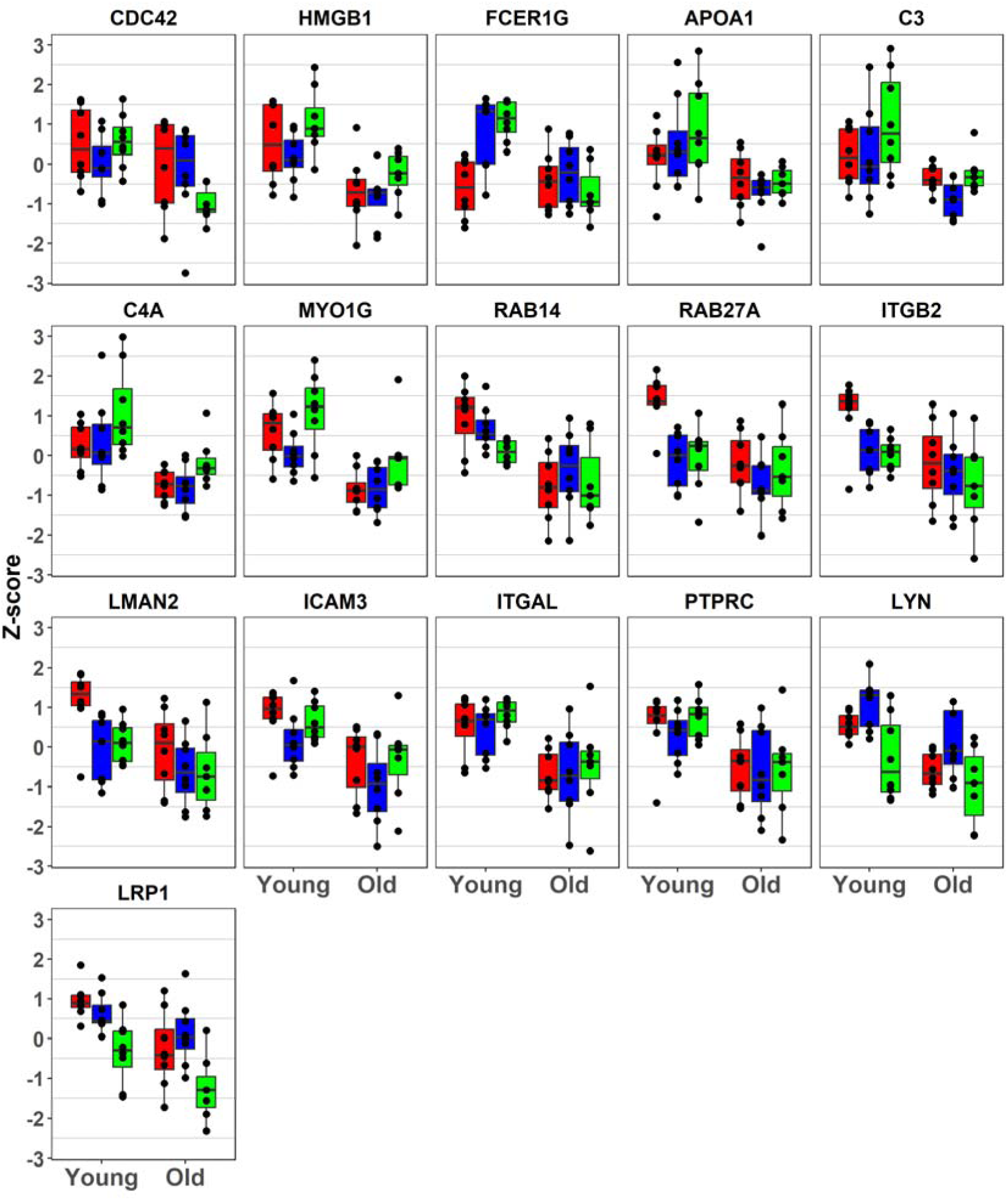
Phagocytosis proteins are significantly downregulated in older monocytes. Phagocytosis associated protein expression classical (CD14+; red), intermediate (CD14+CD16+; green) and non-classical (CD16+; blue) in sorted monocyte populations.

**Supplementary Figure 6:**
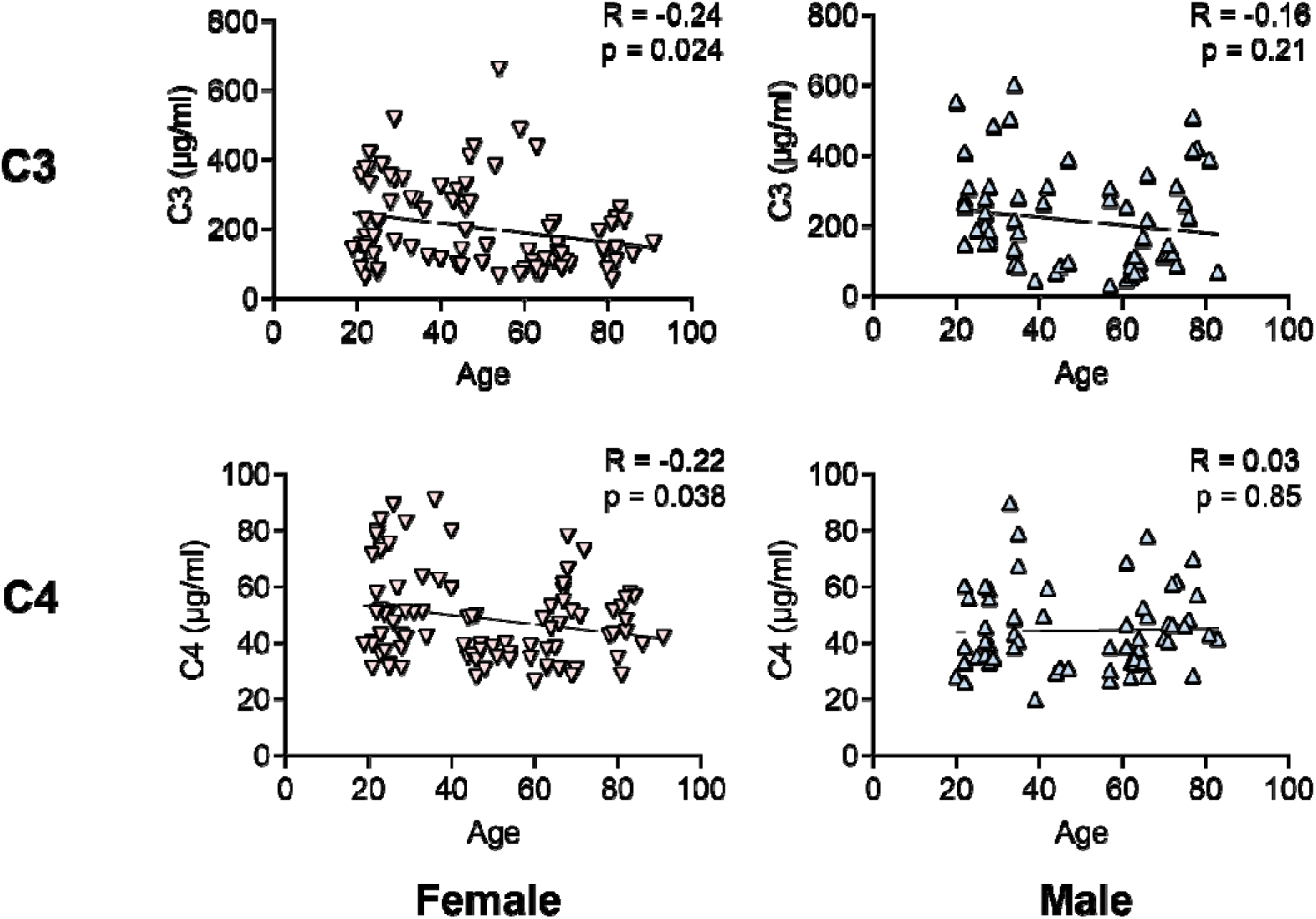
Male sex does not correlate with Complement proteins. Serum samples were assessed for C3 and C4 concentrations by ELISA. C3 and C4 serum concentration was correlated in female (pink downwards triangle) and in males (blue upwards triangle). Assessed by Pearson’s correlation test.

**Supplementary Figure 7:**
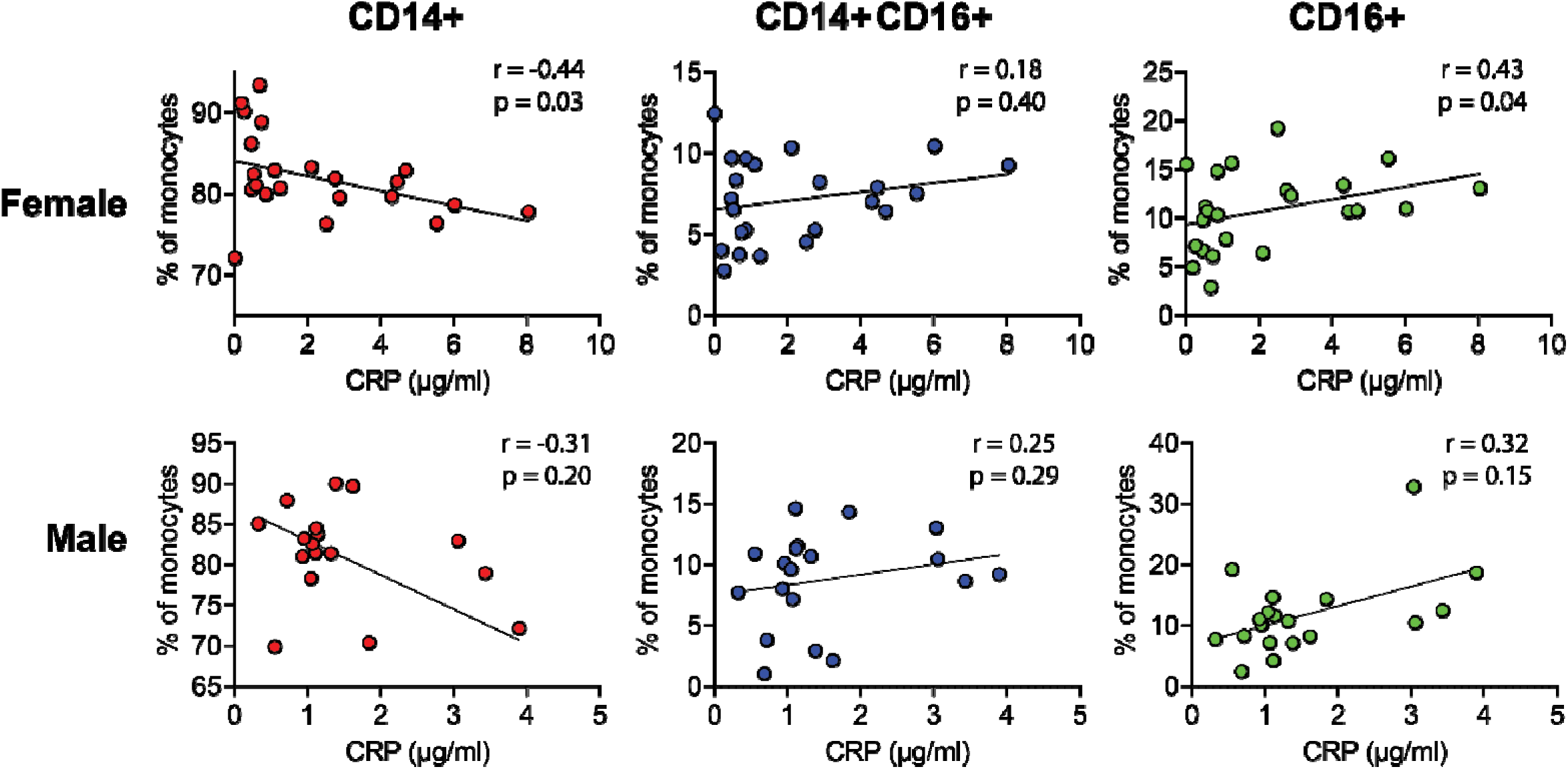
The frequency of monocyte populations correlated with serum CRP split according to biological sex. Serum C Reactive Protein (CRP) concentrations were assessed by ELISA and correlated with frequency of monocyte populations in the peripheral blood. Data was split according to whether the donor was female (top) or male (bottom). Assessed by Pearson’s correlation test.

## Table Legends

**Supplementary Table 1** Table of differentially regulated protein found in the heatmap of Figure 3A.

**Supplementary Table 2** Table of differentially regulated proteins according to monocyte cell type.

**Supplementary Table 3** Table of age-associated differentially regulated proteins.

