## Supplementary Table 1 for "Age-associated inflammatory monocytes are increased in menopausal females and reversed by Hormone Replacement Therapy"

| Heatmap order | Protein Accessions | Gene Names | Protein Descriptions |
|---|---|---|---|
| 1 | P32455 | GBP1 | Guanylate-binding protein 1 |
| 2 | P12277 | CKB | Creatine kinase B-type |
| 3 | P43490 | NAMPT | Nicotinamide phosphoribosyltransferase |
| 4 | P04179 | SOD2 | Superoxide dismutase [Mn], mitochondrial |
| 5 | Q9BW30 | TPPP3 | Tubulin polymerization-promoting protein family member 3 |
| 6 | Q08257 | CRYZ | Quinone oxidoreductase |
| 7 | Q8IV04 | TBC1D10C | Carabin |
| 8 | Q9HC16 | APOBEC3G | DNA dC->dU-editing enzyme APOBEC-3G |
| 9 | O14796 | SH2D1B | SH2 domain-containing protein 1B |
| 10 | P51608-2 | MECP2 | Isoform B of Methyl-CpG-binding protein 2 |
| 11 | Q86YV0 | RASAL3 | RAS protein activator like-3 |
| 12 | P53634 | CTSC | Dipeptidyl peptidase 1 |
| 13 | P14222 | PRF1 | Perforin-1 |
| 14 | P42330 | AKR1C3 | Aldo-keto reductase family 1 member C3 |
| 15 | O76096 | CST7 | Cystatin-F |
| 16 | P59768 | GNG2 | Guanine nucleotide-binding protein G(I)/G(S)/G(O) subunit gamma-2 |
| 17 | Q7Z7L1 | SLFN11 | Schlafen family member 11 |
| 18 | Q7Z4H3 | HDDC2 | 5'-deoxynucleotidase HDDC2 |
| 19 | O75791 | GRAP2 | GRB2-related adapter protein 2 |
| 20 | Q08AF3 | SLFN5 | Schlafen family member 5 |
| 21 | Q9BPX5 | ARPC5L | Actin-related protein 2/3 complex subunit 5-like protein |
| 22 | Q14847 | LASP1 | LIM and SH3 domain protein 1 |
| 23 | Q9UI08-2 | EVL | Isoform 1 of Ena/VASP-like protein |
| 24 | Q9BUH6 | PAXX | C16 |
| 25 | O43670;O43670-2;O43670-4 | ZNF207 | BUB3-interacting and GLEBS motif-containing protein ZNF207;Isoform 2 of BUB3-interacting and GLEBS motif-containing protein ZNF207;Isoform 4 of BUB3-interacting and GLEBS motif-containing protein ZNF207 |
| 26 | P33240;P33240-2 | CSTF2 | Cleavage stimulation factor subunit 2;Isoform 2 of Cleavage stimulation factor subunit 2 |
| 27 | P00813 | ADA | Adenosine deaminase |
| 28 | O95865 | DDAH2 | N(G),N(G)-dimethylarginine dimethylaminohydrolase 2 |
| 29 | Q15942 | ZYX | Zyxin |

| | | | |
|---|---|---|---|
| 30 | Q05315 | CLC | Galectin-10 |
| 31 | P63261 | ACTG1 | Actin, cytoplasmic 2 |
| 32 | Q05519;Q05519-2 | SRSF11 | Serine/arginine-rich splicing factor 11;Isoform 2 of Serine/arginine-rich splicing factor 11 |
| 33 | P01834 | IGKC | Immunoglobulin kappa constant |
| 34 | P0CG38 | POTEI | POTE ankyrin domain family member I |
| 35 | P68871 | HBB | Hemoglobin subunit beta |
| 36 | O43516;O43516-3 | WIPF1 | WAS/WASL-interacting protein family member 1;Isoform 3 of WAS/WASL-interacting protein family member 1 |
| 37 | P08134 | RHOC | Rho-related GTP-binding protein RhoC |
| 38 | Q9P2A4;Q9P2A4-2 | ABI3 | ABI gene family member 3;Isoform 2 of ABI gene family member 3 |
| 39 | O00479 | HMGN4 | High mobility group nucleosome-binding domain-containing protein 4 |
| 40 | P16402 | H1-3 | Histone H1.3 |
| 41 | P22304 | IDS | Iduronate 2-sulfatase |
| 42 | Q8NFU3 | TSTD1 | Thiosulfate:glutathione sulfurtransferase |
| 43 | Q9Y6X5 | ENPP4 | Bis(5'-adenosyl)-triphosphatase ENPP4 |
| 44 | P02452 | COL1A1 | Collagen alpha-1(I) chain |
| 45 | Q04756 | HGFAC | Hepatocyte growth factor activator |
| 46 | Q8WWZ4;Q8WWZ4-2 | ABCA10 | ATP-binding cassette sub-family A member 10;Isoform 2 of ATP-binding cassette sub-family A member 10 |
| 47 | P36955 | SERPINF1 | Pigment epithelium-derived factor |
| 48 | P51884 | LUM | Lumican |
| 49 | Q96RW7;Q96RW7-2 | HMCN1 | Hemicentin-1;Isoform 2 of Hemicentin-1 |
| 50 | Q8IVL0;Q8IVL0-2;Q8IVL0-3 | NAV3 | Neuron navigator 3;Isoform 2 of Neuron navigator 3;Isoform 3 of Neuron navigator 3 |
| 51 | P12109 | COL6A1 | Collagen alpha-1(VI) chain |
| 52 | Q9NYH9 | UTP6 | U3 small nucleolar RNA-associated protein 6 homolog |
| 53 | O95498;O95498-6 | VNN2 | Pantetheine hydrolase VNN2;Isoform 6 of Pantetheine hydrolase VNN2 |
| 54 | P04004 | VTN | Vitronectin |

| | | | |
|---|---|---|---|
| 55 | P05543 | SERPINA7 | Thyroxine-binding globulin |
| 56 | P16403 | H1-2 | Histone H1.2 |
| 57 | Q8IUE6 | H2AC21 | Histone H2A type 2-B |
| 58 | Q9BTD8;Q9BTD8-2;Q9BTD8-3;Q9BTD8-4 | RBM42 | RNA-binding protein 42;Isoform 2 of RNA-binding protein 42;Isoform 3 of RNA-binding protein 42;Isoform 4 of RNA-binding protein 42 |
| 59 | O00151 | PDLIM1 | PDZ and LIM domain protein 1 |
| 60 | P18065 | IGFBP2 | Insulin-like growth factor-binding protein 2 |
| 61 | Q8IWC1;Q8IWC1-3;Q8IWC1-4 | MAP7D3 | MAP7 domain-containing protein 3;Isoform 3 of MAP7 domain-containing protein 3;Isoform 4 of MAP7 domain-containing protein 3 |
| 62 | P69905 | HBA1 | Hemoglobin subunit alpha |
| 63 | Q14118 | DAG1 | Dystroglycan 1 |
| 64 | P18859;P18859-2 | ATP5PF | ATP synthase-coupling factor 6, mitochondrial;Isoform 2 of ATP synthase-coupling factor 6, mitochondrial |
| 65 | P02649 | APOE | Apolipoprotein E |
| 66 | P02656 | APOC3 | Apolipoprotein C-III |
| 67 | P17927 | CR1 | Complement receptor type 1 |
| 68 | P13611 | VCAN | Versican core protein |
| 69 | O95347;O95347-2 | SMC2 | Structural mainte0ce of chromosomes protein 2;Isoform 2 of Structural mainte0ce of chromosomes protein 2 |
| 70 | P25205 | MCM3 | DNA replication licensing factor MCM3 |
| 71 | P02792 | FTL | Ferritin light chain |
| 72 | P09601 | HMOX1 | Heme oxygenase 1 |
| 73 | P31941 | APOBEC3A | DNA dC->dU-editing enzyme APOBEC-3A |
| 74 | P49419;P49419-2 | ALDH7A1 | Alpha-aminoadipic semialdehyde dehydrogenase;Isoform 2 of Alpha-aminoadipic semialdehyde dehydrogenase |
| 75 | P09668 | CTSH | Pro-cathepsin H |
| 76 | P06737;P06737-2 | PYGL | Glycogen phosphorylase, liver form;Isoform 2 of Glycogen phosphorylase, liver form |
| 77 | P08758 | ANXA5 | Annexin A5 |
| 78 | P50225 | SULT1A1 | Sulfotransferase 1A1 |
| 79 | Q9NZK5 | ADA2 | Adenosine deaminase 2 |

| | | | |
|---|---|---|---|
| 80 | O94819 | KBTBD11 | Kelch repeat and BTB domain-containing protein 11 |
| 81 | P50452 | SERPINB8 | Serpin B8 |
| 82 | P50135 | HNMT | Histamine N-methyltransferase |
| 83 | Q8TEH3;Q8TEH3-2;Q8TEH3-3 | DENND1A | DENN domain-containing protein 1A;Isoform 2 of DENN domain-containing protein 1A;Isoform 3 of DENN domain-containing protein 1A |
| 84 | Q7L266 | ASRGL1 | Isoaspartyl peptidase/L-asparaginase |
| 85 | P52209;P52209-2 | PGD | 6-phosphogluconate dehydrogenase, decarboxylating;Isoform 2 of 6-phosphogluconate dehydrogenase, decarboxylating |
| 86 | Q12882 | DPYD | Dihydropyrimidine dehydrogenase [NADP(+)] |
| 87 | O75874 | IDH1 | Isocitrate dehydrogenase [NADP] cytoplasmic |
| 88 | P19878 | NCF2 | Neutrophil cytosol factor 2 |
| 89 | P40121 | CAPG | Macrophage-capping protein |
| 90 | O95716 | RAB3D | Ras-related protein Rab-3D |
| 91 | Q02338 | BDH1 | D-beta-hydroxybutyrate dehydrogenase, mitochondrial |
| 92 | P00738 | HP | Haptoglobin |
| 93 | P08631;P08631-4 | HCK | Tyrosine-protein kinase HCK;Isoform 4 of Tyrosine-protein kinase HCK |
| 94 | Q9H3G5 | CPVL | Probable serine carboxypeptidase CPVL |
| 95 | P05091 | ALDH2 | Aldehyde dehydrogenase, mitochondrial |
| 96 | Q96CX2 | KCTD12 | BTB/POZ domain-containing protein KCTD12 |
| 97 | P30043 | BLVRB | Flavin reductase (NADPH) |
| 98 | P41218 | MNDA | Myeloid cell nuclear differentiation antigen |
| 99 | Q9UBR2 | CTSZ | Cathepsin Z |
| 100 | P07858 | CTSB | Cathepsin B |
| 101 | P23141;P23141-2 | CES1 | Liver carboxylesterase 1;Isoform 2 of Liver carboxylesterase 1 |
| 102 | Q03169 | TNFAIP2 | Tumor necrosis factor alpha-induced protein 2 |

| | | | |
|---|---|---|---|
| 103 | Q12797;Q12797-10 | ASPH | Aspartyl/asparaginyl beta-hydroxylase;Isoform 10 of Aspartyl/asparaginyl beta-hydroxylase |
| 104 | P28676 | GCA | Grancalcin |
| 105 | P05109 | S100A8 | Protein S100-A8 |
| 106 | P06702 | S100A9 | Protein S100-A9 |
| 107 | O00602 | FCN1 | Ficolin-1 |
| 108 | P01903 | HLA-DRA | HLA class II histocompatibility antigen, DR alpha chain |
| 109 | Q04941 | PLP2 | Proteolipid protein 2 |
| 110 | Q9Y336 | SIGLEC9 | Sialic acid-binding Ig-like lectin 9 |
| 111 | P10620 | MGST1 | Microsomal glutathione S-transferase 1 |
| 112 | Q07065 | CKAP4 | Cytoskeleton-associated protein 4 |
| 113 | Q8WY22 | BRI3BP | BRI3-binding protein |
| 114 | O94905 | ERLIN2 | Erlin-2 |
| 115 | Q8NBQ5 | HSD17B11 | Estradiol 17-beta-dehydrogenase 11 |
| 116 | Q96D96 | HVCN1 | Voltage-gated hydrogen channel 1 |
| 117 | Q9H8H3 | METTL7A | Putative methyltransferase-like protein 7A |
| 118 | A6NI72 | NCF1B | Putative neutrophil cytosol factor 1B |
| 119 | O43175 | PHGDH | D-3-phosphoglycerate dehydrogenase |
| 120 | Q15080 | NCF4 | Neutrophil cytosol factor 4 |
| 121 | Q6P4A8 | PLBD1 | Phospholipase B-like 1 |
| 122 | Q13637 | RAB32 | Ras-related protein Rab-32 |
| 123 | P61626 | LYZ | Lysozyme C |
| 124 | P05164;P05164-2;P05164-3 | MPO | Myeloperoxidase;Isoform H14 of Myeloperoxidase;Isoform H7 of Myeloperoxidase |
| 125 | P08311 | CTSG | Cathepsin G |
| 126 | P24158 | PRTN3 | Myeloblastin |
| 127 | P08571 | CD14 | Monocyte differentiation antigen CD14 |
| 128 | O95197-3 | RTN3 | Isoform 3 of Reticulon-3 |
| 129 | Q9NUU6 | OTULINL | Inactive ubiquitin thioesterase OTULINL |
| 130 | P04839 | CYBB | Cytochrome b-245 heavy chain |
| 131 | P13498 | CYBA | Cytochrome b-245 light chain |
| 132 | P10153 | RNASE2 | Non-secretory ribonuclease |
| 133 | Q9UM07 | PADI4 | Protein-arginine deiminase type-4 |
| 134 | P80723 | BASP1 | Brain acid soluble protein 1 |
| 135 | P08246 | ELANE | Neutrophil elastase |
| 136 | P20160 | AZU1 | Azurocidin |
