## Supplementary Table 2 for "Age-associated inflammatory monocytes are increased in menopausal females and reversed by Hormone Replacement Therapy"

| | | CD14+ vs CD14+CD16+ | | | CD14+CD16+ vs CD16+ | | | CD14+ vs CD16+ | | |
|---|---|---|---|---|---|---|---|---|---|---|
| **Protein_id** | **Gene name** | **log2fold change** | **p value** | **qvalue** | **log2fold change** | **p value** | **qvalue** | **log2fold change** | **p value** | **qvalue** |
| P51608-2 | MECP2 | 0.435727671 | 0.020489763 | 0.067019393 | 2.391711137 | 2.63E-12 | 4.15E-10 | 2.555172753 | 1.15E-10 | 7.86E-09 |
| O43583 | DENR | 0.387178923 | 0.008085557 | 0.031712651 | 0.580922159 | 9.13E-05 | 0.000820871 | 0.986418715 | 6.37E-06 | 6.84E-05 |
| Q9UJ70 | NAGK | 0.672555291 | 5.49E-05 | 0.000509383 | -1.326752227 | 5.88E-05 | 0.000580728 | -0.804019133 | 0.011065955 | 0.033982685 |
| P06744 | GPI | -1.94E-15 | 0.999999916 | 1 | -0.624637451 | 0.011027536 | 0.038748383 | -1.068007999 | 4.71E-05 | 0.000366531 |
| P27708 | CAD | -0.226606749 | 0.032458249 | 0.097236861 | -0.505999043 | 0.01374779 | 0.045891431 | -0.673381755 | 0.005537606 | 0.019156724 |
| Q9NXG2 | THUMPD1 | 0.29308227 | 0.07103011 | 0.181487078 | 0.934417243 | 3.68E-07 | 9.06E-06 | 0.924969895 | 0.000106082 | 0.000732809 |
| Q16831 | UPP1 | 0.341729932 | 0.042285035 | 0.120142654 | 0.523434145 | 0.001546992 | 0.008084015 | 0.613960298 | 0.003256872 | 0.012192152 |
| Q96C86 | DCPS | 0.591569826 | 2.93E-06 | 3.88E-05 | 0.745823311 | 1.23E-06 | 2.52E-05 | 1.096322378 | 7.11E-09 | 2.37E-07 |
| P09467 | FBP1 | 0.430047513 | 0.000895877 | 0.005186969 | -1.158212147 | 1.77E-05 | 0.000215295 | -0.776059673 | 0.003095051 | 0.01170443 |
| P04040 | CAT | 0.352384584 | 0.010880243 | 0.040171451 | -0.914551417 | 6.84E-05 | 0.000645829 | -0.73812288 | 0.000927348 | 0.004380455 |
| Q9Y4E8 | USP15 | 0.008028629 | 0.722008216 | 1 | -0.749985256 | 1.57E-05 | 0.000196614 | -1.18890531 | 1.32E-11 | 1.50E-09 |
| P09874 | PARP1 | -0.102611363 | 0.248245547 | 0.483598784 | 1.061053491 | 1.28E-19 | 1.19E-16 | 0.509426554 | 0.000219738 | 0.001303377 |
| Q01081 | U2AF1 | 0.640440034 | 0.008822163 | 0.034108495 | 0.616330729 | 0.004481371 | 0.019010235 | 1.053824334 | 0.000160282 | 0.001010709 |
| Q13637 | RAB32 | -4.32E-17 | 0.999999991 | 1 | -1.658278102 | 7.36E-07 | 1.60E-05 | -1.391572288 | 1.69E-05 | 0.000153134 |
| P33240;P33240-2 | CSTF2 | 0.755375511 | 0.002071839 | 0.010285411 | 1.042953362 | 9.77E-08 | 2.88E-06 | 1.547977527 | 7.71E-09 | 2.54E-07 |
| P06737;P06737-2 | PYGL | -0.023589787 | 0.600456673 | 0.993528967 | -1.255755273 | 0.000188202 | 0.001453767 | -1.823494595 | 3.41E-07 | 5.87E-06 |
| Q8WVY7 | UBLCP1 | 0.203356062 | 0.05520747 | 0.149739645 | 0.973866692 | 5.04E-10 | 2.88E-08 | 1.014350964 | 4.22E-08 | 1.10E-06 |
| Q00610;Q00610-2 | CLTC | 6.32E-16 | 0.999999962 | 1 | -0.634903933 | 4.28E-06 | 7.01E-05 | -0.684112019 | 0.000830751 | 0.003989447 |
| P60981 | DSTN | 0.502804636 | 0.000234792 | 0.001686104 | 0.87949583 | 2.68E-09 | 1.17E-07 | 1.119672477 | 5.42E-08 | 1.34E-06 |
| Q96C19 | EFHD2 | 0.749321087 | 3.71E-11 | 2.18E-09 | 0.890444863 | 3.55E-10 | 2.21E-08 | 1.330327582 | 2.04E-15 | 6.61E-13 |
| P82979 | SARNP | 1.627032895 | 7.61E-05 | 0.000662593 | 0.826936222 | 0.008142475 | 0.031063265 | 2.010016241 | 5.96E-05 | 0.000449318 |
| P00338 | LDHA | 0.142611606 | 0.151962521 | 0.330345159 | -0.591013604 | 0.000431951 | 0.002883786 | -0.521297882 | 0.000573844 | 0.002958031 |
| P48426 | PIP4K2A | 0.482118558 | 6.85E-05 | 0.000614396 | 1.134866151 | 2.67E-12 | 4.15E-10 | 1.364548976 | 6.27E-15 | 1.63E-12 |
| Q9UI08-2 | EVL | 2.141250278 | 5.59E-43 | 1.97E-39 | 1.348727333 | 1.45E-10 | 1.10E-08 | 3.085532738 | 4.06E-18 | 2.63E-15 |
| Q01469 | FABP5 | -0.368343685 | 0.001794596 | 0.009085442 | -0.566614693 | 0.012164551 | 0.041749572 | -1.167461534 | 7.38E-06 | 7.67E-05 |
| Q04917 | YWHAH | 0.059031243 | 0.50936262 | 0.86471478 | -0.815972248 | 1.36E-10 | 1.06E-08 | -0.646797692 | 3.98E-05 | 0.000318753 |
| P62258 | YWHAE | 0.257827774 | 0.003632789 | 0.016401042 | -0.835234932 | 1.01E-11 | 1.18E-09 | -0.780561257 | 1.50E-09 | 6.30E-08 |
| P30740 | SERPINB1 | -0.275462857 | 0.075547317 | 0.190188187 | -0.722659074 | 0.000631117 | 0.003923843 | -1.372099597 | 1.88E-07 | 3.70E-06 |
| P37837 | TALDO1 | 0.31548715 | 0.005469346 | 0.022964643 | -0.75946021 | 0.000856722 | 0.004973597 | -0.596281688 | 0.009592397 | 0.030134279 |
| P50452 | SERPINB8 | -0.264473793 | 0.04349557 | 0.123284068 | -1.155875551 | 1.47E-06 | 2.89E-05 | -1.697218695 | 3.00E-10 | 1.69E-08 |
| P52790 | HK3 | 0.506635511 | 0.000265741 | 0.001864433 | -1.384951838 | 0.001245718 | 0.006717292 | -1.395984866 | 0.001881996 | 0.007785711 |
| P50225 | SULT1A1 | 0.295501412 | 0.150812313 | 0.328958916 | -1.497369978 | 0.000116062 | 0.000977288 | -1.37013428 | 0.000445293 | 0.002375507 |
| O14745 | SLC9A3R1 | 0.225144132 | 0.085466815 | 0.209745203 | 1.610396745 | 1.98E-20 | 5.56E-17 | 1.688846388 | 1.96E-13 | 3.63E-11 |
| Q8ND71 | GIMAP8 | 0.386295457 | 0.00108799 | 0.00605087 | -0.74215579 | 2.71E-05 | 0.000308923 | -0.576362398 | 2.03E-05 | 0.000180916 |
| P10412 | H1-4 | 0.469749872 | 0.134411397 | 0.301454077 | 1.010167322 | 0.011882611 | 0.041033057 | 1.523631918 | 0.000994306 | 0.004671259 |
| P16402 | H1-3 | 1.523814224 | 0.002016493 | 0.010042593 | 1.536276752 | 0.010119937 | 0.036473397 | 3.020124095 | 1.99E-05 | 0.000178402 |
| Q15102 | PAFAH1B3 | 0 | 1 | 1 | -0.710991871 | 0.000395779 | 0.002726693 | -1.089264247 | 1.18E-09 | 5.20E-08 |
| Q07955 | SRSF1 | 0.110192854 | 0.430318047 | 0.753729701 | 0.511887183 | 0.005664751 | 0.023087154 | 0.603918685 | 0.005838627 | 0.019880412 |
| P19878 | NCF2 | 0.068165491 | 0.493743436 | 0.841846884 | -2.967819528 | 3.60E-08 | 1.17E-06 | -3.797735995 | 1.39E-12 | 2.06E-10 |
| O43768;O43768-2 | ENSA | 0.889391406 | 0.004125587 | 0.018160824 | 0.581794324 | 0.011395173 | 0.039692008 | 1.174176143 | 0.000169044 | 0.00105318 |
| P06702 | S100A9 | 0.255410186 | 0.145305418 | 0.319020488 | -3.178246486 | 1.21E-13 | 3.09E-11 | -3.022987782 | 2.46E-10 | 1.45E-08 |
| P21283 | ATP6V1C1 | 5.79E-16 | 0.99999997 | 1 | -0.975846615 | 3.76E-05 | 0.000404395 | -1.094416446 | 5.29E-06 | 6.00E-05 |

| | | | | | | | | | | |
|---|---|---|---|---|---|---|---|---|---|---|
| Q15149 | PLEC | 0.346935344 | 0.011287393 | 0.041457655 | -1.240156821 | 5.78E-12 | 7.37E-10 | -1.00534381 | 4.48E-07 | 7.51E-06 |
| Q96D96 | HVCN1 | -6.43E-17 | 0.999999995 | 1 | -1.596488622 | 4.05E-05 | 0.000421771 | -1.774383728 | 3.89E-05 | 0.000314848 |
| Q9NVA2 | SEPTIN11 | 7.36E-16 | 0.999999947 | 1 | 0.754940417 | 1.28E-07 | 3.56E-06 | 0.566521452 | 3.96E-05 | 0.000318326 |
| P02768 | ALB | 0 | 1 | 1 | 0.847589698 | 0.004823736 | 0.02021787 | 0.873817268 | 0.00232879 | 0.009222859 |
| P51452 | DUSP3 | 0.428997223 | 0.003403665 | 0.015525644 | -1.183000878 | 0.000231916 | 0.001762313 | -0.996334553 | 0.001233169 | 0.005542862 |
| Q14554 | PDIA5 | -0.143446721 | 0.232711725 | 0.460978394 | -0.848107252 | 0.006429833 | 0.025598295 | -0.939324528 | 0.003172134 | 0.011926489 |
| P35475 | IDUA | 0.253511209 | 0.050895584 | 0.140091981 | 0.998681235 | 3.74E-05 | 0.000402954 | 1.126569221 | 3.50E-06 | 4.21E-05 |
| P11586 | MTHFD1 | -1.07E-17 | 0.999999998 | 1 | -0.528989091 | 0.000104805 | 0.000912646 | -0.836602664 | 1.03E-08 | 3.28E-07 |
| Q709C8;Q709C8-3 | VPS13C | 1.41E-15 | 0.99999992 | 1 | -0.53155076 | 7.50E-06 | 0.000110739 | -0.705167967 | 8.56E-07 | 1.32E-05 |
| P08311 | CTSG | -0.545406184 | 0.084347381 | 0.207978227 | -3.012987709 | 5.33E-07 | 1.27E-05 | -4.637119632 | 1.23E-09 | 5.33E-08 |
| P05091 | ALDH2 | 0.278342413 | 0.034666495 | 0.102459397 | -2.41303816 | 9.65E-09 | 3.51E-07 | -2.350301708 | 1.33E-08 | 4.10E-07 |
| Q7L1T6 | CYB5R4 | 0.028059521 | 0.664716254 | 1 | 1.148498682 | 0.004203135 | 0.018103826 | 1.159059099 | 0.002961904 | 0.01128303 |
| Q7Z3J2 | VPS35L | 1.31E-16 | 0.999999983 | 1 | -0.677264161 | 0.000319912 | 0.002270969 | -1.040685952 | 2.48E-06 | 3.17E-05 |
| P09769 | FGR | 0.251706825 | 0.014702282 | 0.051023864 | -1.120585233 | 2.64E-09 | 1.17E-07 | -1.079305734 | 1.39E-09 | 5.92E-08 |
| P08631;P08631-4 | HCK | 0.709938131 | 1.00E-08 | 2.80E-07 | -1.465575653 | 1.48E-05 | 0.000188752 | -1.174799633 | 1.81E-05 | 0.000162452 |
| Q9P289 | STK26 | 0.430149581 | 9.60E-05 | 0.000809666 | 0.757160869 | 8.03E-13 | 1.61E-10 | 1.030424582 | 1.39E-10 | 9.24E-09 |
| P31146 | CORO1A | 0.841101049 | 1.48E-11 | 9.86E-10 | 0.657100779 | 4.82E-06 | 7.68E-05 | 1.226363405 | 7.13E-11 | 5.73E-09 |
| Q9HAF1;Q9HAF1-2;Q9HAF1-3;Q9HAF1-4 | MEAF6 | 0.590569125 | 0.028221337 | 0.08690693 | 0.818065105 | 0.000264074 | 0.001943476 | 1.058107339 | 0.005691498 | 0.019517619 |
| P0DOY3 | IGLC3 | 1.035965966 | 3.18E-08 | 7.38E-07 | 0.631558258 | 0.000555398 | 0.003531376 | 1.381707435 | 2.11E-08 | 5.90E-07 |
| P09104 | ENO2 | 3.61E-17 | 0.999999997 | 1 | -0.49892957 | 0.010574446 | 0.037627851 | -1.376658086 | 2.81E-07 | 5.12E-06 |
| P06733 | ENO1 | 0.437127083 | 0.001431928 | 0.007502196 | -1.063126365 | 1.68E-07 | 4.57E-06 | -0.887837485 | 6.60E-05 | 0.000488774 |
| Q96QK1 | VPS35 | 6.49E-17 | 0.999999991 | 1 | -0.520592769 | 0.013045538 | 0.043965971 | -0.798957476 | 0.000141904 | 0.000926301 |
| Q99536 | VAT1 | 5.66E-18 | 0.999999997 | 1 | -0.560327137 | 9.91E-05 | 0.000868353 | -0.54028067 | 0.001080573 | 0.005032592 |
| O43707 | ACTN4 | 0.134110234 | 0.134483058 | 0.301454077 | 0.71996765 | 3.82E-09 | 1.58E-07 | 0.717582493 | 2.46E-06 | 3.16E-05 |
| O94973 | AP2A2 | 1.50E-13 | 0.999999126 | 1 | -0.469018159 | 0.001147869 | 0.006298676 | -0.650308805 | 1.37E-06 | 1.96E-05 |
| Q14141;Q14141-2;Q14141-4 | SEPTIN6 | 0.031735669 | 0.605760843 | 1 | 0.910257795 | 6.03E-08 | 1.88E-06 | 0.781718314 | 3.32E-06 | 4.01E-05 |
| P00441 | SOD1 | 0.328734096 | 0.029906044 | 0.090825762 | 0.605449065 | 2.26E-06 | 4.16E-05 | 0.751359721 | 2.13E-05 | 0.000187612 |
| P51688 | SGSH | 1.50E-16 | 0.99999998 | 1 | 1.396486038 | 3.84E-07 | 9.28E-06 | 1.268627728 | 9.22E-07 | 1.41E-05 |
| O75165 | DNAJC13 | -1.53E-14 | 0.999999743 | 1 | -1.401918393 | 4.00E-05 | 0.000421771 | -1.634141187 | 4.35E-06 | 5.02E-05 |
| Q9NZN3 | EHD3 | 0 | 1 | 1 | 0.614641997 | 9.09E-06 | 0.000126772 | 0.858249722 | 1.81E-08 | 5.39E-07 |
| Q9UHD8-5 | SEPTIN9 | 0.620346527 | 8.51E-07 | 1.33E-05 | 0.483029844 | 0.000890769 | 0.005128783 | 0.831861364 | 6.41E-07 | 1.03E-05 |
| Q12882 | DPYD | 4.11E-16 | 0.999999978 | 1 | -1.493168927 | 3.60E-05 | 0.000389241 | -1.476985092 | 1.71E-05 | 0.000154594 |
| P29966 | MARCKS | 1.021514738 | 3.72E-06 | 4.76E-05 | -1.631924613 | 4.91E-06 | 7.77E-05 | -1.03427584 | 0.003203745 | 0.012027934 |
| P49662;P49662-2 | CASP4 | 0.043242384 | 0.518976918 | 0.877235194 | 0.804578189 | 0.000952166 | 0.005404601 | 0.560277827 | 0.005575008 | 0.019260465 |
| Q05655 | PRKCD | 0 | 1 | 1 | -0.825295589 | 1.89E-05 | 0.000225745 | -1.2466726 | 4.77E-10 | 2.48E-08 |
| P14618 | PKM | 0.445201805 | 0.002439196 | 0.011781652 | -1.398214101 | 3.23E-11 | 2.92E-09 | -1.051113876 | 9.63E-06 | 9.55E-05 |
| Q9Y5Z4 | HEBP2 | -0.122673665 | 0.276770419 | 0.530088265 | -0.785815667 | 0.000669003 | 0.004087081 | -1.380407456 | 7.17E-08 | 1.68E-06 |
| P46459 | NSF | 0.191474347 | 0.022173898 | 0.071467244 | -0.528270249 | 6.61E-05 | 0.00063052 | -0.604101677 | 2.61E-06 | 3.29E-05 |
| Q14166 | TTLL12 | 0.266968758 | 0.006246263 | 0.025729351 | -0.601038183 | 0.000437656 | 0.00290803 | -0.572259971 | 0.002380922 | 0.009414971 |
| Q6NVY1 | HIBCH | 0.159309471 | 0.266749817 | 0.513125943 | 0.533724313 | 7.19E-05 | 0.000667948 | 0.601131448 | 0.008807371 | 0.028144589 |
| Q684P5;Q684P5-3 | RAP1GAP2 | 0.631421817 | 3.27E-08 | 7.53E-07 | 0.890807002 | 8.40E-14 | 2.62E-11 | 1.266179229 | 5.05E-16 | 1.88E-13 |

| | | | | | | | | | | |
|---|---|---|---|---|---|---|---|---|---|---|
| Q9H3G5 | CPVL | 0.927844774 | 2.35E-08 | 5.57E-07 | -3.002957926 | 1.42E-13 | 3.33E-11 | -2.166560635 | 1.83E-08 | 5.41E-07 |
| Q9HB90 | RRAGC | -5.27E-13 | 0.999998887 | 1 | -0.609856222 | 0.000266332 | 0.001949856 | -1.19541584 | 6.89E-10 | 3.25E-08 |
| O95445 | APOM | 0 | 1 | 1 | 1.233556496 | 0.011461507 | 0.039873529 | 0.964405767 | 0.014139063 | 0.041319781 |
| Q8NBQ5 | HSD17B11 | 0 | 1 | 1 | -1.174852857 | 0.000535876 | 0.003430588 | -1.699662526 | 0.000339265 | 0.001889281 |
| O43813 | LANCL1 | 0 | 1 | 1 | 0.765480072 | 4.45E-06 | 7.13E-05 | 0.509858042 | 0.000348477 | 0.00192365 |
| O00115;O00115-2 | DNASE2 | 0.992766722 | 2.94E-07 | 5.13E-06 | 1.232995105 | 9.78E-12 | 1.18E-09 | 2.152687289 | 3.56E-11 | 3.31E-09 |
| O15067 | PFAS | 5.16E-23 | 1 | 1 | -0.625769865 | 0.004164 | 0.017990533 | -0.813613146 | 0.000107213 | 0.000736877 |
| O00442 | RTCA | 0.586262182 | 0.000164978 | 0.001243684 | 0.510463598 | 0.004234632 | 0.018183627 | 0.740527681 | 0.000215701 | 0.001290741 |
| Q96P48 | ARAP1 | -3.18E-18 | 0.999999999 | 1 | -0.591746341 | 0.003256028 | 0.01486955 | -0.955442106 | 6.91E-06 | 7.24E-05 |
| Q08257 | CRYZ | 1.09E-17 | 0.999999996 | 1 | 1.499558606 | 2.39E-09 | 1.12E-07 | 1.341750404 | 1.24E-06 | 1.84E-05 |
| Q9UBW5 | BIN2 | 0.901376021 | 1.70E-13 | 1.66E-11 | 0.862132447 | 5.87E-06 | 8.94E-05 | 1.288474334 | 1.00E-16 | 5.21E-14 |
| O43760;O43760-2 | SYNGR2 | 2.10E-17 | 0.999999994 | 1 | -1.029885329 | 1.62E-05 | 0.000201517 | -1.186850553 | 1.28E-05 | 0.00012108 |
| Q9HB40 | SCPEP1 | 7.25E-16 | 0.99999995 | 1 | -0.822760876 | 0.000723628 | 0.004307969 | -1.037147859 | 3.31E-05 | 0.000274381 |
| P32455 | GBP1 | 1.983097415 | 7.99E-15 | 1.17E-12 | -0.622156953 | 0.001667786 | 0.008588154 | 1.000228002 | 0.000654708 | 0.003290003 |
| P0C0S8;Q96KK5;Q99878 | H2AC11;H2AC12;H2AC14 | 0.722977983 | 0.030810833 | 0.093027943 | 1.07981191 | 0.004551379 | 0.019238836 | 1.728857557 | 1.60E-05 | 0.00014661 |
| P52209;P52209-2 | PGD | -0.084486563 | 0.421557397 | 0.74320569 | -1.342539008 | 1.21E-05 | 0.000163468 | -1.902117147 | 2.73E-09 | 1.06E-07 |
| Q99439 | CNN2 | 1.048504208 | 4.37E-06 | 5.53E-05 | 0.727374705 | 0.00021929 | 0.001680029 | 1.616074577 | 6.38E-08 | 1.51E-06 |
| P30038 | ALDH4A1 | -0.025608075 | 0.635902974 | 1 | -0.82056582 | 0.000660561 | 0.004052982 | -1.443645357 | 8.11E-08 | 1.86E-06 |
| P23141;P23141-2 | CES1 | 1.10E-17 | 0.999999999 | 1 | -2.905007342 | 2.47E-10 | 1.76E-08 | -2.866395068 | 1.80E-08 | 5.39E-07 |
| Q92947 | GCDH | 0.654626175 | 0.000799635 | 0.004746657 | 0.623218559 | 2.41E-05 | 0.000280455 | 0.954111909 | 3.00E-05 | 0.000252233 |
| P21964;P21964-2 | COMT | -1.06E-17 | 0.999999998 | 1 | -0.807523397 | 0.000511146 | 0.003325415 | -0.864359738 | 0.003351574 | 0.012462275 |
| P13489 | RNH1 | 4.21E-16 | 0.999999964 | 1 | -1.085815756 | 1.35E-06 | 2.72E-05 | -1.338731843 | 4.66E-09 | 1.61E-07 |
| P24666 | ACP1 | 0.747708698 | 7.65E-06 | 9.01E-05 | 0.634528902 | 5.48E-06 | 8.53E-05 | 1.118345441 | 2.25E-09 | 8.93E-08 |
| P62979 | RPS27A | 0.12458085 | 0.316276698 | 0.586336298 | 0.51965441 | 0.010072788 | 0.036350192 | 0.657538799 | 6.49E-05 | 0.000483078 |
| Q9UBE0 | SAE1 | 0.269383042 | 0.009444104 | 0.036233126 | 0.556871828 | 3.20E-05 | 0.000351498 | 0.649812347 | 3.15E-05 | 0.000263981 |
| P54802 | NAGLU | 0 | 1 | 1 | 1.163171056 | 1.33E-06 | 2.70E-05 | 0.705633281 | 0.001809016 | 0.007540356 |
| P07237 | P4HB | 2.91E-16 | 0.999999968 | 1 | -0.603659154 | 1.07E-06 | 2.23E-05 | -0.762121451 | 6.35E-08 | 1.51E-06 |
| Q8NBS9 | TXNDC5 | -0.223938784 | 0.035889708 | 0.104931268 | -0.668024891 | 0.001807725 | 0.009229264 | -1.239719033 | 4.19E-08 | 1.10E-06 |
| Q9BPX5 | ARPC5L | 1.98056912 | 1.46E-09 | 5.15E-08 | 0.971772869 | 0.001217319 | 0.006624621 | 2.771632769 | 2.14E-12 | 2.78E-10 |
| O14920 | IKBKB | -0.23841737 | 0.068381146 | 0.17663877 | -0.954726416 | 0.002279428 | 0.011115682 | -0.856969563 | 0.003353014 | 0.012462275 |
| Q9Y3B8;Q9Y3B8-2;Q9Y3B8-3 | REXO2 | 0 | 1 | 1 | 0.905130087 | 0.000416827 | 0.002823146 | 0.718664927 | 0.007481687 | 0.024418874 |
| P01034 | CST3 | 0.427946364 | 0.000533099 | 0.003344675 | -0.684193337 | 0.00159022 | 0.008272684 | -0.614942835 | 0.004435112 | 0.015719538 |
| P42331-4;P42331-6 | ARHGAP25 | 0.436969684 | 0.00150273 | 0.007815082 | 0.729441472 | 6.13E-07 | 1.40E-05 | 0.781115665 | 1.12E-05 | 0.000108645 |
| Q96E39 | RBMXL1 | 0.291280147 | 0.091698799 | 0.221762664 | 0.604111817 | 0.000889666 | 0.005128783 | 0.740317056 | 0.000378197 | 0.002051264 |
| Q13451 | FKBP5 | 6.12E-17 | 0.999999989 | 1 | -0.576799462 | 7.84E-05 | 0.000716331 | -0.7797058 | 8.01E-06 | 8.23E-05 |
| Q8WXA9-2 | SREK1 | 0.299082375 | 0.093317366 | 0.224598657 | 0.781923935 | 9.67E-05 | 0.000852841 | 0.881190984 | 6.87E-05 | 0.000505329 |
| Q00577 | PURA | 0.506670381 | 5.53E-05 | 0.000510219 | 0.925204438 | 5.61E-10 | 3.14E-08 | 1.202834399 | 3.19E-16 | 1.38E-13 |
| P57088 | TMEM33 | 2.05E-16 | 0.999999993 | 1 | -1.250247602 | 0.001144138 | 0.006290514 | -1.286482425 | 0.00436768 | 0.015565475 |
| P08758 | ANXA5 | 0.920325907 | 2.00E-07 | 3.75E-06 | -1.638503651 | 2.68E-05 | 0.000306657 | -0.955773633 | 0.014045835 | 0.041093559 |
| P05109 | S100A8 | 0.257460787 | 0.225277016 | 0.449817109 | -3.326740007 | 1.05E-15 | 4.92E-13 | -3.146995454 | 9.67E-11 | 6.98E-09 |
| P01857;P01857-1;P0DOX5 | IGHG1 | 0.400672652 | 0.073516457 | 0.186086884 | 0.729315996 | 0.000251505 | 0.001870608 | 1.081399542 | 8.92E-06 | 9.06E-05 |

| | | | | | | | | | | |
|---|---|---|---|---|---|---|---|---|---|---|
| O43516;O43516-3 | WIPF1 | 2.071992853 | 7.51E-05 | 0.000655434 | 0.767689783 | 0.007675108 | 0.029820992 | 2.336705062 | 0.000267521 | 0.001527516 |
| Q14005;Q14005-2;Q14005-3 | IL16 | 0.626143939 | 1.24E-09 | 4.56E-08 | 1.07995548 | 7.95E-18 | 5.57E-15 | 1.13310255 | 4.27E-11 | 3.83E-09 |
| Q8NBJ5 | COLGALT1 | 0.379607075 | 0.000121846 | 0.000972011 | -1.24398794 | 6.12E-05 | 0.00059587 | -0.772178946 | 0.007922489 | 0.025664123 |
| O95865 | DDAH2 | 0.762362116 | 0.005732382 | 0.023807279 | 1.153014621 | 4.33E-11 | 3.57E-09 | 1.791750362 | 8.48E-08 | 1.91E-06 |
| P28676 | GCA | 0 | 1 | 1 | -2.452467852 | 2.05E-07 | 5.42E-06 | -2.638888477 | 3.41E-08 | 9.13E-07 |
| Q86YV0 | RASAL3 | 0.76036021 | 1.37E-06 | 2.01E-05 | 1.212650618 | 2.53E-10 | 1.76E-08 | 1.717586137 | 2.64E-11 | 2.64E-09 |
| Q96S97 | MYADM | -9.70E-07 | 0.998655703 | 1 | -0.99020461 | 0.006244026 | 0.025083452 | -1.312257793 | 0.002916482 | 0.011142678 |
| P47897 | QARS1 | 4.47E-16 | 0.999999975 | 1 | -0.514867758 | 0.001634815 | 0.008457602 | -0.508689077 | 0.00199446 | 0.008172882 |
| P42785 | PRCP | 0.395802008 | 7.48E-05 | 0.000654345 | 0.604945506 | 4.17E-05 | 0.000430239 | 0.875315837 | 4.52E-09 | 1.59E-07 |
| P40121 | CAPG | -0.016178432 | 0.763691947 | 1 | -2.259951759 | 1.21E-06 | 2.49E-05 | -3.067040626 | 3.23E-09 | 1.22E-07 |
| P22087 | FBL | 0.563829302 | 4.09E-07 | 6.86E-06 | 0.607029839 | 8.65E-10 | 4.58E-08 | 0.93613589 | 2.31E-11 | 2.40E-09 |
| P49189 | ALDH9A1 | 0.561055556 | 4.85E-06 | 6.06E-05 | 0.47337017 | 0.000500974 | 0.003282084 | 0.805765232 | 4.84E-07 | 7.96E-06 |
| P46063 | RECQL | 0.560444568 | 4.51E-05 | 0.000429357 | 0.584374773 | 3.05E-05 | 0.000340401 | 0.896941939 | 2.23E-07 | 4.26E-06 |
| Q96B97 | SH3KBP1 | 0.473855571 | 4.56E-09 | 1.33E-07 | 0.648473843 | 1.74E-07 | 4.70E-06 | 0.827073803 | 1.10E-08 | 3.43E-07 |
| Q9NZM1;Q9NZM1-6 | MYOF | 0.573217971 | 0.005026784 | 0.021510242 | -1.082919114 | 4.64E-05 | 0.00046932 | -0.794681247 | 0.000633018 | 0.00321788 |
| O75695 | RP2 | 7.96E-19 | 1 | 1 | -0.918425335 | 0.001696888 | 0.008714421 | -1.108925426 | 0.001570062 | 0.006708917 |
| Q9BRR6;Q9BRR6-2 | ADPGK | -0.005593006 | 0.884169214 | 1 | -0.77495899 | 0.000796434 | 0.004662218 | -1.103153337 | 1.29E-05 | 0.00012149 |
| Q96BX8 | MOB3A | 0.159703967 | 0.23625764 | 0.465908524 | 0.556216024 | 0.000405446 | 0.002780598 | 0.567968268 | 0.003628199 | 0.013276144 |
| P51149 | RAB7A | 7.47E-17 | 0.999999986 | 1 | -0.947089814 | 7.23E-09 | 2.74E-07 | -1.079846967 | 5.29E-11 | 4.58E-09 |
| P14625 | HSP90B1 | -0.120505781 | 0.055728107 | 0.150919589 | -0.533926816 | 8.27E-07 | 1.77E-05 | -0.910347332 | 6.64E-14 | 1.33E-11 |
| P52630;P52630-4 | STAT2 | 4.80E-17 | 0.999999993 | 1 | -0.886301642 | 1.57E-06 | 3.03E-05 | -0.669757109 | 0.000173033 | 0.001070334 |
| Q96SB3 | PPP1R9B | -1.42E-17 | 0.999999998 | 1 | -0.47341756 | 0.011325888 | 0.039581278 | -0.875445625 | 9.39E-06 | 9.41E-05 |
| O00567 | NOP56 | 0.361327385 | 0.001576457 | 0.008138489 | 0.592747469 | 6.56E-06 | 9.89E-05 | 0.778431564 | 4.90E-08 | 1.25E-06 |
| O14974-4 | PPP1R12A | 0.719298271 | 2.07E-08 | 4.97E-07 | 0.851334826 | 2.53E-06 | 4.49E-05 | 1.260680895 | 1.63E-10 | 1.01E-08 |
| Q13951 | CBFB | -3.00E-15 | 0.999999887 | 1 | 1.019022071 | 6.94E-20 | 9.72E-17 | 0.591562927 | 2.60E-06 | 3.29E-05 |
| O43815 | STRN | 0.598159296 | 0.000136474 | 0.001066977 | 0.530203805 | 7.07E-05 | 0.000661127 | 0.897544445 | 1.42E-06 | 1.99E-05 |
| Q13442 | PDAP1 | 0.204223061 | 0.082278219 | 0.203588071 | 0.525606977 | 1.70E-05 | 0.000211124 | 0.537934068 | 0.000198532 | 0.001205109 |
| P08670 | VIM | 0.729852145 | 3.48E-06 | 4.53E-05 | -1.118889686 | 6.29E-10 | 3.46E-08 | -0.550396994 | 0.003417234 | 0.012664729 |
| P36957 | DLST | 0.652288657 | 0.000216682 | 0.001578559 | 1.038323911 | 3.93E-07 | 9.43E-06 | 1.156827787 | 5.80E-10 | 2.84E-08 |
| Q96CN7 | ISOC1 | 0.426718348 | 0.006546027 | 0.026683574 | 0.609207726 | 0.000522389 | 0.003367305 | 0.806441444 | 1.24E-05 | 0.000119751 |
| P38606 | ATP6V1A | 0.361058977 | 0.002651967 | 0.012653365 | -1.011651079 | 5.47E-07 | 1.29E-05 | -0.752530703 | 0.000134227 | 0.000878395 |
| Q9BT09 | CNPY3 | -0.226102958 | 0.071961148 | 0.183334544 | -0.726628135 | 0.000707974 | 0.004241793 | -1.467741695 | 3.09E-10 | 1.71E-08 |
| P53602 | MVD | 0.454946009 | 0.028004781 | 0.086542383 | 1.169457021 | 7.03E-06 | 0.000104876 | 1.440686558 | 5.76E-06 | 6.34E-05 |
| P41218 | MNDA | 0.276706933 | 0.004450856 | 0.019398912 | -2.245823665 | 1.16E-07 | 3.33E-06 | -2.104372753 | 6.30E-07 | 1.02E-05 |
| P32119 | PRDX2 | 2.21E-15 | 0.999999925 | 1 | 0.767045471 | 3.15E-06 | 5.36E-05 | 0.612055428 | 0.000112359 | 0.000762165 |
| O75874 | IDH1 | 0.110857078 | 0.215795019 | 0.435953689 | -1.397508356 | 1.58E-09 | 7.76E-08 | -1.47942404 | 3.65E-09 | 1.35E-07 |
| P50749 | RASSF2 | 0 | 1 | 1 | -0.464678827 | 0.004310399 | 0.018343067 | -0.809828807 | 2.66E-05 | 0.000227341 |
| P61626 | LYZ | -0.835164001 | 0.006267064 | 0.02578491 | -3.624359727 | 5.47E-12 | 7.31E-10 | -4.777228714 | 2.74E-14 | 6.47E-12 |
| P51659 | HSD17B4 | -4.62E-16 | 0.999999967 | 1 | -0.468762362 | 0.014521632 | 0.048017283 | -0.84611619 | 6.82E-06 | 7.17E-05 |
| Q13492;Q13492-2;Q13492-5 | PICALM | 0 | 1 | 1 | -0.86900336 | 0.000793625 | 0.00465549 | -1.103518596 | 2.61E-05 | 0.000224448 |
| Q13363;Q13363-2 | CTBP1 | 0.834772637 | 6.63E-06 | 8.06E-05 | 0.518620008 | 0.00085911 | 0.00497716 | 1.129449474 | 6.85E-07 | 1.10E-05 |
| P45954;P45954-2 | ACADSB | 0 | 1 | 1 | 0.800719325 | 1.20E-07 | 3.40E-06 | 0.620714977 | 0.000468709 | 0.002485117 |

| | | | | | | | | | | |
|---|---|---|---|---|---|---|---|---|---|---|
| P35443 | THBS4 | 0 | 1 | 1 | 1.759746687 | 0.001188145 | 0.006494264 | 1.85227806 | 0.000536552 | 0.002782358 |
| Q96IU4 | ABHD14B | 0.349376811 | 0.031070841 | 0.093717525 | 0.493358263 | 0.000183663 | 0.001426565 | 0.725884212 | 0.000223486 | 0.00131922 |
| Q5SSJ5 | HP1BP3 | 0.024768069 | 0.684637877 | 1 | 0.827513708 | 3.00E-06 | 5.16E-05 | 0.691228087 | 0.00039163 | 0.002117196 |
| P00738 | HP | 0.478431182 | 0.000263223 | 0.001856246 | -1.523559765 | 7.19E-07 | 1.57E-05 | -1.058084386 | 0.000159177 | 0.001006182 |
| O95218;O95218-2 | ZRANB2 | 0.491863593 | 0.023053508 | 0.073629229 | 0.89319344 | 9.31E-06 | 0.000129276 | 1.002065593 | 0.000133503 | 0.000876195 |
| Q9UFN0 | NIPSNAP3A | 0.332888911 | 0.020589194 | 0.067282203 | -0.755756656 | 0.000446262 | 0.002958199 | -0.684017569 | 0.00047773 | 0.002522648 |
| Q96HE7 | ERO1A | 0.231824196 | 0.01382416 | 0.048743989 | -0.571931958 | 0.001244216 | 0.006717292 | -0.607134818 | 0.001243104 | 0.005568248 |
| Q7L5N7 | LPCAT2 | -3.60E-19 | 1 | 1 | -1.693573598 | 2.05E-05 | 0.000242809 | -1.936336256 | 1.68E-05 | 0.000152333 |
| P05164;P05164-2;P05164-3 | MPO | -0.217624862 | 0.246307303 | 0.480885686 | -3.512848337 | 3.76E-12 | 5.55E-10 | -4.4765151 | 3.28E-15 | 9.48E-13 |
| P21281 | ATP6V1B2 | 0.414619881 | 0.00253603 | 0.012149512 | -1.050894294 | 1.22E-05 | 0.000163667 | -0.690407885 | 0.00344843 | 0.012743986 |
| P20645 | M6PR | 9.76E-18 | 0.999999999 | 1 | -0.707367393 | 0.010608885 | 0.037702554 | -1.182401358 | 0.000367706 | 0.002011156 |
| Q15942 | ZYX | 1.27455195 | 4.87E-05 | 0.000460554 | 1.288685666 | 4.43E-06 | 7.13E-05 | 1.80068391 | 1.61E-06 | 2.22E-05 |
| O75563 | SKAP2 | 1.04E-17 | 0.999999996 | 1 | -0.725807097 | 9.64E-05 | 0.000852629 | -1.007012306 | 1.63E-06 | 2.23E-05 |
| Q8NFW8 | CMAS | 0.232003718 | 0.024219187 | 0.076726734 | 0.879822646 | 4.45E-09 | 1.81E-07 | 0.93330902 | 1.40E-06 | 1.96E-05 |
| O00479 | HMGN4 | 0.493832762 | 0.067102833 | 0.173846134 | 1.891018157 | 1.06E-06 | 2.22E-05 | 2.121481704 | 1.89E-08 | 5.50E-07 |
| P53999 | SUB1 | 0.480304477 | 0.016450401 | 0.055826867 | 0.625773001 | 6.29E-05 | 0.000605808 | 1.361300175 | 2.02E-08 | 5.71E-07 |
| Q5T1M5 | FKBP15 | 1.40E-17 | 0.999999994 | 1 | -0.694609324 | 3.34E-06 | 5.64E-05 | -0.842268319 | 6.33E-05 | 0.000473733 |
| P11177;P11177-2 | PDHB | 0.644547719 | 0.000480964 | 0.003083419 | 0.605350016 | 0.000150421 | 0.001215503 | 0.900615411 | 3.09E-07 | 5.46E-06 |
| P42765 | ACAA2 | 0.782282202 | 0.000121762 | 0.000972011 | 0.857185175 | 2.98E-08 | 9.94E-07 | 1.438997854 | 8.99E-11 | 6.87E-09 |
| P30040 | ERP29 | 0.208942749 | 0.042504267 | 0.120668313 | -1.002914353 | 3.71E-08 | 1.20E-06 | -0.852637381 | 9.11E-06 | 9.17E-05 |
| P49902 | NT5C2 | -1.33E-18 | 0.999999999 | 1 | -0.683581115 | 0.001505489 | 0.00792744 | -1.184941335 | 1.29E-06 | 1.89E-05 |
| P04792 | HSPB1 | 4.66E-16 | 0.999999976 | 1 | 0.802103855 | 0.006020305 | 0.024359214 | 0.976171138 | 0.001757148 | 0.007386846 |
| O75083 | WDR1 | 0.446103776 | 0.001315404 | 0.007069701 | 0.505929202 | 0.000272385 | 0.001983814 | 0.761927767 | 4.37E-06 | 5.02E-05 |
| Q6NYC8 | PPP1R18 | 0.801024973 | 0.004242358 | 0.018651565 | 0.877181112 | 7.74E-05 | 0.000709338 | 1.181119202 | 0.000601173 | 0.003086654 |
| Q15208 | STK38 | 0.134099761 | 0.113558318 | 0.262217832 | 0.84136334 | 1.87E-11 | 1.87E-09 | 0.754152536 | 7.07E-10 | 3.28E-08 |
| P07858 | CTSB | 0 | 1 | 1 | -1.995399677 | 1.47E-08 | 5.21E-07 | -2.367745335 | 9.28E-11 | 6.89E-09 |
| Q9NZK5 | ADA2 | -0.28200125 | 0.056919937 | 0.153205875 | -0.932486503 | 1.54E-05 | 0.000193986 | -1.546158839 | 3.19E-07 | 5.56E-06 |
| P50395 | GDI2 | 0.129684757 | 0.137900167 | 0.306773495 | -0.588102562 | 0.001119288 | 0.006205204 | -0.603986147 | 0.001652457 | 0.007014844 |
| O15427 | SLC16A3 | -0.709316162 | 0.008339268 | 0.032526837 | -0.861656714 | 0.000405586 | 0.002780598 | -2.423761291 | 2.28E-10 | 1.38E-08 |
| P00492 | HPRT1 | -1.09E-15 | 0.999999932 | 1 | -0.929985025 | 5.85E-08 | 1.84E-06 | -1.000793672 | 1.44E-07 | 2.99E-06 |
| Q96CX2 | KCTD12 | 7.44E-16 | 0.999999958 | 1 | -2.350328243 | 1.11E-07 | 3.22E-06 | -2.725962233 | 6.31E-08 | 1.51E-06 |
| Q8TDZ2;Q8TDZ2-4 | MICAL1 | 1.84E-15 | 0.999999914 | 1 | -0.523743886 | 0.002829891 | 0.013247103 | -1.185552121 | 9.05E-10 | 4.13E-08 |
| Q9UBR2 | CTSZ | -2.66E-05 | 0.98795026 | 1 | -1.70971459 | 2.64E-10 | 1.76E-08 | -2.114373602 | 1.41E-11 | 1.53E-09 |
| P22392-2 | NME2 | 4.96E-17 | 0.999999986 | 1 | -0.634819827 | 8.63E-07 | 1.83E-05 | -0.59791254 | 0.000529227 | 0.00276497 |
| P46109 | CRKL | 0.856169768 | 3.84E-12 | 3.15E-10 | 0.541432063 | 6.85E-07 | 1.52E-05 | 0.987376747 | 1.45E-12 | 2.06E-10 |
| P0DOX7 | sp\|P0DOX7\|IGK_HUMAN | 0.320389441 | 0.170930211 | 0.363163961 | 1.058089921 | 8.13E-06 | 0.000117488 | 1.184846808 | 4.06E-05 | 0.000324862 |
| Q7L591 | DOK3 | 0 | 1 | 1 | -0.823067814 | 0.002300946 | 0.011181718 | -1.035866272 | 0.000169856 | 0.00105571 |
| O43776 | NARS1 | 4.70E-17 | 0.999999986 | 1 | -0.56167382 | 2.95E-05 | 0.000331419 | -0.682606987 | 1.28E-05 | 0.00012108 |
| P04839 | CYBB | 0.420532354 | 0.20365098 | 0.418213951 | -3.61896285 | 4.95E-10 | 2.88E-08 | -3.24640089 | 1.35E-06 | 1.93E-05 |
| Q6VY07 | PACS1 | 0.128564889 | 0.141965771 | 0.313248628 | 0.721236231 | 5.17E-12 | 7.25E-10 | 0.694197632 | 0.006768376 | 0.022543898 |
| P17693;P17693-5 | HLA-G | 2.70E-16 | 0.999999977 | 1 | 1.011440251 | 0.008006768 | 0.030752983 | 1.193134919 | 0.001116508 | 0.00516908 |
| P32456 | GBP2 | 1.211896833 | 6.61E-11 | 3.53E-09 | -0.618612314 | 0.001163042 | 0.00636947 | 0.692290318 | 0.008581718 | 0.027593198 |

| | | | | | | | | | | |
|---|---|---|---|---|---|---|---|---|---|---|
| P20839;P20839-3;P20839-5;P20839-6;P20839-7 | IMPDH1 | 0.690099219 | 1.66E-12 | 1.50E-10 | -0.928012643 | 5.12E-05 | 0.000513146 | -0.691221295 | 0.00110252 | 0.005124057 |
| P25774 | CTSS | 0.255921583 | 0.032696631 | 0.097867845 | -1.071958953 | 9.23E-05 | 0.000824654 | -1.052763938 | 9.74E-05 | 0.000687935 |
| Q6P4A8 | PLBD1 | -7.33E-16 | 0.99999996 | 1 | -1.867818358 | 2.31E-08 | 7.98E-07 | -2.038914584 | 4.87E-06 | 5.55E-05 |
| P35244 | RPA3 | 0.422414534 | 0.09112177 | 0.220973427 | 0.883970836 | 0.001488848 | 0.007862015 | 1.244645312 | 1.15E-05 | 0.000111331 |
| P43405 | SYK | -0.215381582 | 0.057012953 | 0.153224124 | -0.895050426 | 6.59E-07 | 1.48E-05 | -1.414902841 | 3.91E-10 | 2.07E-08 |
| Q9UGI8 | TES | 0.31163229 | 0.110157078 | 0.255872107 | 0.899587528 | 1.42E-06 | 2.81E-05 | 1.047613789 | 5.37E-06 | 6.06E-05 |
| Q9UKD2 | MRTO4 | -2.04E-16 | 0.999999988 | 1 | 1.141007567 | 0.000657899 | 0.004045503 | 1.155804534 | 0.000119278 | 0.00080281 |
| Q06187 | BTK | 3.20E-16 | 0.999999959 | 1 | -0.824809805 | 0.005497992 | 0.022604649 | -1.144898098 | 0.002144523 | 0.0086255 |
| Q14012 | CAMK1 | 1.497122263 | 1.31E-16 | 2.88E-14 | -1.256150783 | 1.64E-07 | 4.51E-06 | 0.961063164 | 4.88E-05 | 0.000378767 |
| Q9H4M9 | EHD1 | 0.190160805 | 0.009898377 | 0.037528686 | 0.707576482 | 9.13E-10 | 4.74E-08 | 0.711929778 | 5.08E-10 | 2.59E-08 |
| Q5TEC6 | NA | 0.208321393 | 0.293872656 | 0.556196986 | 0.865126731 | 0.001957413 | 0.009848491 | 1.038473675 | 0.000133554 | 0.000876195 |
| Q15555-5 | MAPRE2 | 0.35171325 | 0.063165247 | 0.165714777 | 0.623975428 | 0.000915247 | 0.005248167 | 0.702922006 | 0.003536038 | 0.013012219 |
| P29218;P29218-3 | IMPA1 | 0.199214903 | 0.055981438 | 0.151373121 | 0.844469803 | 1.53E-09 | 7.66E-08 | 0.841776693 | 5.90E-06 | 6.47E-05 |
| O75436 | VPS26A | 0.348266872 | 0.000854173 | 0.00501134 | -0.848731412 | 1.17E-05 | 0.000161395 | -0.638067111 | 0.004250219 | 0.015293725 |
| P83111 | LACTB | 0.46582303 | 0.006742523 | 0.02738956 | -0.822766452 | 0.003241021 | 0.01482516 | -0.91143001 | 0.000279862 | 0.001587517 |
| P30536 | TSPO | 1.08E-15 | 0.999999978 | 1 | -1.38063226 | 0.000650588 | 0.004018166 | -1.48588751 | 0.001721258 | 0.00725946 |
| Q9BQS8;Q9BQS8-4 | FYCO1 | 5.71E-18 | 0.999999997 | 1 | 0.58486668 | 0.004273852 | 0.01826811 | 1.137425554 | 7.04E-07 | 1.12E-05 |
| Q9H0A8 | COMMD4 | 0.309290375 | 0.037036876 | 0.107394758 | -0.610561854 | 0.000117836 | 0.000986306 | -0.509155553 | 0.007096744 | 0.023427372 |
| P09972 | ALDOC | 0.274140734 | 0.039002321 | 0.112354724 | 0.680678695 | 3.17E-05 | 0.000349686 | 0.902455773 | 5.45E-06 | 6.11E-05 |
| Q9NTX5;Q9NTX5-2;Q9NTX5-6 | ECHDC1 | 0.065647303 | 0.382584069 | 0.685811605 | -0.512622429 | 6.21E-06 | 9.41E-05 | -0.622230892 | 5.97E-05 | 0.000449318 |
| P01859;P01859-1 | IGHG2 | 0.514798731 | 0.013953967 | 0.049054524 | 1.159860837 | 1.07E-07 | 3.12E-06 | 1.456448671 | 1.63E-08 | 4.97E-07 |
| O43852;O43852-3 | CALU | 0.917389444 | 0.001153135 | 0.006343144 | 0.70487688 | 0.000775285 | 0.004567012 | 1.39511643 | 2.85E-06 | 3.51E-05 |
| Q13435 | SF3B2 | 0.826023028 | 5.47E-07 | 8.93E-06 | 0.468885497 | 0.000938089 | 0.005351645 | 0.953279803 | 9.99E-07 | 1.51E-05 |
| Q53H82 | LACTB2 | 0 | 1 | 1 | -1.030932351 | 0.001467482 | 0.007773867 | -0.924567649 | 0.001636387 | 0.006957992 |
| Q9UK45 | LSM7 | 0.043092149 | 0.599948365 | 0.99315396 | 0.583976546 | 0.001121982 | 0.006205204 | 0.535379808 | 0.003105114 | 0.0117084 |
| P58876 | H2BC5 | 0.502661661 | 0.002956099 | 0.013805568 | 0.583014872 | 0.006889797 | 0.027171575 | 0.704300365 | 1.04E-05 | 0.000102244 |
| O43670;O43670-2;O43670-4 | ZNF207 | 0.572722202 | 0.05019126 | 0.138489621 | 1.422517337 | 8.04E-09 | 3.01E-07 | 1.207878994 | 0.000314035 | 0.001773617 |
| Q5VVQ6;Q5VVQ6-2 | YOD1 | 0 | 1 | 1 | 0.817794533 | 0.001054677 | 0.005914627 | 0.638796674 | 0.010672714 | 0.032930774 |
| O95834;O95834-2;O95834-3 | EML2 | 0.025499477 | 0.649775747 | 1 | 0.679039937 | 1.38E-05 | 0.0001793 | 0.600164371 | 0.000148289 | 0.000958344 |
| P08236;P08236-2 | GUSB | 0.225958499 | 0.022299363 | 0.071740468 | 0.564616289 | 5.54E-05 | 0.000551254 | 0.665752969 | 1.00E-06 | 1.51E-05 |
| O00182;O00182-2;O00182-3;O00182-6 | LGALS9 | 1.165906773 | 6.91E-10 | 2.80E-08 | -1.372330798 | 4.77E-08 | 1.52E-06 | -0.618158707 | 0.00902992 | 0.028714481 |
| Q30154 | HLA-DRB5 | 0.641862194 | 0.056381282 | 0.152220826 | -2.34358523 | 6.12E-07 | 1.40E-05 | -2.079006578 | 4.59E-06 | 5.26E-05 |
| Q93091 | RNASE6 | -0.61345481 | 0.000154987 | 0.00118801 | -1.912495493 | 2.17E-07 | 5.58E-06 | -2.371764825 | 1.31E-07 | 2.79E-06 |
| Q8TCT9;Q8TCT9-5 | HM13 | 0 | 1 | 1 | -1.071086434 | 0.002151857 | 0.010585624 | -1.069571814 | 0.002250449 | 0.008994872 |
| Q96QR8 | PURB | 0.373472917 | 0.020673685 | 0.067495752 | 0.510620887 | 0.001469383 | 0.007773867 | 0.585041892 | 0.000634163 | 0.00321788 |
| Q9BUH6 | PAXX | 0.901010168 | 0.000480942 | 0.003083419 | 1.558370617 | 1.21E-13 | 3.09E-11 | 2.419797765 | 2.19E-09 | 8.88E-08 |
| P17480;P17480-2 | UBTF | -3.50E-18 | 0.999999998 | 1 | 1.098088735 | 8.88E-08 | 2.65E-06 | 0.806681816 | 0.000120886 | 0.000809437 |

| | | | | | | | | | | |
|---|---|---|---|---|---|---|---|---|---|---|
| Q8TBX8;Q8TBX8-3 | PIP4K2C | 6.84E-18 | 0.999999999 | 1 | 1.28737655 | 4.13E-10 | 2.51E-08 | 1.047269307 | 3.85E-08 | 1.02E-06 |
| P20933 | AGA | -4.57E-17 | 0.999999992 | 1 | 1.253960943 | 1.55E-11 | 1.74E-09 | 0.940211276 | 2.82E-07 | 5.12E-06 |
| Q05315 | CLC | 0.684392794 | 0.07751609 | 0.194396681 | 2.285828334 | 2.91E-06 | 5.07E-05 | 3.009126969 | 2.74E-07 | 5.04E-06 |
| Q2M2I8;Q2M2I8-2 | AAK1 | 0 | 1 | 1 | 0.711016879 | 0.000284627 | 0.002062258 | 0.724550056 | 0.001405379 | 0.006136796 |
| Q15637;Q15637-2;Q15637-3;Q15637-4;Q15637-6 | SF1 | 0.230067305 | 0.22335028 | 0.44670056 | 0.498347462 | 0.000129408 | 0.001073554 | 1.026345732 | 9.57E-05 | 0.000679444 |
| P48960 | ADGRE5 | 1.17E-17 | 0.999999996 | 1 | -0.893052348 | 0.000384716 | 0.002663568 | -0.607694466 | 0.016921702 | 0.04831053 |
| A0A075B6P5;A0A087WW87;P01614;P01615 | IGKV2-28;IGKV2-40;IGKV2D-40;IGKV2D-28 | 0 | 1 | 1 | 1.080332829 | 4.30E-06 | 7.01E-05 | 1.045782022 | 0.000492628 | 0.00258555 |
| Q9NY12;Q9NY12-2 | GAR1 | 0.821543197 | 5.56E-05 | 0.000512212 | 0.612591766 | 0.001219082 | 0.006624621 | 1.161462363 | 4.35E-05 | 0.000344475 |
| Q92572 | AP3S1 | 0 | 1 | 1 | -0.502166384 | 0.003641414 | 0.016104928 | -0.774190673 | 0.000163525 | 0.001026179 |
| P49419;P49419-2 | ALDH7A1 | -3.56E-16 | 0.999999967 | 1 | -0.611921342 | 0.009260854 | 0.034134149 | -1.078547967 | 0.000118224 | 0.000797779 |
| P22830;P22830-2 | FECH | 0.207095679 | 0.177192749 | 0.374569325 | 1.009191233 | 6.08E-07 | 1.40E-05 | 1.005365195 | 3.37E-05 | 0.00027915 |
| O00602 | FCN1 | -0.51988107 | 0.067198619 | 0.173966469 | -3.877856884 | 7.47E-08 | 2.25E-06 | -3.765314423 | 1.21E-06 | 1.80E-05 |
| Q9BV40 | VAMP8 | -7.44E-19 | 0.999999999 | 1 | -0.942467802 | 0.004194505 | 0.01809445 | -1.207600591 | 0.000245573 | 0.001424106 |
| Q14847 | LASP1 | 1.979487162 | 5.96E-08 | 1.33E-06 | 1.590209025 | 3.22E-11 | 2.92E-09 | 2.906340808 | 6.51E-10 | 3.13E-08 |
| P56192 | MARS1 | 0 | 1 | 1 | -0.549327166 | 2.01E-06 | 3.76E-05 | -0.755986837 | 7.16E-07 | 1.13E-05 |
| Q6RW13 | AGTRAP | -0.117564639 | 0.482509206 | 0.828299641 | -0.692618941 | 0.005272169 | 0.021864371 | -2.105909587 | 2.37E-06 | 3.07E-05 |
| Q9UBF2 | COPG2 | 3.14E-16 | 0.999999969 | 1 | -0.850041762 | 7.11E-05 | 0.000662361 | -1.051059498 | 0.000100556 | 0.000706067 |
| O00151 | PDLIM1 | 0.424824744 | 0.225881006 | 0.450484405 | 1.178097443 | 0.002237836 | 0.01093187 | 2.004588267 | 1.58E-05 | 0.000145649 |
| P09668 | CTSH | 0.582742229 | 0.000881574 | 0.005120973 | -1.689500974 | 1.55E-06 | 3.01E-05 | -1.452460139 | 7.73E-06 | 7.97E-05 |
| Q6UW68 | TMEM205 | 5.98E-17 | 0.999999997 | 1 | -1.051693301 | 0.003525664 | 0.015692002 | -1.3487393 | 0.001556953 | 0.006685889 |
| Q8IYJ3 | SYTL1 | 0.405017975 | 0.000191528 | 0.001412823 | 0.989097548 | 1.93E-09 | 9.35E-08 | 0.966763539 | 1.42E-07 | 2.97E-06 |
| O95400 | CD2BP2 | 0.302193069 | 0.035384568 | 0.103798657 | 0.671998888 | 2.36E-06 | 4.32E-05 | 0.630580975 | 0.001024876 | 0.004806188 |
| Q9BT23 | LIMD2 | 9.20E-19 | 0.999999999 | 1 | 0.932458634 | 1.53E-09 | 7.66E-08 | 0.721524315 | 0.000436296 | 0.002332296 |
| P35813;P35813-3 | PPM1A | 0.136889722 | 0.207335883 | 0.423561022 | 0.648542003 | 1.20E-05 | 0.000163147 | 0.694579392 | 0.000396135 | 0.002130762 |
| Q9HC16 | APOBEC3G | 0.329732118 | 0.100494083 | 0.238293299 | 1.813604904 | 3.41E-11 | 2.98E-09 | 2.130712495 | 5.09E-09 | 1.74E-07 |
| O94925 | GLS | 0.628566406 | 0.003974522 | 0.017694652 | 1.132337133 | 6.58E-09 | 2.53E-07 | 1.493643294 | 3.09E-07 | 5.46E-06 |
| Q04941 | PLP2 | -0.803634774 | 0.019159581 | 0.063552854 | -1.165661162 | 0.000243631 | 0.001826584 | -2.330212141 | 7.73E-08 | 1.79E-06 |
| P27816 | MAP4 | 0.278902501 | 0.136620067 | 0.304502121 | 0.643735076 | 0.006878059 | 0.027163489 | 1.274725471 | 2.98E-06 | 3.65E-05 |
| Q9BVG4 | PBDC1 | 0.220766618 | 0.087673998 | 0.214233205 | 0.531611384 | 2.70E-06 | 4.77E-05 | 0.72053968 | 1.26E-05 | 0.00012077 |
| Q8TCE6 | DENND10 | 0.299036189 | 0.00746699 | 0.029783491 | -0.750152393 | 7.96E-06 | 0.000115675 | -0.753019674 | 3.87E-06 | 4.54E-05 |
| P17174 | GOT1 | 0.119997326 | 0.341470174 | 0.623523477 | 0.636190346 | 0.00140747 | 0.007503265 | 0.542702397 | 0.012316125 | 0.03711983 |
| Q9HAU5 | UPF2 | 0.302426416 | 0.05724124 | 0.153718669 | 0.543483995 | 0.001959864 | 0.009848491 | 0.545479627 | 0.006065821 | 0.020519534 |
| O15347 | HMGB3 | -3.44E-17 | 0.999999989 | 1 | -1.123475474 | 2.55E-05 | 0.000294424 | -1.001349054 | 0.000206528 | 0.001247814 |
| Q9NTM9 | CUTC | 2.75E-15 | 0.999999923 | 1 | 1.046191082 | 4.20E-10 | 2.51E-08 | 0.869657693 | 3.41E-05 | 0.000281051 |
| Q5VSL9;Q5VSL9-2 | STRIP1 | 0.41180855 | 0.00179596 | 0.009085442 | 0.51239881 | 0.000239813 | 0.001812201 | 0.614920288 | 4.53E-05 | 0.000355936 |
| Q9Y2I7 | PIKFYVE | 1.805488733 | 6.29E-08 | 1.39E-06 | -0.480008743 | 0.009901826 | 0.035825445 | 0.847127562 | 0.001146808 | 0.005245437 |
| P17612 | PRKACA | 0.702155369 | 5.82E-05 | 0.000531755 | -1.184921703 | 7.03E-08 | 2.17E-06 | -0.819989972 | 0.002038991 | 0.008289981 |
| P98175;P98175-2;P98175-5 | RBM10 | 0.282177702 | 0.040690214 | 0.116645281 | 0.601314486 | 0.000828366 | 0.004824332 | 0.997823881 | 0.00180824 | 0.007540356 |
| Q5VTL8 | PRPF38B | 0.363410624 | 0.001901871 | 0.009541143 | 0.497042583 | 2.33E-05 | 0.000272989 | 0.611261235 | 9.69E-07 | 1.47E-05 |

| | | | | | | | | | | |
|---|---|---|---|---|---|---|---|---|---|---|
| Q8TBC4;Q8TBC4-2 | UBA3 | 0.137654155 | 0.172797084 | 0.366595979 | 0.774841247 | 3.23E-08 | 1.06E-06 | 0.758087111 | 5.48E-06 | 6.11E-05 |
| Q10567-3 | AP1B1 | 0.013421588 | 0.780298011 | 1 | -0.689660808 | 4.93E-05 | 0.000495909 | -0.730764472 | 0.000357703 | 0.001964717 |
| O94905 | ERLIN2 | -3.80E-19 | 1 | 1 | -1.139966305 | 0.000645073 | 0.004001739 | -2.233045683 | 1.64E-06 | 2.23E-05 |
| Q8WVT3 | TRAPPC12 | 6.89E-16 | 0.999999967 | 1 | -0.65221463 | 0.002138791 | 0.010572328 | -0.949678219 | 1.38E-05 | 0.000129235 |
| Q9HBH0 | RHOF | 0 | 1 | 1 | 1.17605695 | 0.000134678 | 0.0011042 | 1.188275995 | 3.84E-06 | 4.53E-05 |
| Q5R3I4 | TTC38 | 0.370161979 | 0.076001012 | 0.191140918 | 1.254675098 | 3.50E-06 | 5.87E-05 | 1.106755593 | 0.00091763 | 0.004342448 |
| Q05209 | PTPN12 | 0.341905629 | 0.013243039 | 0.047214311 | 0.846701558 | 1.21E-05 | 0.000163468 | 0.643210994 | 0.001854327 | 0.007683479 |
| Q15904 | ATP6AP1 | 3.64E-16 | 0.99999999 | 1 | -1.104694833 | 0.000392269 | 0.002709169 | -1.170783095 | 0.003667812 | 0.013402214 |
| O95721 | SNAP29 | 0.842985957 | 0.004920671 | 0.021158886 | 0.635407195 | 0.006093194 | 0.024547868 | 0.958721882 | 0.00115022 | 0.005251794 |
| P53634 | CTSC | 0.368904846 | 0.005770797 | 0.023938624 | 1.773479307 | 4.39E-16 | 2.46E-13 | 2.136217223 | 1.14E-18 | 1.48E-15 |
| O14618 | CCS | 0.444072049 | 0.034762649 | 0.102571632 | 0.646926163 | 4.11E-05 | 0.000426245 | 0.563965146 | 0.007325205 | 0.023998592 |
| P30533 | LRPAP1 | -0.246859629 | 0.058625578 | 0.156517776 | -0.639162073 | 0.00082929 | 0.004824332 | -1.310707035 | 2.27E-09 | 8.93E-08 |
| Q8IV04 | TBC1D10C | 0.37087441 | 0.065329506 | 0.170252651 | 1.389867419 | 6.45E-09 | 2.51E-07 | 1.509432395 | 4.57E-08 | 1.18E-06 |
| Q9UEW8;Q9UEW8-2 | STK39 | -0.381377484 | 0.076630723 | 0.192313117 | 1.116504426 | 5.04E-06 | 7.93E-05 | 0.757845977 | 0.003698598 | 0.013495728 |
| Q99733;Q99733-2 | NAP1L4 | 0.95167726 | 0.00025526 | 0.001807326 | 0.667775142 | 0.000107377 | 0.000925392 | 1.520421049 | 0.000124868 | 0.000833952 |
| Q9NYB0 | TERF2IP | 0.153075466 | 0.229682425 | 0.455489444 | 0.628502314 | 1.74E-12 | 3.25E-10 | 0.598775933 | 0.000119881 | 0.000804784 |
| P24158 | PRTN3 | -0.93138002 | 0.007755905 | 0.030692842 | -3.457445174 | 1.73E-11 | 1.85E-09 | -5.075542177 | 6.14E-13 | 1.06E-10 |
| O43399;O43399-5;O43399-7 | TPD52L2 | 1.345674079 | 2.05E-06 | 2.88E-05 | -0.823435075 | 3.72E-06 | 6.17E-05 | 0.52808817 | 0.010819733 | 0.033305292 |
| Q9C0A0;Q9C0A0-2 | CNTNAP4 | 0.687438316 | 0.111694251 | 0.258628247 | 2.297022812 | 1.77E-06 | 3.40E-05 | 3.328673138 | 3.53E-11 | 3.31E-09 |
| Q8WUD4 | CCDC12 | 0.796273207 | 0.005003543 | 0.021462885 | 0.774075265 | 0.000340297 | 0.002397467 | 1.436558666 | 1.48E-05 | 0.000138753 |
| A0A0A0MRZ8;P04433 | IGKV3D-11;IGKV3-11 | 0.710797197 | 0.001990393 | 0.009926629 | 1.294328993 | 7.14E-08 | 2.17E-06 | 1.747674029 | 4.97E-08 | 1.25E-06 |
| P01619 | IGKV3-20 | 0.57201441 | 0.027984788 | 0.086542383 | 0.966044984 | 0.000236842 | 0.001794875 | 1.364708131 | 2.21E-05 | 0.000194188 |
| A0A075B6H7;A0A0C4DH55;P01624 | IGKV3-7;IGKV3D-7;IGKV3-15 | 0.87304851 | 0.000742962 | 0.004432629 | 1.321816841 | 3.15E-06 | 5.36E-05 | 2.108791169 | 6.43E-12 | 7.95E-10 |
| P50750;P50750-2 | CDK9 | 0.338092039 | 0.029798138 | 0.090654214 | 0.644732354 | 0.011707534 | 0.040528302 | 0.765684921 | 0.000487117 | 0.002561801 |
| P30043 | BLVRB | 0.258219076 | 0.024430709 | 0.077321551 | -1.759497667 | 6.59E-07 | 1.48E-05 | -1.717614149 | 1.91E-06 | 2.54E-05 |
| Q14118 | DAG1 | 0.139262304 | 0.511097691 | 0.867242761 | 1.545448991 | 1.52E-05 | 0.000192793 | 1.843515994 | 3.67E-05 | 0.000298661 |
| Q9BPW8 | NIPSNAP1 | 0 | 1 | 1 | 0.822211587 | 3.43E-05 | 0.000374033 | 0.78603927 | 0.000838267 | 0.004010716 |
| P16150 | SPN | 3.55E-16 | 0.999999972 | 1 | 1.382121293 | 6.31E-05 | 0.000605808 | 1.728552134 | 1.39E-06 | 1.96E-05 |
| P04908;Q7L7L0 | H2AC4;H2AW | 1.100104835 | 0.006296211 | 0.025874638 | 1.537944339 | 0.000114409 | 0.00096627 | 2.531137303 | 1.91E-08 | 5.50E-07 |
| P15531;P15531-2 | NME1 | -0.238508125 | 0.21668141 | 0.436675949 | -0.621584244 | 0.004251855 | 0.018208161 | -1.393147032 | 4.32E-06 | 5.01E-05 |
| P05090 | APOD | 2.64E-16 | 0.999999982 | 1 | 1.057906977 | 0.007678587 | 0.029820992 | 1.263860399 | 0.009703188 | 0.030408785 |
| Q9NX24 | NHP2 | 0.429049315 | 0.012092934 | 0.043913167 | 0.481149931 | 0.000173552 | 0.001374688 | 0.852601802 | 6.64E-06 | 7.04E-05 |
| Q15427 | SF3B4 | 0.766743389 | 0.029351425 | 0.089682083 | 0.850429419 | 0.000177909 | 0.001401281 | 1.427908472 | 0.000341558 | 0.001896085 |
| O94966;O94966-3;O94966-4;O94966-5;O94966-6;O94966-7 | USP19 | 0 | 1 | 1 | 1.628075147 | 0.003106377 | 0.01430763 | 1.452302039 | 0.00116573 | 0.005313275 |
| Q9UDT6;Q9UDT6-2 | CLIP2 | 0 | 1 | 1 | 1.956883039 | 6.18E-05 | 0.000599317 | 1.992888677 | 0.000228717 | 0.001341323 |
| P08246 | ELANE | -1.52499677 | 3.18E-05 | 0.000311942 | -3.722049913 | 3.15E-10 | 2.01E-08 | -5.706215737 | 3.42E-14 | 7.40E-12 |
| Q8ND56;Q8ND56-2;Q8ND56-3 | LSM14A | 4.70E-16 | 0.999999969 | 1 | 0.831788791 | 2.88E-06 | 5.05E-05 | 0.560240693 | 0.007293741 | 0.023925681 |

| | | | | | | | | | | |
|---|---|---|---|---|---|---|---|---|---|---|
| Q96M27;Q96M27-2;Q96M27-3 | PRRC1 | 0.336633483 | 0.001039069 | 0.005834011 | 0.533008085 | 0.002141613 | 0.010572328 | 0.520610077 | 0.009250168 | 0.029271542 |
| Q13315 | ATM | 0.276289486 | 0.143462942 | 0.315761756 | 0.6969864 | 0.000249894 | 0.001863571 | 0.638879749 | 0.013457176 | 0.039594272 |
| Q14644 | RASA3 | 0 | 1 | 1 | 1.241604218 | 1.23E-07 | 3.46E-06 | 0.88688024 | 0.000844541 | 0.004033304 |
| P51668 | UBE2D1 | 0 | 1 | 1 | -1.04293535 | 7.66E-05 | 0.000704136 | -1.456663329 | 6.02E-08 | 1.46E-06 |
| Q96F07-2 | CYFIP2 | 0.714129186 | 3.77E-05 | 0.000364393 | 0.995949947 | 2.54E-09 | 1.15E-07 | 1.425290807 | 1.43E-10 | 9.24E-09 |
| Q16799 | RTN1 | -0.607224724 | 0.013014148 | 0.046539438 | -1.209784434 | 0.000162343 | 0.0013006 | -2.198895976 | 2.69E-08 | 7.29E-07 |
| Q16270;Q16270-2 | IGFBP7 | 0.027031711 | 0.698367887 | 1 | 0.940854023 | 0.000107588 | 0.000925392 | 0.913943797 | 2.72E-05 | 0.000230184 |
| P22033 | MMUT | 0.538190205 | 0.022498566 | 0.072315355 | 0.843406112 | 0.000242427 | 0.00182243 | 1.212061756 | 8.52E-06 | 8.68E-05 |
| Q96I59 | NARS2 | 0.518785078 | 0.084940781 | 0.209148879 | -0.98272328 | 0.002015072 | 0.010071766 | -0.729700966 | 0.001547525 | 0.006656409 |
| O95197-3 | RTN3 | -0.356399174 | 0.021663957 | 0.070144272 | -1.209394937 | 5.36E-06 | 8.40E-05 | -2.141291994 | 1.03E-08 | 3.28E-07 |
| P01834 | IGKC | 0.597652311 | 0.046988008 | 0.131491837 | 0.634754635 | 0.009671029 | 0.035217617 | 1.090831873 | 0.003103562 | 0.0117084 |
| Q8IWA5;Q8IWA5-3 | SLC44A2 | 0.623901434 | 0.009895657 | 0.037528686 | 0.724642686 | 0.009223622 | 0.034079683 | 1.145412818 | 0.000150011 | 0.000967066 |
| O14966 | RAB29 | 0.474590426 | 0.024763247 | 0.078239435 | 1.14338881 | 0.001905051 | 0.009624799 | 1.64838349 | 3.19E-06 | 3.87E-05 |
| Q15149-3 | PLEC | 0.0753402 | 0.442139822 | 0.769869141 | -1.00585214 | 8.55E-06 | 0.000121131 | -1.060997941 | 1.27E-05 | 0.000120813 |
| Q15149-4 | PLEC | 3.21E-20 | 1 | 1 | -0.789710402 | 0.000186465 | 0.001444332 | -1.05987127 | 0.000162195 | 0.001020297 |
| P20290;P20290-2 | BTF3 | 0.743713827 | 0.005956711 | 0.024622933 | 0.519528191 | 0.008938059 | 0.033461037 | 1.081539008 | 0.000664543 | 0.003332981 |
| Q9UKD1 | GMEB2 | 0.602438707 | 0.003297444 | 0.015119361 | 0.590667765 | 0.008520723 | 0.03224306 | 0.838938074 | 0.001678281 | 0.007112845 |
| P05114 | HMGN1 | 0 | 1 | 1 | 1.147966455 | 0.001069331 | 0.005961042 | 1.06498665 | 0.003305583 | 0.012356698 |
| Q9UFW8 | CGGBP1 | 0.91940737 | 8.65E-05 | 0.000737014 | 0.837092052 | 0.000304573 | 0.002178626 | 0.68225497 | 0.010366174 | 0.032164795 |
| B9A064;P0DOX8 | IGLL5 | 0 | 1 | 1 | 1.131546539 | 0.008596605 | 0.03248636 | 0.986223773 | 0.01261256 | 0.037681337 |
| Q14011 | CIRBP | 0 | 1 | 1 | 1.008100535 | 0.00050222 | 0.003282573 | 1.037159702 | 0.000153135 | 0.000979911 |
