## Supplementary Table 3 for "Age-associated inflammatory monocytes are increased in menopausal females and reversed by Hormone Replacement Therapy"

| Protein_id | Gene name | log2fold change | p value | qvalue |
|---|---|---|---|---|
| P62995;P62995-3 | TRA2B | -1.100278505 | 1.67E-18 | 5.03E-15 |
| P84103;P84103-2 | SRSF3 | -0.865215356 | 3.22E-17 | 4.86E-14 |
| Q16629;Q16629-2;Q16629-3;Q16629-4 | SRSF7 | -0.720324636 | 1.28E-12 | 1.29E-09 |
| Q99729-2;Q99729-3 | HNRNPAB | -0.626094543 | 3.39E-12 | 2.55E-09 |
| Q01130 | SRSF2 | -0.724361888 | 1.55E-11 | 9.34E-09 |
| P68371 | TUBB4B | -0.706807053 | 1.07E-10 | 4.04E-08 |
| P31943 | HNRNPH1 | -0.494218011 | 9.61E-11 | 4.04E-08 |
| P08621 | SNRNP70 | -0.505421785 | 8.59E-11 | 4.04E-08 |
| Q92882 | OSTF1 | -0.771671616 | 4.56E-10 | 1.25E-07 |
| P06899 | H2BC11 | -1.143044434 | 4.39E-10 | 1.25E-07 |
| Q14103;Q14103-3 | HNRNPD | -0.515548282 | 3.80E-10 | 1.25E-07 |
| P16070;P16070-10;P16070-11;P16070-12;P16070-13;P16070-14;P16070-16;P16070-17;P16070-18;P16070-3;P16070-4;P16070-5;P16070-6;P16070-7;P16070-8 | CD44 | -1.605318862 | 6.73E-10 | 1.69E-07 |
| Q9BTT0-3 | ANP32E | -0.849210569 | 7.42E-10 | 1.72E-07 |
| P37235 | HPCAL1 | -0.795650408 | 1.06E-09 | 2.28E-07 |
| O75340 | PDCD6 | -0.937547218 | 2.48E-09 | 4.67E-07 |
| P18077 | RPL35A | -0.439747371 | 2.42E-09 | 4.67E-07 |
| P61225 | RAP2B | -1.806110088 | 5.45E-09 | 9.12E-07 |
| P17661 | DES | -2.629406468 | 5.17E-09 | 9.12E-07 |
| P19338 | NCL | -0.609576108 | 6.68E-09 | 1.06E-06 |
| P43243 | MATR3 | -0.445319307 | 7.20E-09 | 1.08E-06 |
| P50914 | RPL14 | -0.659315108 | 7.60E-09 | 1.09E-06 |
| P07437 | TUBB | -1.132395098 | 1.03E-08 | 1.32E-06 |
| P68400 | CSNK2A1 | -0.438006873 | 1.05E-08 | 1.32E-06 |
| Q14697 | GANAB | -0.4215744 | 1.65E-08 | 1.99E-06 |
| P54852 | EMP3 | -1.385359913 | 2.15E-08 | 2.42E-06 |
| Q9Y6A9 | SPCS1 | -1.470980547 | 2.17E-08 | 2.42E-06 |
| P36578 | RPL4 | -0.853228865 | 2.32E-08 | 2.49E-06 |
| P83731 | RPL24 | -0.659715706 | 2.61E-08 | 2.71E-06 |
| P23497 | SP100 | -1.011276676 | 2.83E-08 | 2.85E-06 |
| P78527 | PRKDC | -0.504891505 | 3.45E-08 | 3.35E-06 |
| Q16512 | PKN1 | -0.418904333 | 5.14E-08 | 4.84E-06 |
| Q16563-2 | SYPL1 | -1.418122746 | 7.22E-08 | 6.59E-06 |
| P01042;P01042-2;P01042-3 | KNG1 | -2.930471938 | 8.47E-08 | 7.50E-06 |
| O60888;O60888-2;O60888-3 | CUTA | -2.019768966 | 9.26E-08 | 7.97E-06 |
| P61106 | RAB14 | -0.606922987 | 1.52E-07 | 1.27E-05 |
| P62136 | PPP1CA | -0.511231026 | 1.58E-07 | 1.28E-05 |
| P09429 | HMGB1 | -0.662339957 | 2.60E-07 | 2.01E-05 |
| O75915 | ARL6IP5 | -1.112629058 | 3.30E-07 | 2.48E-05 |
| P51970 | NDUFA8 | -1.387048418 | 3.72E-07 | 2.73E-05 |
| P20701;P20701-2 | ITGAL | -0.803513336 | 3.86E-07 | 2.77E-05 |
| P62280 | RPS11 | -0.602525414 | 4.09E-07 | 2.86E-05 |
| P68402 | PAFAH1B2 | -1.393639612 | 4.20E-07 | 2.87E-05 |
| B0I1T2 | MYO1G | -0.715491382 | 5.39E-07 | 3.53E-05 |
| Q9H2H8 | PPIL3 | -0.505347499 | 6.63E-07 | 4.25E-05 |
| P05543 | SERPINA7 | -1.999165173 | 7.27E-07 | 4.49E-05 |
| Q12982;Q12982-2 | BNIP2 | -1.120184521 | 7.45E-07 | 4.49E-05 |

| | | | | |
|---|---|---|---|---|
| Q9UPN7 | PPP6R1 | -0.592266027 | 7.60E-07 | 4.49E-05 |
| P62266 | RPS23 | -0.613613137 | 7.83E-07 | 4.53E-05 |
| O14964 | HGS | 0.469724046 | 8.13E-07 | 4.59E-05 |
| Q92804;Q92804-2 | TAF15 | -0.509856004 | 8.22E-07 | 4.59E-05 |
| P20742 | PZP | -2.073240983 | 1.03E-06 | 5.35E-05 |
| P39687 | ANP32A | -0.919035851 | 9.99E-07 | 5.35E-05 |
| P13473-2 | LAMP2 | -1.162236261 | 1.02E-06 | 5.35E-05 |
| P06748 | NPM1 | -0.59203409 | 1.28E-06 | 6.46E-05 |
| Q9NRW3 | APOBEC3C | -0.966384319 | 1.34E-06 | 6.61E-05 |
| Q86XR7-2;Q9Y3B3;Q9Y3B3-2 | TICAM2;TMED7 | -1.543720516 | 1.36E-06 | 6.61E-05 |
| Q68D06;Q68D06-2 | SLFN13 | -1.950562139 | 1.61E-06 | 7.68E-05 |
| P68363 | TUBA1B | -0.846959537 | 1.93E-06 | 9.09E-05 |
| P32942 | ICAM3 | -0.993455759 | 2.00E-06 | 9.29E-05 |
| P62847;P62847-2;P62847-3;P62847-4 | RPS24 | -0.862893756 | 2.17E-06 | 9.90E-05 |
| P62273 | RPS29 | -1.810396347 | 2.61E-06 | 0.000117305 |
| P02452 | COL1A1 | -1.287479784 | 2.69E-06 | 0.000119174 |
| Q9Y6R0 | NUMBL | -1.199178511 | 2.78E-06 | 0.000121355 |
| P18084 | ITGB5 | -1.187454113 | 2.87E-06 | 0.000123401 |
| Q12800;Q12800-3 | TFCP2 | -0.724683868 | 3.54E-06 | 0.000150398 |
| Q07666 | KHDRBS1 | -0.457226937 | 3.62E-06 | 0.000151658 |
| Q15287;Q15287-2;Q15287-3 | RNPS1 | -0.761825795 | 4.28E-06 | 0.000176776 |
| Q13247;Q13247-3 | SRSF6 | -0.567482458 | 4.76E-06 | 0.000192613 |
| P57772 | EEFSEC | 0.408264666 | 4.79E-06 | 0.000192613 |
| P61960 | UFM1 | -1.415766173 | 5.19E-06 | 0.000205624 |
| P04899 | GNAI2 | -0.76651167 | 5.46E-06 | 0.000213637 |
| P28065 | PSMB9 | -0.495799515 | 5.81E-06 | 0.000224428 |
| O75995 | SASH3 | -0.72307495 | 6.84E-06 | 0.000257605 |
| Q8N257 | H2BU1 | -2.09192943 | 7.19E-06 | 0.000267478 |
| P32969 | RPL9 | -0.443578656 | 7.41E-06 | 0.000272107 |
| P61254 | RPL26 | -0.767630358 | 7.67E-06 | 0.000278543 |
| Q07021 | C1QBP | -0.994248924 | 8.07E-06 | 0.000289418 |
| P01008 | SERPINC1 | -2.075147213 | 8.43E-06 | 0.000298953 |
| P0DP23;P0DP24;P0DP25 | CALM1;CALM2;CALM3 | -1.507381944 | 1.04E-05 | 0.000353423 |
| P23527 | H2BC17 | -2.4078951 | 1.04E-05 | 0.000353423 |
| P08575;P08575-10;P08575-4;P08575-5;P08575-6;P08575-7;P08575-8;P08575-9 | PTPRC | -0.822010993 | 1.04E-05 | 0.000353423 |
| O00160 | MYO1F | -0.499020831 | 1.04E-05 | 0.000353423 |
| Q07955 | SRSF1 | -0.733092915 | 1.08E-05 | 0.000360709 |
| Q7Z7H5;Q7Z7H5-3 | TMED4 | -0.730118753 | 1.09E-05 | 0.000361148 |
| P28066 | PSMA5 | -0.552288719 | 1.37E-05 | 0.000449235 |
| Q8N6M0 | OTUD6B | 0.689897462 | 1.39E-05 | 0.000450854 |
| P52272;P52272-2 | HNRNPM | -0.415824839 | 1.51E-05 | 0.000482863 |
| Q05519;Q05519-2 | SRSF11 | -1.66875139 | 1.61E-05 | 0.000509453 |
| Q13595;Q13595-3;Q13595-4 | TRA2A | -0.53041489 | 1.68E-05 | 0.00052884 |
| P62906 | RPL10A | -0.606182749 | 1.78E-05 | 0.000553686 |
| Q8N386 | LRRC25 | -2.153579892 | 2.04E-05 | 0.000626016 |
| Q15075 | EEA1 | 0.447975313 | 2.14E-05 | 0.000643474 |
| Q8IUE6 | H2AC21 | -1.553137221 | 2.25E-05 | 0.000672674 |
| P02749 | APOH | -1.791442161 | 2.39E-05 | 0.000673317 |
| P61020 | RAB5B | -0.434471406 | 2.39E-05 | 0.000673317 |

| | | | | |
|---|---|---|---|---|
| P14678;P14678-2;P14678- | SNRPB;SNRPN | -0.417116776 | 2.30E-05 | 0.000673317 |
| P02753 | RBP4 | -0.901590938 | 2.39E-05 | 0.000673317 |
| P43652 | AFM | -1.729902327 | 2.45E-05 | 0.000683274 |
| P0C0L4;P0C0L4-2 | C4A | -1.633320165 | 2.52E-05 | 0.000695739 |
| P58876 | H2BC5 | -0.624679822 | 2.76E-05 | 0.000756818 |
| P21912 | SDHB | -0.615575686 | 3.09E-05 | 0.00083047 |
| P62854 | RPS26 | -0.657183576 | 3.21E-05 | 0.000854623 |
| P07339 | CTSD | -0.55034268 | 3.28E-05 | 0.000867626 |
| P02647 | APOA1 | -1.80120452 | 3.64E-05 | 0.000954486 |
| Q9Y6E2 | BZW2 | -0.561455478 | 4.01E-05 | 0.001041588 |
| P62249 | RPS16 | -0.547507286 | 4.48E-05 | 0.001126423 |
| Q92688;Q92688-2 | ANP32B | -0.549701038 | 4.41E-05 | 0.001126423 |
| Q8IVL0;Q8IVL0-2;Q8IVL0-3 | NAV3 | -1.768108822 | 4.49E-05 | 0.001126423 |
| O43314;O43314-2 | PPIP5K2 | 0.74147028 | 4.62E-05 | 0.00114938 |
| P61026 | RAB10 | -0.553341504 | 4.69E-05 | 0.001157719 |
| P16150 | SPN | -1.206427078 | 5.16E-05 | 0.001263833 |
| P62937 | PPIA | -0.426233664 | 5.41E-05 | 0.00131335 |
| O95470 | SGPL1 | -1.196749403 | 5.49E-05 | 0.001322937 |
| P51531;P51531-2 | SMARCA2 | -0.485506574 | 5.72E-05 | 0.001367881 |
| P01023 | A2M | -1.634473678 | 6.50E-05 | 0.001530279 |
| Q13510;Q13510-2 | ASAH1 | -0.506899943 | 6.83E-05 | 0.001594089 |
| Q53RT3;Q53RT3-2 | ASPRV1 | -1.129667274 | 7.23E-05 | 0.0016748 |
| P34810;P34810-2;P34810-3 | CD68 | -1.457028973 | 7.53E-05 | 0.001730839 |
| P51571 | SSR4 | -1.099868944 | 7.91E-05 | 0.001804116 |
| Q01105 | SET | -0.572692467 | 8.33E-05 | 0.001872037 |
| P46778 | RPL21 | -0.57965366 | 8.39E-05 | 0.001872037 |
| Q86YQ8 | CPNE8 | -0.853365074 | 8.78E-05 | 0.001944475 |
| P20340-2 | RAB6A | -0.561634551 | 9.13E-05 | 0.002008191 |
| P43307 | SSR1 | -1.228668913 | 9.39E-05 | 0.002020777 |
| Q9NVS9;Q9NVS9-3 | PNPO | -1.768919107 | 9.31E-05 | 0.002020777 |
| P60953 | CDC42 | -0.435008796 | 9.82E-05 | 0.002099039 |
| Q9Y608;Q9Y608-4 | LRRFIP2 | 0.829781975 | 0.000103153 | 0.002188727 |
| Q9H0X9;Q9H0X9-3 | OSBPL5 | -0.905889119 | 0.000106141 | 0.002236385 |
| P26599;P26599-1;P26599-2 | PTBP1 | -0.460769813 | 0.000107191 | 0.00224283 |
| Q86SX6 | GLRX5 | -1.13897207 | 0.000121225 | 0.002501709 |
| P83876 | TXNL4A | 0.438757268 | 0.000122475 | 0.002510316 |
| P61313 | RPL15 | -0.705819305 | 0.000125643 | 0.002557857 |
| Q13126 | MTAP | -0.52086429 | 0.000137226 | 0.002756415 |
| Q13619 | CUL4A | 0.626045587 | 0.000140469 | 0.00280287 |
| Q15904 | ATP6AP1 | -0.822451612 | 0.000145213 | 0.00285966 |
| Q16864;Q16864-2 | ATP6V1F | -1.450328045 | 0.000146758 | 0.002871317 |
| O95445 | APOM | -1.434441411 | 0.000155361 | 0.002910655 |
| P19823 | ITIH2 | -1.866699155 | 0.000155531 | 0.002910655 |
| P62805 | H4C1 | -1.037376947 | 0.000155329 | 0.002910655 |
| Q9H299 | SH3BGRL3 | -1.63132635 | 0.000150884 | 0.002910655 |
| Q86UP2 | KTN1 | 0.406644914 | 0.000160937 | 0.002956733 |
| P00734 | F2 | -1.313304135 | 0.000160177 | 0.002956733 |
| P40429 | RPL13A | -0.789275413 | 0.000161922 | 0.002956795 |
| Q9UIG0;Q9UIG0-2 | BAZ1B | -0.42363066 | 0.000169446 | 0.003024668 |
| O75489 | NDUFS3 | -0.926426199 | 0.000167473 | 0.003024668 |

| | | | | |
|---|---|---|---|---|
| P08697;P08697-2 | SERPINF2 | -1.630079166 | 0.000169158 | 0.003024668 |
| Q96HY6 | DDRGK1 | -0.663687271 | 0.000169654 | 0.003024668 |
| P26583 | HMGB2 | -0.524376332 | 0.000176811 | 0.003102819 |
| Q9BUF5 | TUBB6 | -1.292934534 | 0.000175216 | 0.003102819 |
| Q9Y5B9 | SUPT16H | -0.435686433 | 0.000182448 | 0.003177542 |
| P35443 | THBS4 | -1.769553659 | 0.000183891 | 0.003184266 |
| P10809 | HSPD1 | -0.499612055 | 0.000190204 | 0.003256162 |
| P13073 | COX4I1 | -1.047306136 | 0.000189386 | 0.003256162 |
| P54709 | ATP1B3 | -0.911124503 | 0.000200328 | 0.003410106 |
| O60216 | RAD21 | -0.465405533 | 0.000202637 | 0.003430031 |
| P0C0S8;Q96KK5;Q99878 | H2AC11;H2AC12;H2AC14 | -1.253277398 | 0.00020963 | 0.003455044 |
| P51114;P51114-2 | FXR1 | -0.573132955 | 0.000208045 | 0.003455044 |
| Q8IXH7;Q8IXH7-4 | NELFCD | -0.955647007 | 0.000210995 | 0.003455044 |
| P01024 | C3 | -1.418723685 | 0.000215408 | 0.003508245 |
| Q13151 | HNRNPA0 | -0.473900911 | 0.00023192 | 0.003756854 |
| P07948 | LYN | -0.537983768 | 0.00024664 | 0.003949255 |
| Q96AG4 | LRRC59 | -0.984224738 | 0.00024773 | 0.003949255 |
| P41227 | NAA10 | -0.420013943 | 0.000258351 | 0.004075449 |
| P67775 | PPP2CA | -0.755610959 | 0.000257532 | 0.004075449 |
| P63173 | RPL38 | -0.976757638 | 0.000264355 | 0.004148446 |
| Q9Y2K7 | KDM2A | -0.799870269 | 0.000268801 | 0.00417473 |
| P07948-2 | LYN | -0.694417004 | 0.000267547 | 0.00417473 |
| Q07954 | LRP1 | -1.050319676 | 0.000284078 | 0.004389366 |
| Q9NRP0 | OSTC | -1.379801148 | 0.000306688 | 0.004714539 |
| P09211 | GSTP1 | -0.644388071 | 0.000312073 | 0.004772969 |
| P19827 | ITIH1 | -1.350319307 | 0.000315862 | 0.004806527 |
| Q8NHV1 | GIMAP7 | -0.614408968 | 0.000317961 | 0.004814148 |
| P62753 | RPS6 | -0.523175878 | 0.000324572 | 0.004889678 |
| Q96AX2 | RAB37 | -0.399434343 | 0.000338369 | 0.005047066 |
| Q12824 | SMARCB1 | 0.763460558 | 0.000356317 | 0.005262663 |
| Q9BSJ8 | ESYT1 | -0.572221945 | 0.000360743 | 0.005285927 |
| Q08945 | SSRP1 | -0.503780964 | 0.000361401 | 0.005285927 |
| P62879 | GNB2 | -0.593256084 | 0.000368286 | 0.005360605 |
| Q96CS3 | FAF2 | -0.734827112 | 0.000372589 | 0.005385135 |
| Q8TAF3 | WDR48 | 0.556779623 | 0.000373546 | 0.005385135 |
| Q9Y241 | HIGD1A | -1.236204056 | 0.000387987 | 0.00551464 |
| O15533;O15533-3 | TAPBP | -1.278844968 | 0.00038802 | 0.00551464 |
| P61019 | RAB2A | -0.666457883 | 0.000395688 | 0.005571067 |
| Q09028;Q09028-3 | RBBP4 | -0.429304405 | 0.00040994 | 0.00574488 |
| Q9BRT3 | MIEN1 | -0.842025939 | 0.000412268 | 0.005750754 |
| Q8N2U0 | TMEM256 | -0.774046411 | 0.000439963 | 0.006080768 |
| Q92506 | HSD17B8 | 0.654281678 | 0.000455361 | 0.006264859 |
| Q96L46 | CAPNS2 | -0.767914716 | 0.000462736 | 0.006308707 |
| P23368 | ME2 | -0.398222213 | 0.000476482 | 0.006466851 |
| Q96T25 | ZIC5 | -0.723692883 | 0.00049053 | 0.006607269 |
| P05976;P05976-2 | MYL1 | -0.75428631 | 0.000491214 | 0.006607269 |
| Q8WZ82 | OVCA2 | 0.495838207 | 0.000495757 | 0.00660936 |
| P12236 | SLC25A6 | -1.447871851 | 0.000504898 | 0.00667444 |
| P09326 | CD48 | -0.685470183 | 0.000505069 | 0.00667444 |
| O14521;O14521-2 | SDHD | -1.212280501 | 0.000510475 | 0.006716418 |

| | | | | |
|---|---|---|---|---|
| P49821;P49821-2 | NDUFV1 | -0.671408848 | 0.000528739 | 0.006896497 |
| O43592 | XPOT | -0.601912871 | 0.000526789 | 0.006896497 |
| P46783 | RPS10 | -0.927207948 | 0.000534467 | 0.006936463 |
| P62310 | LSM3 | -1.332934037 | 0.000538457 | 0.006936463 |
| O14773 | TPP1 | -0.536330797 | 0.000552875 | 0.007058523 |
| P04406 | GAPDH | -0.488995661 | 0.000579913 | 0.007372478 |
| P84090 | ERH | -0.998370626 | 0.000594164 | 0.007521912 |
| P23469;P23469-2;P23469-3 | PTPRE | -0.593151096 | 0.000599448 | 0.007557055 |
| Q9Y547 | HSPB11 | -0.868602952 | 0.000604739 | 0.007592 |
| P18124 | RPL7 | -0.521936749 | 0.000627122 | 0.007780311 |
| Q06033;Q06033-2 | ITIH3 | -1.163384082 | 0.000630721 | 0.007780311 |
| Q9Y4Z0 | LSM4 | -0.775918534 | 0.000624538 | 0.007780311 |
| P67812 | SEC11A | -1.21466899 | 0.000646581 | 0.007887235 |
| P40616;P40616-2 | ARL1 | -0.651119774 | 0.000652147 | 0.00792306 |
| P04080 | CSTB | -0.913651978 | 0.000662029 | 0.008010811 |
| P17096-2 | HMGA1 | -1.032402199 | 0.000685004 | 0.00822278 |
| Q9NVZ3 | NECAP2 | -0.486213389 | 0.000689775 | 0.008247187 |
| P31749 | AKT1 | -0.55057971 | 0.000715962 | 0.008526462 |
| P61970 | NUTF2 | -1.026854592 | 0.000776539 | 0.009175345 |
| Q14031;Q14031-2 | COL4A6 | -1.252521414 | 0.00078894 | 0.009285451 |
| Q13283 | G3BP1 | -0.540937805 | 0.00083571 | 0.009610669 |
| P45985;P45985-2 | MAP2K4 | 0.599590921 | 0.000843975 | 0.009668809 |
| Q99447-4 | PCYT2 | -0.803921533 | 0.000851447 | 0.009680795 |
| Q03181;Q03181-2;Q03181-3;Q03181-4 | PPARD | -1.313441757 | 0.000866444 | 0.009777507 |
| P35244 | RPA3 | -0.855942463 | 0.000883774 | 0.009862262 |
| P11234;P11234-2 | RALB | -0.417158411 | 0.000883747 | 0.009862262 |
| Q9BY42 | RTF2 | 0.674241463 | 0.000903389 | 0.010043956 |
| P17655 | CAPN2 | -0.402607025 | 0.000973034 | 0.010660917 |
| Q03518;Q03518-2 | TAP1 | -1.255019603 | 0.000971704 | 0.010660917 |
| Q9BTV4 | TMEM43 | -0.963091513 | 0.000971815 | 0.010660917 |
| Q15005 | SPCS2 | -0.896770804 | 0.000967234 | 0.010660917 |
| A8MWD9;P62308 | SNRPGP15;SNRPG | -1.058418203 | 0.001036739 | 0.011276879 |
| Q9Y251;Q9Y251-2 | HPSE | 0.708159112 | 0.001045303 | 0.011329132 |
| P15880 | RPS2 | -0.44257068 | 0.001067325 | 0.011515004 |
| P62979 | RPS27A | -0.525391939 | 0.001070097 | 0.011515004 |
| Q9NVG8 | TBC1D13 | 0.409573153 | 0.001081096 | 0.011591969 |
| Q9UHA4 | LAMTOR3 | -0.992699334 | 0.001121853 | 0.011986318 |
| P68133 | ACTA1 | -1.402583641 | 0.001126721 | 0.011995795 |
| P37840;P37840-2 | SNCA | 1.115455986 | 0.001136698 | 0.012059407 |
| P48960 | ADGRE5 | -0.746173919 | 0.001153533 | 0.012195074 |
| Q9H3N1 | TMX1 | -1.090325405 | 0.001183776 | 0.012471034 |
| O43676 | NDUFB3 | -0.904086458 | 0.001211488 | 0.012718517 |
| Q15018 | ABRAXAS2 | 0.50764136 | 0.00122428 | 0.012719851 |
| P62277 | RPS13 | -0.552959091 | 0.001222575 | 0.012719851 |
| P02771 | AFP | -1.535012252 | 0.00121947 | 0.012719851 |
| P49755 | TMED10 | -0.866138511 | 0.001256827 | 0.013013129 |
| Q92522 | H1-10 | -0.738459926 | 0.001292925 | 0.013250279 |
| P35237 | SERPINB6 | -0.403442287 | 0.001286785 | 0.013250279 |
| A0A024RBG1;Q9NZJ9-2 | NUDT4B;NUDT4 | 0.730550148 | 0.00132993 | 0.013446574 |
| P21796 | VDAC1 | -1.052923583 | 0.001345896 | 0.013562489 |

| | | | | |
|---|---|---|---|---|
| Q15746;Q15746-11;Q15746-2;Q15746-6;Q15746-7;Q15746-9 | MYLK | -0.668487801 | 0.001372837 | 0.013787859 |
| P38159 | RBMX | -0.470196804 | 0.001390475 | 0.013904943 |
| P35754 | GLRX | -0.768058013 | 0.001396007 | 0.013904943 |
| O15260;O15260-2 | SURF4 | -0.967005078 | 0.00139834 | 0.013904943 |
| O75695 | RP2 | -0.76652013 | 0.001410953 | 0.01398421 |
| Q15125 | EBP | -0.861496917 | 0.001466381 | 0.014438582 |
| Q9Y3C8 | UFC1 | -0.396105393 | 0.001500698 | 0.014728348 |
| P07108 | DBI | -0.87152395 | 0.001536665 | 0.01503238 |
| P00491 | PNP | -0.42297187 | 0.001548636 | 0.015100455 |
| P20702 | ITGAX | -0.708633249 | 0.001573235 | 0.015290832 |
| Q07020 | RPL18 | -0.469504673 | 0.001667471 | 0.01592604 |
| O94851;O94851-1;O94851-3 | MICAL2 | -1.233771068 | 0.001668853 | 0.01592604 |
| Q9BPW8 | NIPSNAP1 | -0.512374185 | 0.001667496 | 0.01592604 |
| Q99879 | H2BC14 | -1.180557638 | 0.001670305 | 0.01592604 |
| P62263 | RPS14 | -0.452239564 | 0.001697834 | 0.016137459 |
| Q9UHG3 | PCYOX1 | -0.493745795 | 0.001723237 | 0.016327398 |
| O75251;O75251-2 | NDUFS7 | -0.730396029 | 0.001786494 | 0.016873684 |
| Q02790 | FKBP4 | 0.457643441 | 0.001793345 | 0.016885462 |
| Q8TDH9 | BLOC1S5 | 0.48459762 | 0.001804954 | 0.016941831 |
| P62736 | ACTA2 | -0.516915543 | 0.001899795 | 0.01772162 |
| Q13724;Q13724-2 | MOGS | -0.763007984 | 0.001915453 | 0.017768489 |
| P46781 | RPS9 | -0.458185759 | 0.001916614 | 0.017768489 |
| Q96JC1;Q96JC1-2 | VPS39 | -0.487369571 | 0.001925964 | 0.017800396 |
| Q9Y6C9 | MTCH2 | -0.867615977 | 0.001974864 | 0.018147672 |
| P12109 | COL6A1 | -1.198821208 | 0.001996521 | 0.018228842 |
| P08574 | CYC1 | -0.984518429 | 0.002002605 | 0.018229152 |
| P08236;P08236-2 | GUSB | -0.401906237 | 0.002054497 | 0.018589187 |
| P05107 | ITGB2 | -0.851494185 | 0.002064494 | 0.018623712 |
| P12235 | SLC25A4 | -1.214617165 | 0.002088198 | 0.018763936 |
| Q14624;Q14624-2;Q14624-3;Q14624-4 | ITIH4 | -1.265772275 | 0.002092493 | 0.018763936 |
| Q9H078-2 | CLPB | 0.483669429 | 0.002130307 | 0.019046338 |
| P62081 | RPS7 | -0.451257226 | 0.002152998 | 0.019192258 |
| Q6DD88 | ATL3 | -0.615984297 | 0.002190599 | 0.019415686 |
| P36542 | ATP5F1C | -0.510505098 | 0.00219095 | 0.019415686 |
| P42126 | ECI1 | -0.52716493 | 0.002269559 | 0.020053317 |
| Q6GTX8;Q6GTX8-2;Q6GTX8-3;Q6GTX8-4 | LAIR1 | -0.600869807 | 0.002279712 | 0.020084128 |
| P04439 | HLA-A | -0.677712873 | 0.002292252 | 0.020135732 |
| P19623 | SRM | -0.555789782 | 0.002349043 | 0.020514973 |
| Q9NWY4 | HPF1 | 0.591875024 | 0.002348584 | 0.020514973 |
| P58546 | MTPN | -0.926955637 | 0.002428071 | 0.021143871 |
| Q13242 | SRSF9 | -0.455346902 | 0.002473438 | 0.021415143 |
| O95861 | BPNT1 | 0.409344598 | 0.002492764 | 0.021520626 |
| Q8N6H7 | ARFGAP2 | 0.587767496 | 0.002542213 | 0.021884823 |
| P39656 | DDOST | -1.013633583 | 0.002559933 | 0.021912155 |
| P51159 | RAB27A | -0.672721429 | 0.002601248 | 0.022150317 |
| P36873;P36873-2 | PPP1CC | -0.476250071 | 0.002762502 | 0.023314898 |
| P63272 | SUPT4H1 | 0.647953782 | 0.002774392 | 0.023349839 |
| P60903 | S100A10 | -0.66454975 | 0.002790465 | 0.02335464 |
| Q13409-3 | DYNC1I2 | 0.423255657 | 0.002872671 | 0.023803952 |

| | | | | |
|---|---|---|---|---|
| Q9HDC9 | APMAP | -1.000241974 | 0.002875751 | 0.023803952 |
| P14618-2 | PKM | -0.482353017 | 0.002855446 | 0.023803952 |
| P27824;P27824-2 | CANX | -0.748953786 | 0.002917336 | 0.02408201 |
| P30273 | FCER1G | -1.259553647 | 0.002934209 | 0.024155115 |
| P05141 | SLC25A5 | -1.141562068 | 0.003009674 | 0.024708853 |
| Q16698;Q16698-2 | DECR1 | -0.428857845 | 0.003118642 | 0.025327408 |
| P46776 | RPL27A | -0.45543991 | 0.003114302 | 0.025327408 |
| P53680 | AP2S1 | -0.681715153 | 0.003139501 | 0.025349726 |
| Q96GG9 | DCUN1D1 | 0.429429698 | 0.003129943 | 0.025349726 |
| Q7RTS7 | KRT74 | -1.106452925 | 0.00314663 | 0.025349726 |
| P00387;P00387-2;P00387-3 | CYB5R3 | -0.676773923 | 0.003181436 | 0.025531692 |
| Q12907 | LMAN2 | -0.642515167 | 0.00330301 | 0.026120652 |
| P27482 | CALML3 | -0.91324683 | 0.003316426 | 0.026158095 |
| P62241 | RPS8 | -0.469779519 | 0.003362934 | 0.026455666 |
| P80217;P80217-2 | IFI35 | -0.667663324 | 0.003475651 | 0.027187801 |
| Q9H8S9 | MOB1A | -0.733841519 | 0.00348572 | 0.027187801 |
| P57088 | TMEM33 | -1.049534463 | 0.003492094 | 0.027187801 |
| Q96RW7;Q96RW7-2 | HMCN1 | -1.137884726 | 0.003524717 | 0.027300698 |
| Q8TEA8 | DTD1 | 0.467326267 | 0.003584931 | 0.027695888 |
| O15258 | RER1 | -0.802552877 | 0.003598885 | 0.027732584 |
| Q9Y6M9 | NDUFB9 | -0.925637691 | 0.00365969 | 0.028057621 |
| P26885 | FKBP2 | -0.408682367 | 0.003708448 | 0.028216047 |
| Q8NBJ7 | SUMF2 | -0.553935426 | 0.003771051 | 0.02862009 |
| P35232 | PHB | -0.967073567 | 0.003854964 | 0.02918343 |
| P46779;P46779-2;P46779-3 | RPL28 | -0.455270703 | 0.003916853 | 0.029503693 |
| P56556 | NDUFA6 | -0.898258851 | 0.003916604 | 0.029503693 |
| P14406 | COX7A2 | -1.273216556 | 0.00418887 | 0.031009992 |
| Q02978 | SLC25A11 | -1.063221026 | 0.004245122 | 0.031349397 |
| Q9H8H3 | METTL7A | -0.834816087 | 0.004275654 | 0.03149767 |
| P10515 | DLAT | -0.553266971 | 0.004315998 | 0.031717321 |
| P41223 | BUD31 | 0.578728166 | 0.004374123 | 0.031910976 |
| P07951 | TPM2 | -1.035692649 | 0.004471808 | 0.032388363 |
| O75582 | RPS6KA5 | 0.465004359 | 0.004547474 | 0.032778803 |
| P35613;P35613-2 | BSG | -0.666091084 | 0.004567882 | 0.032847322 |
| Q13155 | AIMP2 | 0.399681891 | 0.004630024 | 0.033136012 |
| Q9BQ61 | TRIR | -0.461051509 | 0.004736169 | 0.033815351 |
| O14828;O14828-2 | SCAMP3 | -0.736918731 | 0.00480646 | 0.034236088 |
| P63261 | ACTG1 | -1.632108331 | 0.004847482 | 0.034446845 |
| Q9NZ08 | ERAP1 | -0.459428147 | 0.004871177 | 0.034452712 |
| Q8NBM8 | PCYOX1L | -0.457138428 | 0.00487071 | 0.034452712 |
| Q92542;Q92542-2 | NCSTN | -0.669361895 | 0.004882684 | 0.034453225 |
| O00505 | KPNA3 | -0.601188746 | 0.004901386 | 0.034472898 |
| P45880 | VDAC2 | -0.945311571 | 0.004908355 | 0.034472898 |
| O75494;O75494-2;O75494-3;O75494-4;O75494-5;O75494-6 | SRSF10 | -0.46927164 | 0.005036759 | 0.035292454 |
| P34896;P34896-2 | SHMT1 | 0.401645799 | 0.005115354 | 0.035719916 |
| Q9UBQ5 | EIF3K | -0.406590501 | 0.005355408 | 0.037008814 |
| Q9UH99 | SUN2 | -0.751200404 | 0.005355189 | 0.037008814 |
| P13612 | ITGA4 | -0.567692488 | 0.005387077 | 0.037142476 |
| P28331;P28331-2 | NDUFS1 | -0.515377819 | 0.005482767 | 0.037715928 |

| | | | | |
|---|---|---|---|---|
| Q96GD0 | PDXP | 0.427116627 | 0.005572342 | 0.0382448 |
| Q562R1 | ACTBL2 | -0.681960914 | 0.005746983 | 0.039264537 |
| Q9NRV9 | HEBP1 | 0.571108576 | 0.005926326 | 0.040125886 |
| Q5EBM0 | CMPK2 | 0.506263562 | 0.006026583 | 0.040713214 |
| P80303;P80303-2 | NUCB2 | -0.431154269 | 0.006109529 | 0.041089309 |
| Q9Y315 | DERA | -0.462064005 | 0.006189505 | 0.041534474 |
| Q8N131;Q8N131-2 | TMEM123 | -0.652585851 | 0.006338709 | 0.042347077 |
| P09493-5 | TPM1 | 0.92159521 | 0.006515725 | 0.04305237 |
| Q8TCT9;Q8TCT9-5 | HM13 | -0.715997356 | 0.00653975 | 0.043116555 |
| P00846 | MT-ATP6 | -0.921142235 | 0.006555836 | 0.043128237 |
| P21964;P21964-2 | COMT | -0.611902756 | 0.006651545 | 0.043567619 |
| O60762 | DPM1 | -0.620659299 | 0.006715291 | 0.043794744 |
| O75964 | ATP5MG | -1.228566311 | 0.006708656 | 0.043794744 |
| Q96BM9 | ARL8A | -0.540232983 | 0.006972467 | 0.045373743 |
| P50213 | IDH3A | -0.448357504 | 0.007137115 | 0.046245437 |
| O00483 | NDUFA4 | -1.298937724 | 0.007154759 | 0.046260277 |
| P48059 | LIMS1 | 0.5477221 | 0.007205997 | 0.046491798 |
| Q13867 | BLMH | -0.414831679 | 0.0072938 | 0.046880663 |
| Q8WVM8 | SCFD1 | -0.417290281 | 0.007297388 | 0.046880663 |
| P09110 | ACAA1 | -0.580101061 | 0.007342577 | 0.046970667 |
| Q8NBX0 | SCCPDH | -0.849976052 | 0.007331023 | 0.046970667 |
| Q99623 | PHB2 | -0.959586447 | 0.007453043 | 0.047475728 |
| B9A064;P0DOX8 | IGLL5 | -0.966091454 | 0.007476057 | 0.047521855 |
| Q9P0L0 | VAPA | -0.440299588 | 0.00761221 | 0.048184013 |
| P60520 | GABARAPL2 | 0.476878199 | 0.007805951 | 0.049306774 |
| Q9Y277 | VDAC3 | -0.966108043 | 0.007834781 | 0.049385346 |
| Q14498;Q14498-2;Q14498-3 | RBM39 | -0.399819433 | 0.007904958 | 0.049628003 |
